# A time for every purpose: using time-dependent sensitivity analysis to help understand and manage dynamic ecological systems

**DOI:** 10.1101/2023.04.13.536769

**Authors:** Wee Hao Ng, Christopher R. Myers, Scott McArt, Stephen P. Ellner

**Affiliations:** Cornell University, Ithaca, New York, 14853

**Keywords:** Sensitivity analysis, time-dependent sensitivity, spillover, optimal control, management, dynamic models.

## Abstract

Sensitivity analysis is often used to help understand and manage ecological systems, by assessing how a constant change in vital rates or other model parameters might affect the management outcome. This allows the manager to identify the most favorable course of action. However, realistic changes are often localized in time—for example, a short period of culling leads to a temporary increase in the mortality rate over the period. Hence, knowing when to act may be just as important as knowing what to act upon. In this article, we introduce the method of time-dependent sensitivity analysis (TDSA) that simultaneously addresses both questions. We illustrate TDSA using three case studies: transient dynamics in static disease transmission networks, disease dynamics in a reservoir species with seasonal life-history events, and endogenously-driven population cycles in herbivorous invertebrate forest pests. We demonstrate how TDSA often provides useful biological insights, which are understandable on hindsight but would not have been easily discovered without the help of TDSA. However, as a caution, we also show how TDSA can produce results that mainly reflect uncertain modeling choices and are therefore potentially misleading. We provide guidelines to help users maximize the utility of TDSA while avoiding pitfalls.

## Introduction

*It is not an overstatement to say that no model is ever fully understood if it does not include a sensitivity analysis*. (Caswell, 2019, p. 4)

Sensitivity analysis is used to help us understand the past, to predict and manage the future, and to identify the key processes in complex systems with multiple feedbacks. The many varieties of sensitivity analysis differ in their mechanics, but all involve making some changes to a model and observing how its projections change. To help us understand the past, a retrospective sensitivity analysis asks how observed past variation in each parameter contributed to relevant features of observed past system dynamics. Life Table Response Experiment analysis in population ecology (e.g., Caswell, 1989, 1996; Hernández et al., 2022; Oli et al., 2001; Oro and Doak, 2020) is perhaps the most familiar example, decomposing the variation (across time or space) in the dominant eigenvalue of a population projection matrix into contributions from variation in each matrix element or demographic parameter. To help us predict and manage the future, prospective sensitivity analyses ask how changes corresponding to potential policy changes or management interventions affect projected outcomes, seeking to find targets of opportunity where relatively small (and hopefully inexpensive) interventions have a large impact on outcomes of interest (e.g., Caswell, 2000; Morris and Doak, 2002). Sensitivity analysis of complex models (e.g., Saltelli et al., 2008) helps us identify which parameters need to be estimated accurately (and which do not) to reliably project properties of interest, and which processes or assumptions are most tightly linked to which features of model projections. It is rare to find a paper that includes a mechanistic model of a biological system but does not include at least one figure showing how solution trajectories, or steady-state model properties, change as some parameters are varied—a sensitivity analysis, or the start of one. Prospective analyses typically involve time-invariant perturbations (e.g., elasticity analysis of matrix projection models (Caswell, 2001)). But in many cases, *when to act* may be just as important as *how to act*. The importance of “when” was impressed on us by our studies of bee parasites transmitted at flowers in eastern U.S. old-field communities (Graystock et al. (2020); Fig. 1(A)). An infected bee defecating on a flower may deposit parasites that can infect other bees visiting the flower subsequently (Burnham et al. (2021); Figueroa et al. (2019); Graystock et al. (2020)). This allows between-species transmission of multi-host parasites, including possible spillover from managed or non-native bees to wild native bees (Arbetman et al., 2013; Fü rst et al., 2014; Graystock et al., 2016, 2013; Manley et al., 2019). Early in the season, our data suggest that the trypanosome parasite *C. bombi* is most prevalent in *Ceratina* and possibly other bee genera, some of which visit flowers that are also visited by bumble bees (*Bombus*), including rare species of conservation concern (Cameron et al., 2011). Later in the season, the parasite is most prevalent in common species of *Bombus* such as *B. impatiens*, some of whom again share floral resources with other native bee species of conservation concern (Bartomeus et al., 2013). As a consequence, an intervention to protect species of concern—for example, by reducing spillover from common *Bombus* species—is likely to be far more effective at some times than others.

**Figure 1:**
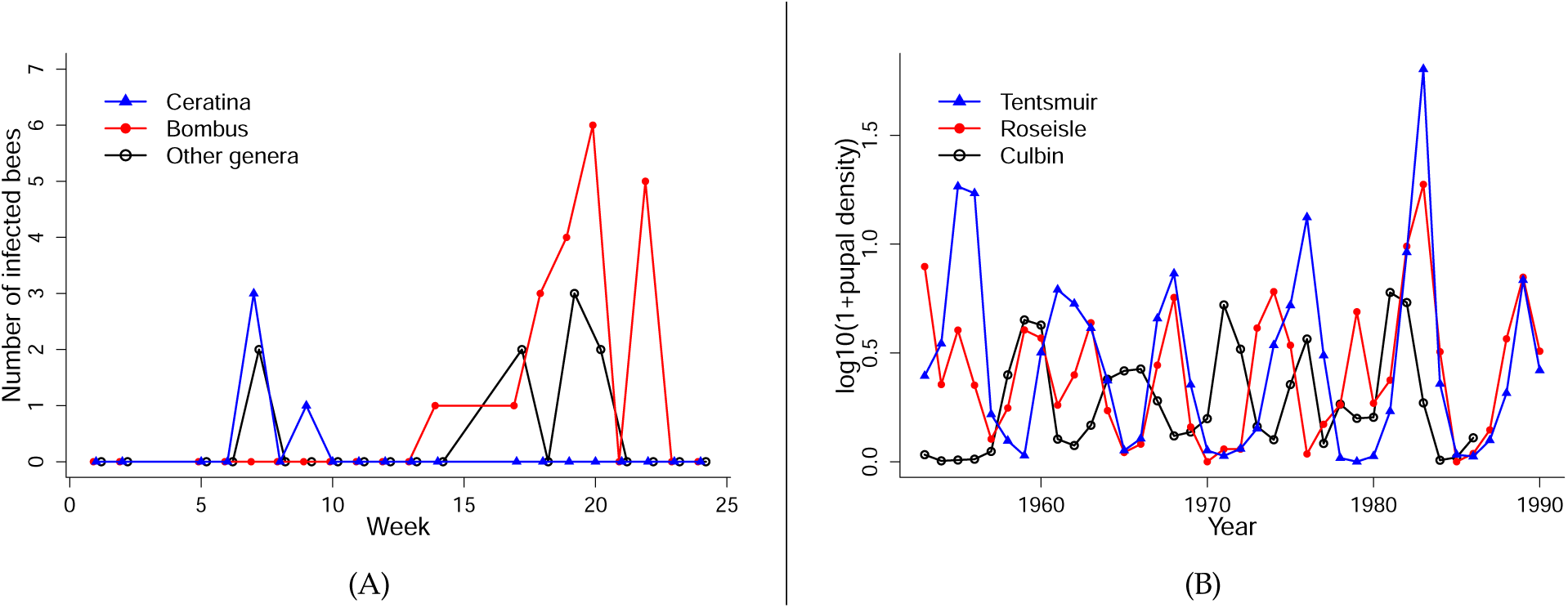
Examples of systems with strong temporal dynamics, where the timing of management interventions might be important. **(A)** Numbers of bees infected with *Crithidia bombi* captured at an old-field site in Lansing, NY, sampled in 2017. Data replotted from Graystock et al. (2020), Supplementary Fig. 2d. **(B)** Oscillations in the abundance of pine looper moth *Bupalus piniarius* in three forests in the UK. Data are log_10_(1+*x*)-transformed annual estimates of spatially averaged pupal abundance by the UK Forestry Commission, from Kendall et al. (2005).

Seasonal turnover in species, likely implying time-varying interaction strengths and therefore time-varying sensitivities, is a common feature of natural and managed systems (e.g., freshwater plankton: Sommer et al. (2012); mycorrhizal fungi: Dumbrell et al. (2011); plant-pollinator communities: CaraDonna and Waser (2020); pests in agroecosystems: Nelson et al. (2013)). Hence, timing is important if humans seek to manage these systems optimally. For example, multivoltine agricultural insect pests may overwinter as inactive eggs or pupae, and then have several semi-discrete generations during the growing season with large changes in the abundance of crop-damaging life stages (e.g., Nelson et al., 2013). On longer time scales, forest insect pests are notorious for having occasional eruptions causing extensive damage, followed by a population crash (e.g. Berryman (1986, Ch. 4), Turchin et al. (2003), Kendall et al. (2005), Myers and Cory (2013)). Dynamics of this sort are illustrated in Fig. 1(B). In such cases, is it better to nip in the bud a growing generation of a multivoltine species or a growing pest outbreak, or to wait until the next peak when an intervention might claim more victims among the pests for the same cost?

Time-dependent sensitivity analysis (TDSA) to address such questions can be done in principle by brute-force computation: simulate the impacts of brief changes to each parameter, and to each state variable, at a fine grid of time points. That may or may not be feasible, depending on model complexity and on how much computing power and time are available. Our goal in this paper is to explain and illustrate a very general and straightforward method for efficiently performing TDSA, called adjoint sensitivity analysis (ASA).

ASA is not new (see for example Cacuci et al. (2003, 1980); Cao et al. (2002, 2003); Errico (1997)), but its biological applications have been very restricted. In some areas of computational science including meteorology, oceanography and earth systems modeling, it is often used in data assimilation, as a numerical method for efficiently computing the derivatives of a likelihood function or other measure of model-data fit, with respect to time-invariant changes in model parameters (e.g., Fröhlich et al., 2017; Lyu et al., 2018; Moore, 2011). But otherwise, it has seen little use in the ecological or epidemiological literature—we did not unearth even one example in our literature search. Here, we apply it to a very different type of question: how and when should we perturb a system to have maximum impact on a biologically-motivated objective function? For instance, we might want to minimize the spread of a disease to a species of concern, or to minimize the damage to a crop plant by an invertebrate pest. Besides the obvious management relevance, we show later that such questions are also interesting theoretically because the answers may provide insights into the dynamics of the system. In addition, we make connections between this approach and optimal control theory (Bressan and Piccoli, 2007; Lenhart and Workman, 2007), which are largely missing from existing literature.

The structure of this paper is as follows. First, we present the mathematical formalism used to perform TDSA, both for deterministic continuous-time and discrete-time models. We then illustrate TDSA using three case studies. The first is a continuous-time disease transmission in hypothetical multispecies networks. These are meant to showcase how TDSA can reveal changes in sensitivities resulting from system dynamics, even when all parameters and dynamic equations are time-invariant. The second and third case studies are empirical examples meant to demonstrate the variety of empirically-fitted models where TDSA can be used to guide the management of real systems. The second is an integral projection model with seasonal dynamics that describes disease maintenance in a reservoir species, while the third involves two discrete-time models of invertebrate pest species that exhibit population cycles, one single-patch and the other spatially explicit. We also use specific instances in the second and third examples to illustrate some potential pitfalls when performing TDSA, and we suggest best practices that can help the practitioner avoid these pitfalls; this is especially important if the results are meant to inform management actions. An R (R Core Team, 2021) package implementing the methods presented here is in development, and will be described in detail elsewhere.

### Calculating time-dependent sensitivities

The steps involved in TDSA are remarkably similar for continuous- and discrete-time models. We therefore give a detailed explanation for continuous time, followed by a brief explanation for discrete time.

#### Continuous-time models

We consider models that can be written as a time-dependent, finite-dimensional system of ordinary differential equations (ODE)

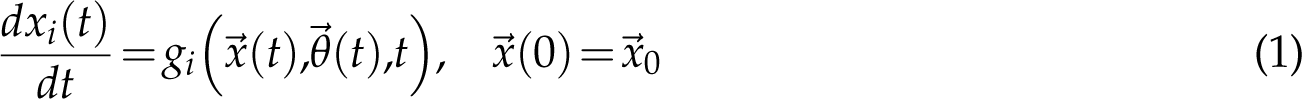

where *⃗x*(*t*) =(*x*_1_(*t*),*x*_2_(*t*),*···*,*x_d_*(*t*))^T^ is the *d*-dimensional state vector, and ^⃗^*θ*(*t*) is a vector of (possibly time-dependent) parameters. (Note that this excludes models that involve integro-differential or delay differential equations, but numerical methods for solving such models, e.g. the linear chain trick (MacDonald, 1978), often involve approximating them by a larger ODE system where Eqn. (1) does apply.) For notational simplicity, we will usually drop the argument ^⃗^*θ*(*t*). We assume that the management goal can be represented by a *reward function J*,

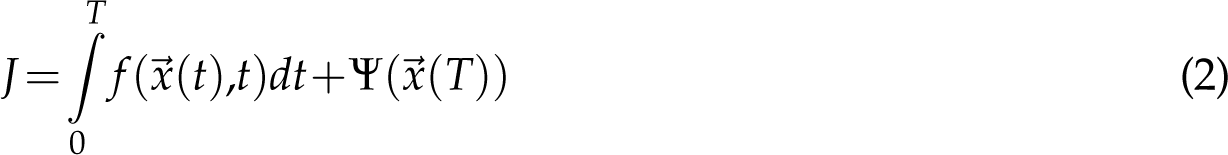

that is to be maximized. *T* is called the time horizon and demarcates the period of interest, *f* represents rewards that accumulate over this time period (hence the integral), while Ψ represents a *terminal payoff* at the end of the period.

As a simple example, consider an organism in a sink habitat, where the per-capita loss rate *µ* (mortality and emigration combined) exceeds the per-capita unregulated birth rate *b*, so the population is only maintained through immigration at a rate *σ*. However, due to ongoing habitat restoration efforts, *µ* begins to decrease over time, so the population should eventually become self-sustaining (see Fig. 2(A)). The dynamics is given by

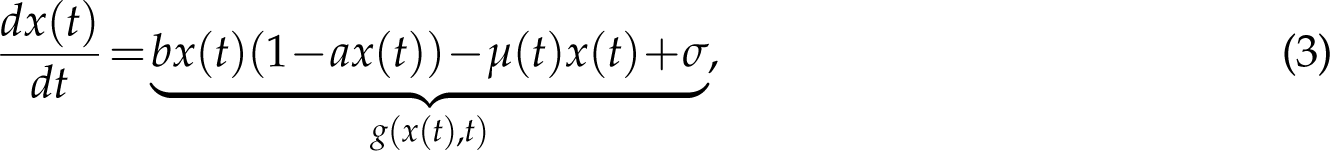

**Figure 2:**
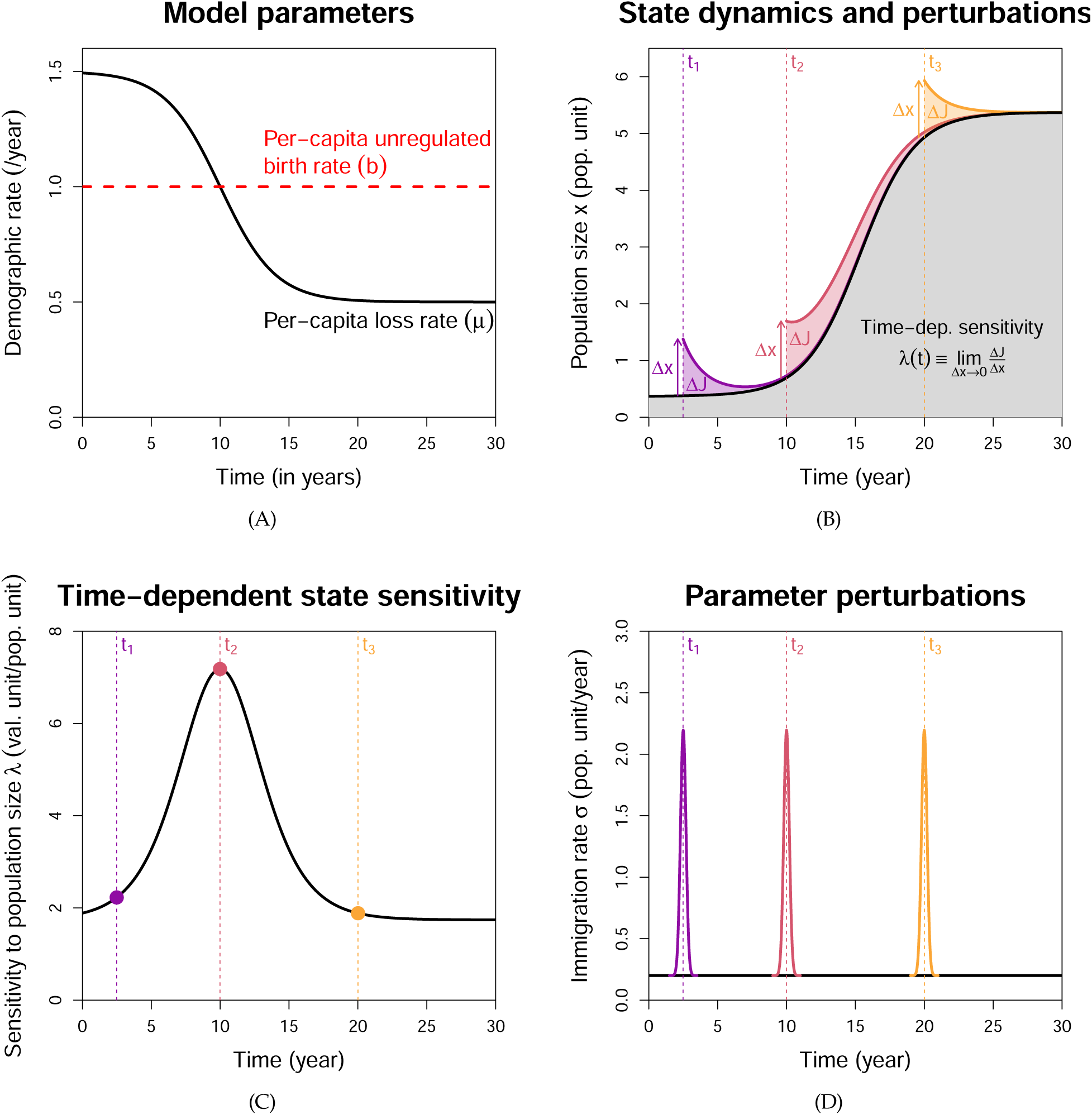
Illustration of time-dependent sensitivity analysis. **(A)** We consider an organism in a sink habitat that is being improved through restoration efforts, so the per-capita loss rate *µ*(*t*) will eventually fall below the per-capita unregulated birth date *b*. **(B)** The population trajectory *x*(*t*) is shown in black, and we assume the reward function *J* is the grey area under the trajectory, plus a terminal payoff (not shown). Now consider a one-off translocation effort to speed up the population recovery at time *t*. This corresponds to a perturbation *x*(*t*) *→x*(*t*)+Δ*x*, and leads to a change Δ*J* in the reward. Δ*J* can depend on the translocation time *t*; for example, it is larger at *t*_2_ than at *t*_1_ or *t*_3_. **(C)** Not surprisingly, the state sensitivity *λ*(*t*) is also higher at time *t*_2_. Hence, translocation is most effective right around when *µ*(*t*) = *b*, so the population has just become self-sustaining. **(D)** Time-dependent parameter sensitivities can be calculated from the state sensitivities. A brief spike in the immigration rate parameter *σ* at time *t* produces a state perturbation at time *t*, and the resulting change in *J*can be inferred from *λ*(*t*). Generalizing this to arbitrary parameter perturbations is straightforward, see Eqn. (A9).

where *x*(*t*) is the population size at time *t*, and *a* the coefficient for reproductive competition. At the same time, the organism provides an important ecosystem service, so over a management period from *t* = 0 to *T*, one can define the reward function

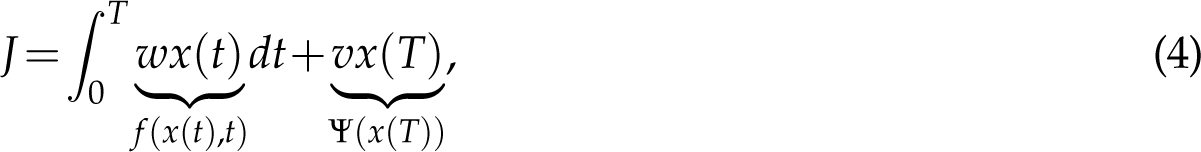

where the first term represents the total value of the service over the period (so *w* is the per-capita rate of contribution), and the second term is a terminal payoff that ascribes value to having a large population at the end of the period (so *v* is the value per individual). See Online Supplement Sec. S1 for parameter values.

TDSA addresses the question of how the value of the reward *J* changes in response to a small, sudden perturbation of a state variable at some time *t*, after which the state is then allowed to continue along its dynamic trajectory starting from the modified value. Returning to our example, we may want to translocate individuals to the habitat to speed up the recovery of the population and increase the reward *J*. A one-off translocation would cause a small, sudden increase in the population size as shown in Fig. 2(B). Formally, we define the sensitivity to state variable *x_i_* at time 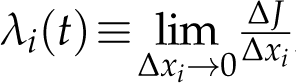, where Δ*J* is the change in *J* resulting from a sudden perturbation *x_i_*(*t*) *→x_i_*(*t*)+Δ*x_i_*. Hence the change in reward Δ*J* is approximately *λ_i_*(*t*)Δ*x_i_* when the perturbation Δ*x_i_* is small. The sensitivities will depend on the time of perturbation *t*, and so can tell us when certain management actions, such as a translocation, would have the most effect on *J*.

Sensitivities can be calculated directly from their definition (perturb a state variable, recalculate the state trajectory, and determine the change in *J*), but it is computationally much more efficient to use the adjoint method. The state sensitivities *λ_i_*(*t*) themselves satisfy an ODE system called the adjoint

equations,

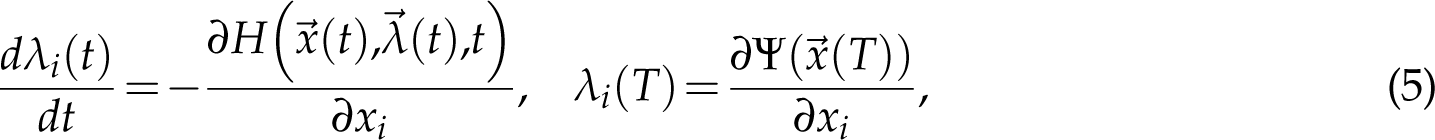

Where

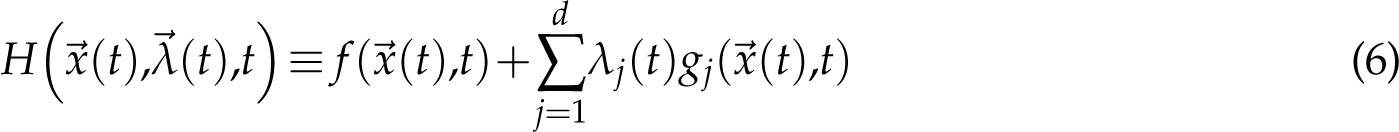

is called the Hamiltonian; see Appendix A for the derivation. For the purpose of this article *H* can be regarded as a construct that simplifies the expression in Eqn. (5), but an in-depth explanation can be found in Dixit (1990), Chapter 10. Because the terminal conditions *λ_i_*(*T*) are known, the adjoint equations are solved *backward* in time from *t* = *T* to *t* = 0, giving the sensitivity values at all times 0 *≤ t ≤ T*. In the context of this method, the state sensitivities are called adjoint variables, and there is one adjoint variable *λ_i_* for each state variable *x_i_*.

For our example, from Eqns. (3) and (4), we can write down the Hamiltonian

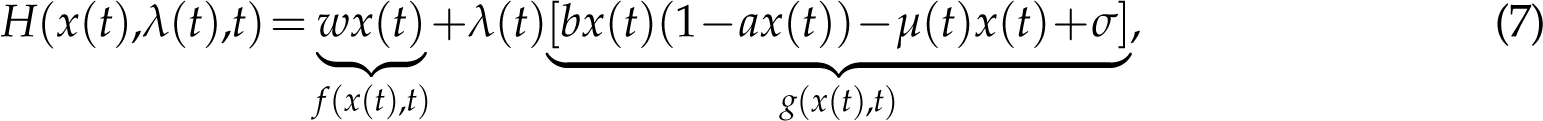

Differentiating *H*(*x*(*t*),*λ*(*t*),*t*) and Ψ(*x*(*T*)) (from Eqn. (4)) in *x*, we obtain the adjoint equation and terminal condition

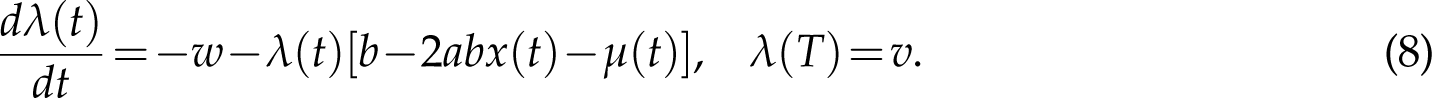

Once we have solved Eqn. (3) for the state trajectory *x*(*t*) (black curve in Fig. 2(B)), the right side of the adjoint equation is fully specified (except for *λ*(*t*)). We can then solve the adjoint equation backward in time to obtain *λ*(*t*) at all *t* (Fig. 2(C)). We see that translocation is most effective roughly when *µ*(*t*) has decreased below *b* so the population has become self-sustaining, an intuitive result. Translocate too early, and few translocated individuals will survive long due to the still-high *µ*(*t*). Translocate too late and the population has already recovered back to its carrying capacity, so even though translocated individuals survive longer due to the low *µ*(*t*), they will also suppress the per-capita birth rate *b*(1*−ax*(*t*)) below *µ*(*t*).

We came upon the idea of using the adjoint method for time-dependent sensitivity calculations through optimal control theory (OCT). Conceptually, OCT also involves a system/reward combination like Eqns. (1–2), except that the functions *g* and *f* now depend on an additional variable *u*(*t*) that quantifies external manipulation (control) of the system, so *g* describes the manipulated dynamics while *f* incorporates the cost of implementing the control. Adjoint variables first show up when we apply Pontryagin’s maximum principle (Pontryagin et al., 1962) to find the optimal control strategy *u^∗^*(*t*) that maximises *J*. More importantly, it is known that the adjoint variable *λ_i_*(*t*) can be interpreted as a “shadow price” (Lenhart and Workman (2007), Section 2.2), the additional profit associated with an increment of *x_i_* at time *t*. For an unmanipulated system, this is equivalent to the time-dependent sensitivity that we have defined earlier, hence providing the connection with TDSA.

Adjoint variables also provide a way to compute time-dependent *parameter* sensitivities (Cao et al., 2002). Consider a brief change in the value of the parameter *θ_i_* at time *t*, by which we mean a rapid change followed by rapid return to its original value (i.e., a spike or dip). This causes a brief change in 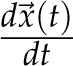 via Eqn. (1), which in turn leads to a sudden perturbation of *⃗x*(*t*). For example, a brief spike in the immigration rate is equivalent to a brief small translocation causing a sudden increase in the population size (see Fig. 2(D)). Hence, the sensitivities to a brief parameter perturbation can be inferred from the state sensitivities. Sensitivities to an arbitrary temporal pattern of perturbation can be calculated using Eqn. (A9), by treating the temporal pattern as a series of brief perturbations chained together (see Appendix B).

Time-dependent sensitivities are easy to compute numerically for the low-dimensional models we consider here. We first solve the state equations Eqn. (1) forward in time from 0 to *T* (using the **deSolve** package (Soetaert et al., 2010) in **R** (R Core Team, 2021)), saving values at a fine grid of times *t_k_* = *kT*/*n*, where *k* = 0,1,*···*,*n* with *n≫* 1. We then solve the adjoint equations Eqn. (5) backwards in time from *T* to 0 using approximate state variable trajectories obtained by linearly interpolating the values at times *t_k_*. We confirmed that this method works with simulations in which state variables were slightly perturbed by hand at various times. The effects of these perturbations on the value of *J* (integrals evaluated numerically by the trapezoid rule) always matched the predicted effect based on the state sensitivities *λ_i_*(*t*). Numerical methods for large-scale models are available (Cao et al., 2002, 2003).

Sensitivities allow us to compare between state variables the effects of perturbations by the same absolute amount. However, sometimes perturbations by the same *proportional* amount might be the more appropriate comparison, for example if the state variables differ vastly in scale, or if the potential management actions (e.g., spraying of pesticides) perturb a state variable (e.g., insect density) by an amount proportional to its value. The time-dependent *demi-elasticity*^1^ of state variable *x_i_* is defined as 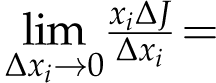 *x_i_*λ_*i*_. One can also calculate the elasticity, 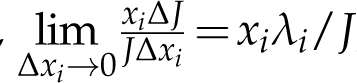, but these can be misleading if the reward function *J* represents deviations from a baseline value. For example, if the goal is to maximize plant yield by minimizing damage from herbivory, it is convenient to define the reward function *J* as *−*1 times the damage due to herbivory. In that case, because *J* differs from plant yield by a constant, the sensitivities and demi-elasticities of *J* would be the same as those of plant yield, but the elasticities would be different.

#### Discrete-time models

TDSA of discrete-time models is also motivated by its counterpart in optimal control (Lenhart and Workman (2007), Chapter 23). We consider a model that can be written as a system of forward recursions

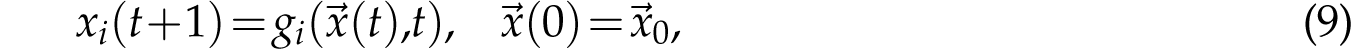

where *t* = 0,1,2,…,*T* denotes the time step, and *⃗x*(*t*) the state vector at time *t*, with *i*th component *x_i_*(*t*). We consider a reward function of the form

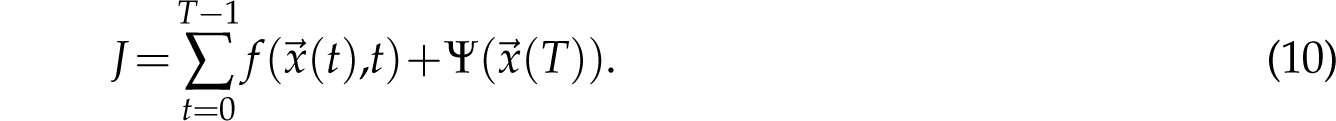

The contribution from the final time step has been separated from the rest, because it will be used later to determine the terminal conditions when solving the adjoint equations backward in time. We introduce an adjoint vector ^⃗^*λ*(*t*) with the same number of components as the state vector *⃗x*(*t*); the *i*th component *λ_i_*(*t*) gives the sensitivity of *J* to perturbations of the state variable *x_i_*(*t*) at time *t*. The adjoint vector satisfies the adjoint equations and terminal conditions

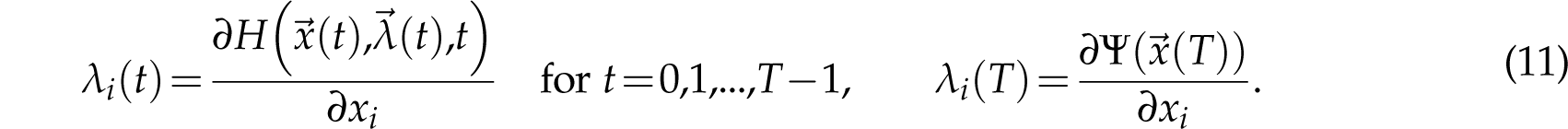

where the Hamiltonian *H* is defined as

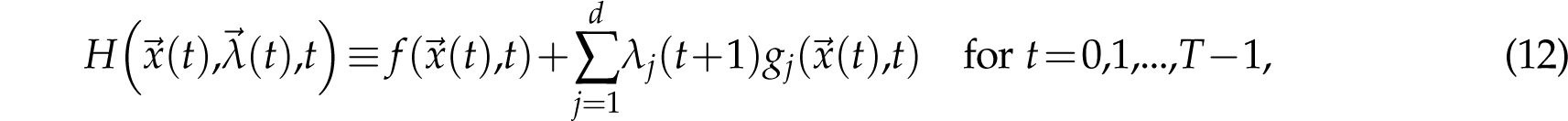

(Unlike Eqn. (5), here there is no minus sign in front of the derivatives of *H*.) Eqn. (11) is a system of backward recurrence equations, which we can solve backward in time to obtain the sensitivity *λ_i_*(*t*) at any time *t*.

### Applications and their Implications

Having explained how to calculate time-dependent sensitivities, we now embark on a series of applications to illustrate the potential payoffs from applying TDSA, and to point out some potential pitfalls. Our first case studies, in section *Example 1: Exogenous disease spillover in multi-species sink networks*, are theoretical examples designed to illustrate how state and parameter sensitivities can be strongly time-varying even if model equations and parameters are constant. Our second case study (section *Example 2: Leopard frogs as reservoirs of the amphibian chytrid fungus*) is an empirically-fitted model with periodic dynamics driven by seasonality, and shows how TDSA can identify the key period in the annual cycle—the timing of which may be surprising at first sight, but becomes intuitively clear in hindsight. This example also demonstrates how discretization allows us to apply TDSA to models with continuous independent variables such as Integral Projection Models (IPM). Our third case study (section *Example 3: Population cycles in the pine looper and the larch budmoth*) are empirically-fitted autonomous models with endogenously-driven oscillatory dynamics, and highlight some of the practical challenges in applying the results from TDSA to management actions. In both the second and third examples, we also demonstrate the importance of making an effort to interpret TDSA results rather than taking them at face value, to avoid drawing spurious conclusions that reflect aspects of the mathematical models but do not correspond to real biological phenomena.

### Example 1: Exogenous disease spillover in multi-species sink networks

#### Overview

Our first examples developed from our work on disease spillover (Ng et al., in press). We consider a multi-species sink community that cannot maintain a disease by itself; the disease only persists via spillover from an exogenous source. Disease transmission within the community is represented by a *static* unipartite network, meaning that the intra- and inter-species transmission coefficients are assumed to be constant parameters. The exogenous spillover rate is also assumed to be constant. The time-dependent phenomena of interest are the *transient dynamics* of disease spread within an active season; this is relevant if the disease is seasonal, in that it dies out in the sink community between one active season and the next (via an unmodeled process), but is re-introduced at the start of each active season via exogenous spillover.

We consider two hypothetical network designs. Although partially motivated by disease transmission in plant-pollinator communities (e.g. trait-matching networks from Truitt et al. (2019)), these networks are not meant to represent any specific empirical system. Rather, they were designed to illustrate how TDSA can highlight qualitative features in the dynamics induced by network structure. In each case, the objective is to reduce the negative disease impact on a species of concern in the sink community.

#### Mathematical model

We consider a community of *m* host species, where individuals can either be susceptible or infected. The state variables *S_j_*(*t*) and *I_j_*(*t*) represent the number of susceptible and infected individuals in species *j*, while *N_j_*(*t*) *≡S_j_*(*t*)+ *I_j_*(*t*) represents the species population. The dynamic equations are

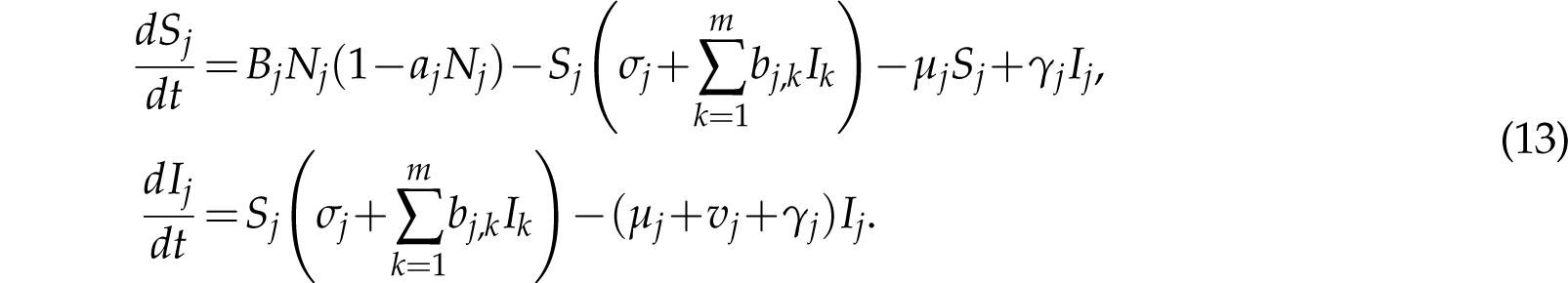

For species *j*, *B_j_* is the unregulated per-capita birth rate. We assume infection does not affect fecundity. We also assume only intra-specific competition for limiting resources necessary for reproduction (e.g., breeding sites), represented by the competition coefficient *a_j_*; the carrying capacity in a disease-free population is then given by *K_j_* =(1*−µ_j_*/*B_j_*)/*a_j_*. *µ_j_* is the mortality rate of a susceptible individual, *v_j_* the additional mortality rate arising from infection, and *γ_j_* the recovery rate. Within-community transmission is parametrized by *b_j_*_,*k*_ representing the transmission rate from a species-*k* infective to a species-*j* susceptible, while exogenous spillover is parametrized by *σ_j_* representing the per-capita spillover infection rate in species *j*. We assume no vertical transmission, so all individuals are born uninfected. Parameters were chosen so that the basic reproduction number *R*_0_ of the disease (Diekmann et al., 2013) is less than one, so that disease is only maintained in the sink community by the exogenous spillover.

#### Objective function

To create scenarios in which transient dynamics are important, we make the following assumptions. All species are active each year between *t* = 0 and *T* (the active season). All active individuals die at the end of the season, while a new generation of active individuals emerge disease-free at the start of next season. The population size at the end of one season influences the population size at the start of the next season.

In both hypothetical networks, we assume there is a species of concern (*j* = *j_C_*) that provides an important ecosystem service (e.g., being an efficient natural pollinator of a crop plant), but whose population is negatively impacted by the disease, due to a combination of the species being vulnerable (*B_jC_* only slightly greater than *µ_jC_*) and a high disease-induced mortality rate (*ν_jC_ ≫µ_jC_*). The goal of TDSA is to identify control measures that reduce infection in this species, to reduce the impact on the ecosystem service. The reward function *J* represents the economic value of the service, and is given by

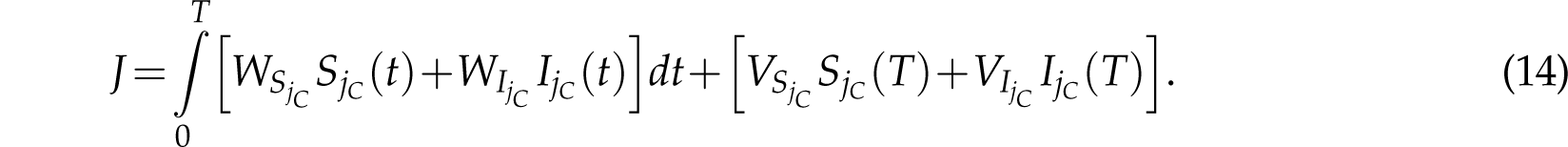

The integral represents the value of the service over the current season, assuming the service is equally valuable throughout (so *W_SjC_* and *W_IjC_* are constants), and scales linearly with the number of individuals. The terminal payoff terms represent the value of maintaining a large population at the end of the season, since this will affect the population size at the start of the next season. Since both susceptible and infected individuals are equally fecund and produce healthy offspring, *V_SjC_* = *V_IjC_*. For notational simplicity, we also introduce coefficients *W_Sj_*, *W_Ij_*, *V_Sj_* and *V_Ij_* for other species (*j ̸*= *j_C_*) but they are all zero.

#### Adjoint equations

From Eqns. (5–6), the adjoint variables satisfy

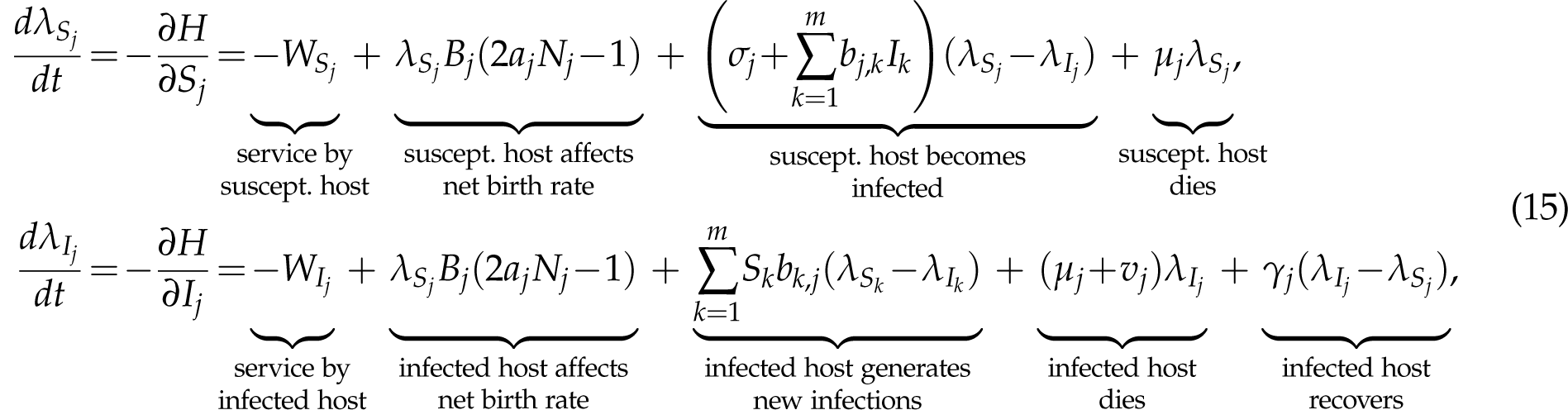

with terminal conditions

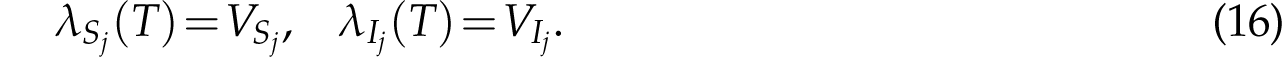

To interpret each term in Eq. (15), recall that *λ_Sj_* (*t*) is the shadow price of a susceptible host of species *j* at time *t*. If *dλ_Sj_* /*dt* is negative, *λ_Sj_* (*t*) will increase when *t* decreases. Hence negative terms in *dλ_Sj_*/*dt* tend to increase the reward from adding a susceptible host at an earlier time. For example, focusing on the species of concern (*j* = *j_C_*), the first term *−W_SjC_* is negative since the earlier we add the host, the more service the host can provide before the season ends at time *T*. Conversely, the last term *µ_j_λ_Sj_* is positive, since the earlier we add a susceptible host, the more likely it dies before time *T*, hence limiting the amount of service provided (which would otherwise have grown linearly as *T−t* when *t* decreases), as well as reducing its likelihood of contributing to the terminal payoff *V_SjC_*. The third term depends on the sign of *λ_Sj_−λ_Ij_*, since the earlier we introduce a susceptible host, the more likely it becomes infected before time *T*, thus changing its shadow price to that of an infected host. Similar interpretations can be made for the other terms in *dλ_Sj_* /*dt* and *dλ_Ij_* /*dt*.

We assume that all species other than the species of concern are not at risk (*B_j_ ≫µ_j_* for *j ̸*= *j_C_*), so one possible control measure is to cull these species to slow down the spread of infection. Because culling is often performed indiscriminately regardless of infection status, it is useful to examine the sensitivity of the reward *J* to *random removal* of individuals from species *j ̸*= *j_C_*. This is given by *−λ_Nj_*, where *λ_Nj_* is the adjoint variable for the total population size. Although *λ_Nj_*can be formally derived by working with *N_j_* and *p_j_ ≡ I_j_*/*N_j_* (the infection prevalence) instead of *S_j_* and *I_j_* as the state variables, one can show that

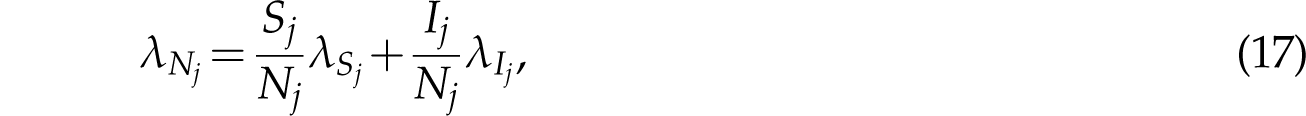

a rather intuitive expression. In Online Supplement Sec. S3, we derive a general expression for the change of adjoint variables under a change of state variables.

For simplicity, we assume that only one species *j* = *j_E_* receives exogenous infection. This makes it easier to interpret how sensitivities reflect network structure. Parameter values (stated in Online Supplement Sec. S4) were chosen to illustrate interesting features in time-dependent sensitivities which may be less obvious at other parameter values.

#### Network 1: Nearest-neighbor network

As our first hypothetical network, we consider a community of *m* = 5 species with only nearest-neighbor interactions as shown in Fig. 3A. (See Fig. S1(A) for the matrix representation of this network.) This can be thought of as an extreme example of a trait-matching network (Truitt et al., 2019), where each species only interacts with other species that are adjacent along a one-dimensional trait space. Exogenous spillover occurs in species 1 (*j_E_* = 1) while species 5 is the species of concern (*j_C_* = 5), so the disease will have to be progressively relayed from species 1 to 5 via the intermediate species. Indeed, we see in Fig. 3(C) that the highest rate of infection per capita (maximum 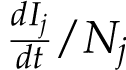, indicated by the dots) occurs at a later time for a species further down the network. Despite low disease prevalence in species 5, the fact that it is vulnerable (due to a low excess of births over natural mortality) means that the population decrease across a season can be rather substantial, and the cumulative decrease over multiple years quite large, as shown in Fig. S1(B), hence creating the need for control measures.

**Figure 3:**
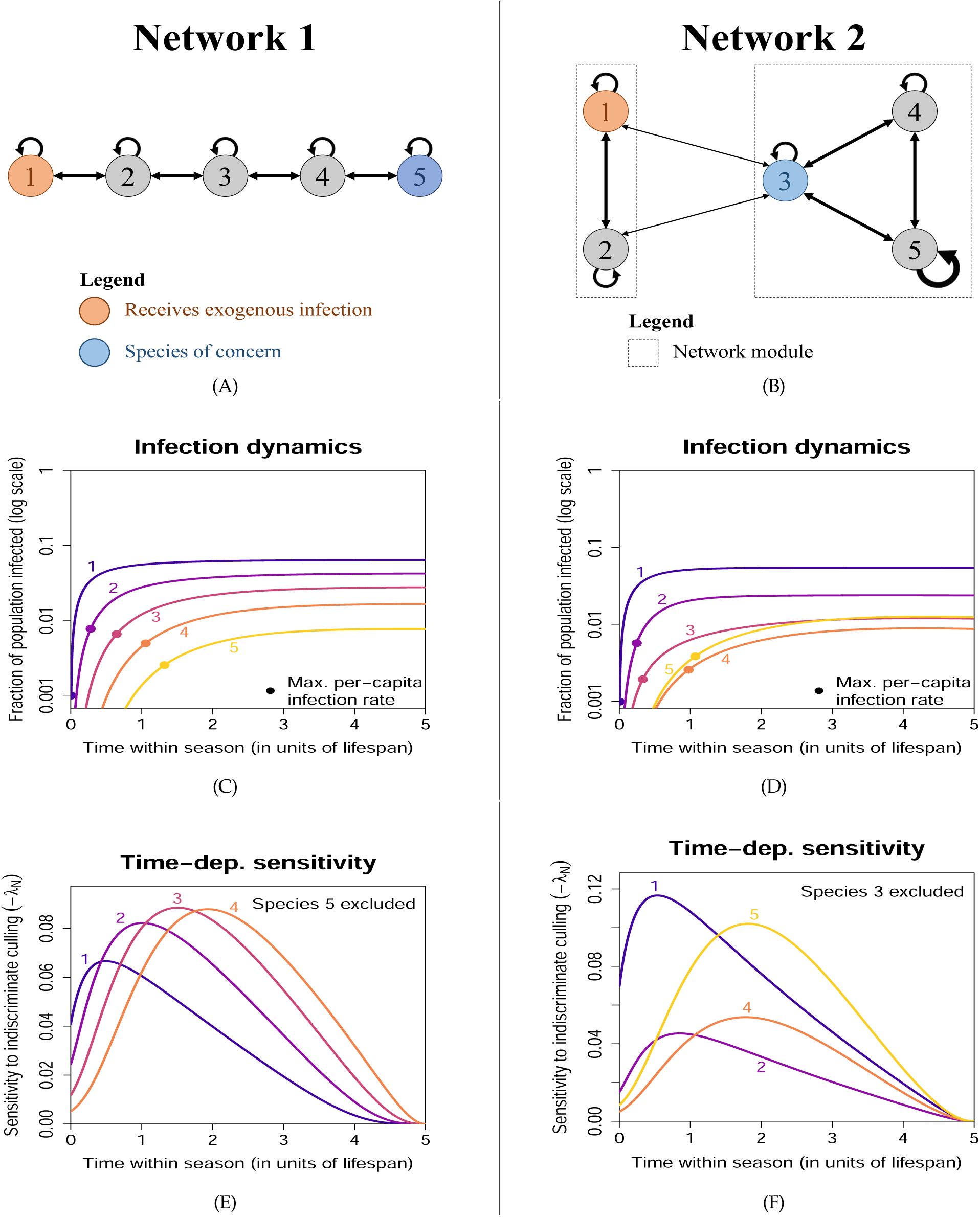
Time-dependent sensitivities to state perturbations from Example 1. Each column corresponds to one network configuration. Disease spread is subcritical (*R*_0_ <1), so the disease is maintained by exogenous spillover. **(A)** and **(B)**: Infection pathways; line thickness roughly scales with the size of the transmission coefficients *b_j_*_,*k*_. **(C)** and **(D)**: Disease prevalence in each species. Each dot indicates when the per-capita rate of infection is the highest (note the vertical log scale). Despite the low infection prevalence in the species of concern, the population decline from disease-induced mortality can be substantial (see Figs. S1(B) and S2(B)). **(E)** and **(F)**: Sensitivity of the reward function to indiscriminate culling (removal of random individuals regardless of infection status) of each species, excluding the species of concern.

The sensitivity of each intermediate species should exhibit a peak in time. For example, culling species 4 is ineffective at the start of the season because its population size would have mostly recovered before disease prevalence starts to increase. Culling becomes much more effective when the chain of infection reaches species 4, because removing susceptibles has an immediate impact on density-dependent transmission, and also because indiscriminate culling removes more infected individuals. Culling species 4 becomes ineffective again late in the season, because there is little time for species 5 to benefit from the reduced infection rate before the season ends (see Fig. S3). Because the time of peak sensitivity varies with species, the optimal species to target should vary over the active season. Fig. 3(E) shows that this is indeed the case: the most important species changes progressively from species 1 *→* 2 *→* 3 *→* 4. The progression in the most important species from 1 *→* 2 *→* 3 *→* 4 also depends on the fact that the peak sensitivities are of comparable height; otherwise a species may remain unimportant even at its peak sensitivity if the peak is low. Why does this occur? An infected individual in a species further down the chain of infection is more likely to cause infection in species 5 than one further up the chain (hence the large differences in *−λ_Ij_* shown in Fig. S1(D)). However, the per-capita rate of infection is also lower for a species further down the chain. These opposing effects lead to comparable peak heights in *−λ_Sj_* as shown in Fig. S1(C). Also, the indiscriminate culling sensitivity *−λ_Nj_* is a weighted sum of *−λ_Sj_*and *−λ_Ij_*, and the lower prevalence down the chain means a lower weight for *−λ_Ij_*, which again opposes the higher value of *−λ_Ij_*.

#### Network 2: Modular network with disease spillback

In this network, we consider *m* = 5 species grouped into two modules as shown in Fig. 3(C). Modules might also arise from trait-matching, where each module is associated with specialization on a particular resource type, and indirect interactions via the shared resource type lead to within-module disease transmission. Exogenous spillover again occurs in species 1 in the first module (*j_E_* = 1), while species 3 in the second module is the species of concern (*j_C_* = 3). However, we also choose species 3 to interact weakly with the first module (for example, it may be less specialized), and hence bridge disease transmission between the two modules.

Unlike Network 1, here other species are not needed to relay the disease from species 1 to 3, and in fact species 3 is the one relaying to the rest of the second module. Therefore, one would expect species 1 to remain the most important species (highest sensitivity) throughout the season. However, suppose that species 5 in the second module is a highly social species with strong within-species transmission (indicated by the thicker self-loop in Fig. 3(B)). This allows species 5 to reach high disease prevalence. “Spill-back” from species 5 to species 3 may then become more important than the transmission from species 1. In Fig. 3(F), we see that indeed species 5 becomes the most important later in the season. However, this relies on *R*_0_ being sufficiently close to 1 (Fig. S4(A)), so that multi-step within-module transmission can occur. We also find that at a higher exogenous spillover rate *σ*_1_, the most important species may switch back to species 1 towards the end of the season (Fig. S4(B)). This is because the higher spillover rate leads to a large decline in the population of species 3, which affects multi-step within-module transmission (recall that transmission is density-dependent in our model) and hence reduces the importance of species 5. To confirm this explanation, we replaced disease-induced mortality by recovery in species 3 (so that there is negligible population decline) and found that this switch no longer occurred (Fig. S4(C)).

#### Time-dependent parameter sensitivities

As explained earlier, time-dependent parameter sensitivities can be obtained from the adjoint variables using Eqn. (A9). We demonstrate this using Network 1, the nearest-neighbor network. First, we consider what happens if we increase the mortalities *µ_j_* briefly between *t*_0_ and *t*_1_ (e.g. via culling). This perturbation can be written as *µ_j_ →µ_j_* +*ɛ*Θ(*t*), where Θ(*t*) is a normalised indicator function equal to 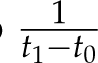 if *t∈* [*t*_0_,*t*_1_], and 0 otherwise, and *ɛ* is a small parameter representing perturbation size. Using Eqn. (A9), the time-dependent parameter sensitivity for *µ_j_* is given by the integral

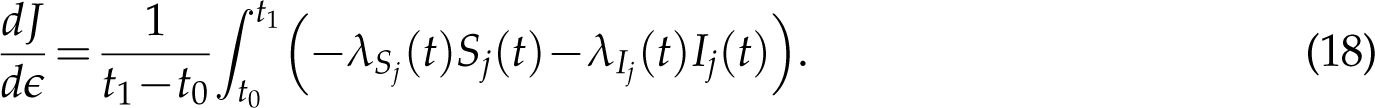

In Fig. 4(A) we show the sensitivities for different choices of the start time *t*_0_, assuming a window length *t*_1_*−t*_0_ = 0.2. Since the integrand is proportion to *−λ_Nj_* and the integration window is relatively short, not surprisingly, the results are qualitatively similar to *−λ_Nj_* as shown in Fig. 3(G).

**Figure 4:**
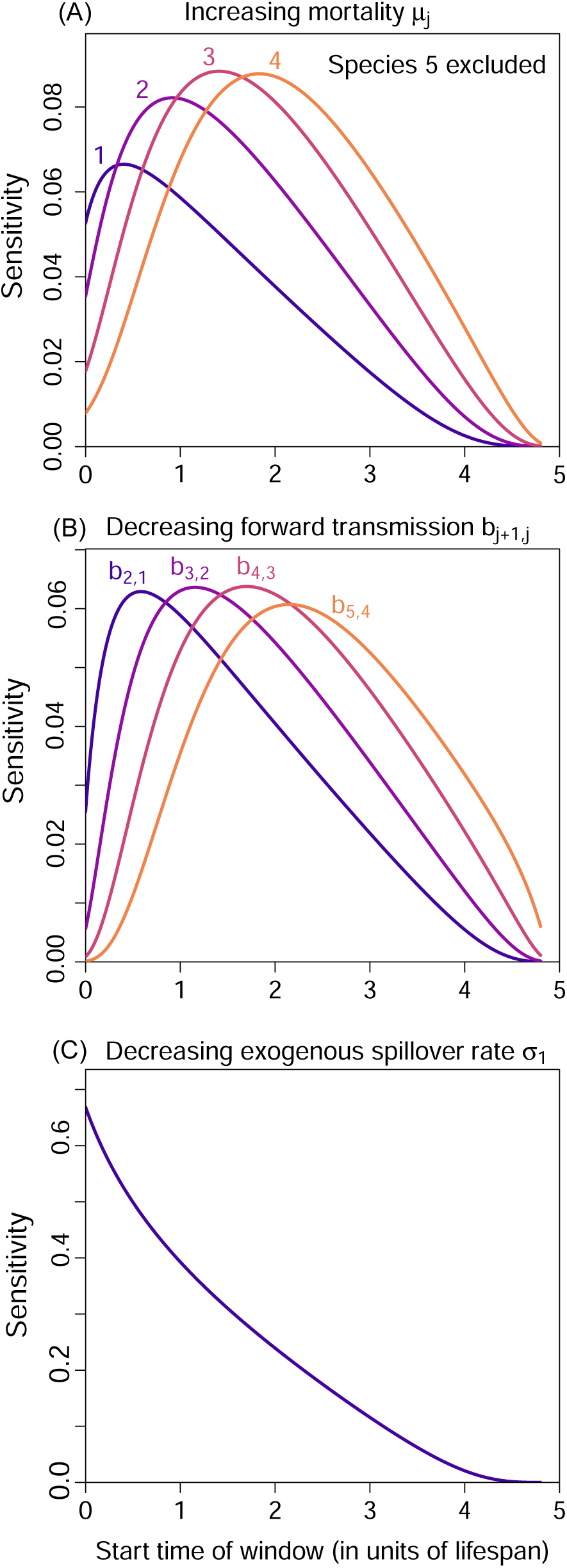
Time-dependent parameter sensitivities for the nearest-neighbor network (Example 1, Network 1). We consider brief perturbations of 0.2 time units to **(A)** the mortality rates *µ_j_*, **(B)** the forward transmission rates *b_j_*_+1,*j*_, and **(C)** the exogenous spillover rate *σ*_1_ to Species 1. Each panel shows the sensitivity of the reward to the perturbation as we vary the start time of a short perturbation window.

Next, we consider a decrease in the forward transmission rates *b_j_*_+1,*j*_ along the network, again over a short time window; this may arise from measures taken to briefly reduce contact between species in the network. For the perturbation *b_j_*_+1,*j*_ *→b_j_*_+1,_*_j_−ɛ*Θ(*t*), the sensitivity is given by

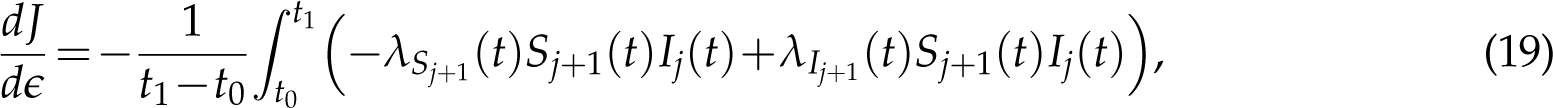

and the results at varying start times *t*_0_ are shown in Fig. 4(B). We see that targeting transmission links further down the chain is more effective at later times.

Finally, it might be possible to directly target the source of exogenous spillover, so we consider what happens if we briefly decrease the exogenous spillover rate *σ*_1_. Fig. 4(C) shows the sensitivities at varying start times *t*_0_. Unlike *µ*_1_ and *b*_2,1_, here the sensitivity is maximized at the start of the active season. To understand why, since disease prevalence in all species is zero at the start of each active season, culling Species 1 or reducing transmission from Species 1 to 2 become more effective after a slight delay as the disease re-establishes in Species 1, but this buildup of infection is irrelevant for *σ*_1_.

### Example 2: Leopard frogs as reservoirs of the amphibian chytrid fungus

In this second example, we demonstrate how TDSA can be applied to an integral projection model (IPM Ellner et al., 2016) with seasonal dynamics, by discretizing the continuous structure in the IPM into discrete bins. Wilber et al. (2022) proposed a series of models invoking different factors to explain the seasonal dynamics of the fungal pathogen *Batrachochytrium dendrobatidis* (*Bd*) in two species of North American leopard frogs, *Rana pipiens* and *Rana sphenocephala*. The models incorporate seasonal movements between aquatic and non-aquatic habitats, seasonal breeding, temperature-dependent pathogen load dynamics on infected frogs, and temperature-dependent zoospore survival in the water. Wilber et al. (2022) focused on reduced compartment models derived from the full model using moment closure approximations, in order to allow model fitting by Markov Chain Monte Carlo. But here we choose to work with the full model, because TDSA on the full model is not computationally burdensome even with fine discretization of the continuous population structure.

The IPM proceeds in steps of one week, with state variables *L*(*t*), *S*(*t*), *I*(*x*,*t*), and *Z*(*t*), representing larvae (tadpoles), susceptible adults, infected adults with log-transformed pathogen load *x*, and zoospores. Each year, all adults are in a shared aquatic habitat during the breeding season; otherwise, they are nonaquatic. Half the adults are female, and each female produces *r^′^*tadpoles at the midpoint of the breeding season. Each tadpole has a probability *s_L_* of surviving each week, and a probability *m_L_* of undergoing metamorphosis. However, recruitment is density-dependent, so only a fraction *e^−KN^*^(^*^t^*^)^ of these metamorphosed tadpoles successfully become adults, where 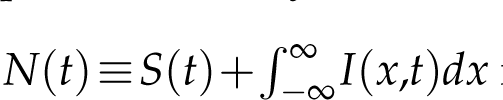 is the total number of adults. Susceptible and infected adults have survival probabilities *s*_0_ and *s*_0_*s_I_* respectively.

When aquatic, susceptible frogs become infected with a probability 1 *− e^−βZ^*^(^*^t^*^)^ that increases monotonically with *Z*(*t*). Newly-infected frogs have log load *x* drawn from a distribution *G*_0_(*x*), with a mean *a*(*T*(*t*)) that decreases linearly with the temperature *T*(*t*). For an already-infected frog with log load *x*, the new log load *x^′^* at the next timestep is drawn from a distribution *G*(*x^′^|x*), with a mean *a*(*T*(*t*))+*bx*. Infected adults also have a probability ℓ(*x*) of recovery that decreases monotonically with *x*. When aquatic, infected frogs shed an amount of zoospores each week proportional to their linear load *e^x^*, with proportionality constant *λ*. Zoospores survive each week with a probability *s_Z_*(*T*(*t*)) that decreases monotonically with the temperature *T*(*t*). *T*(*t*) varies sinusoidally across the year, being the lowest at the start/end of the year and the highest mid-year. Finally, zoospores are also being added at a constant rate *ω* from exogenous sources not represented in the model. Altogether, we obtain the equations

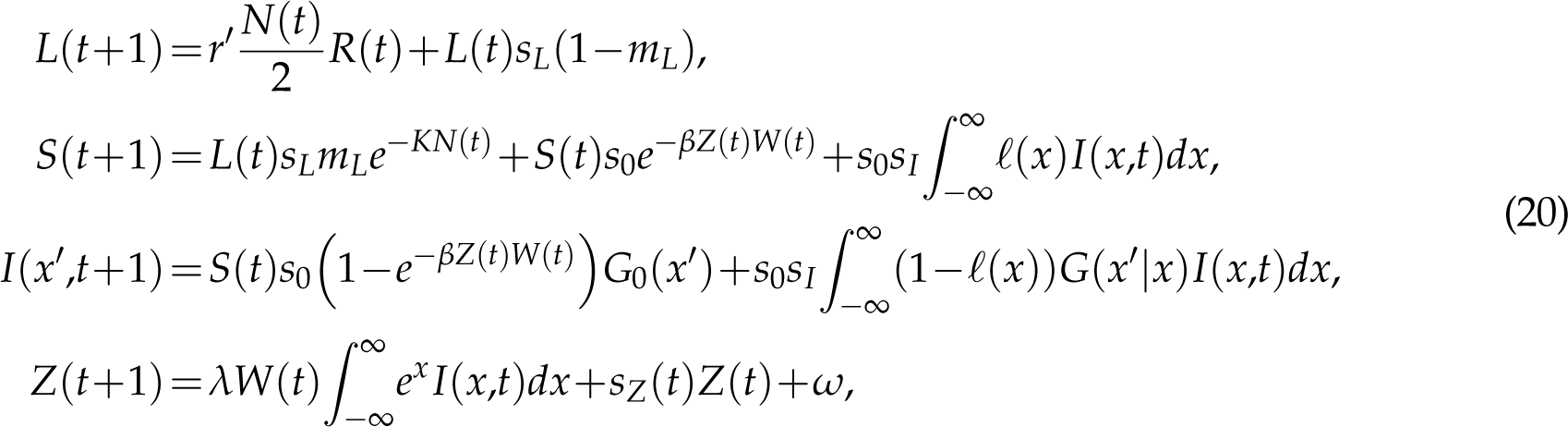

where both *W*(*t*) and *R*(*t*) are periodic indicator functions that can take values *{*0,1*}*; *W*(*t*) = 1 when the adults are aquatic, while *R*(*t*) = 1 at the midpoint of the breeding season (where new tadpoles are produced). More details can be found in Online Supplement Sec. S5.1.

Fig. 5 shows the steady-state dynamics of the *Bd*-bullfrog system. At the start of the breeding season, adults that still carry infection from the previous year return to water and shed zoospores, leading to a rapid increase in *Z*, which in turn causes a rapid rise in the number of infected adults and depletion of susceptibles. The midseason production and metamorphosis of larvae leads to a small jump in the number of susceptible and infected adults. Towards the end of the breeding season, higher temperature decreases the log *Bd* load on infected frogs (see Fig. S5), which in turn decreases shedding. This, together with the lower zoospore survival at higher temperatures, causes *Z* to decrease.

**Figure 5:**
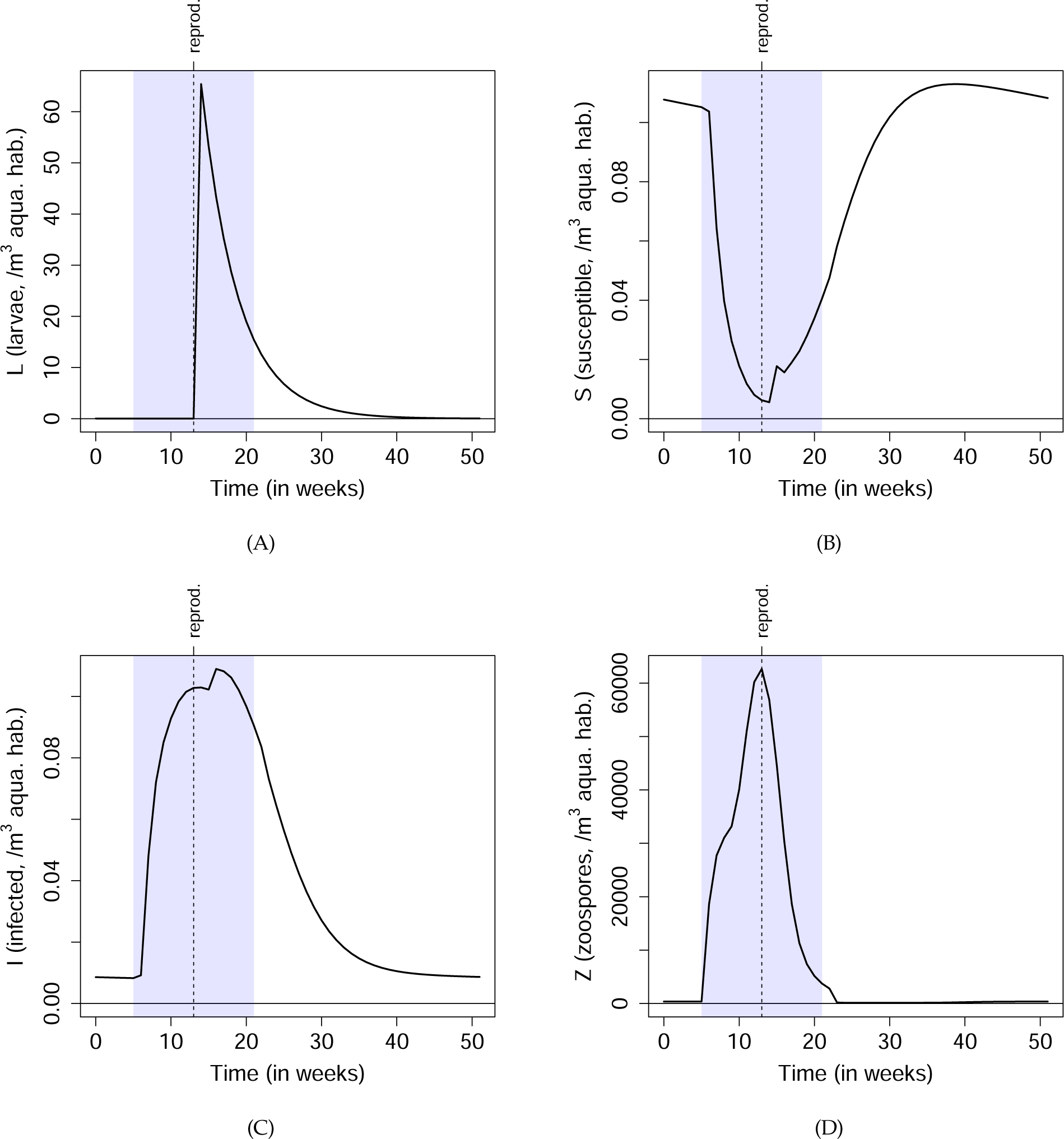
Steady-state model dynamics of the *Bd*-bullfrog system. Details of the model are described in Sec. *Example 2: Leopard frogs as reservoirs of the amphibian chytrid fungus*. The blue shaded regions show the period of each year when the bullfrogs are aquatic and exposed to potential infection, and the vertical dashed lines show when new bullfrog larvae are produced. *I*(*t*) in the bottom-left panel is the total number of infected frogs, ^J ∞^ *I*(*x*,*t*)*dx*.

Now consider a scenario where other vulnerable amphibian species of concern, also susceptible to *Bd*, share the same aquatic habitat with the bullfrogs. Therefore, we want to minimize the exposure of these species to zoospores during their breeding seasons. A possible objective function is given by

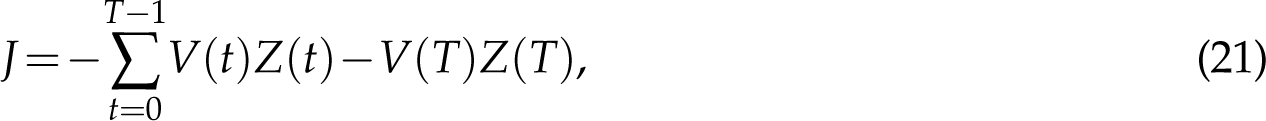

where *V*(*t*) is a periodic indicator function; *V*(*t*) = 1 when the vulnerable species are aquatic. The negative sign is so that maximizing the objective function minimizes exposure to *Bd*. Because we want to protect the vulnerable species as long as possible, ideally we would like the time horizon *T* to be infinite. In practice, since the effects of any small perturbation are expected to die off over time, and since each year starts off in the same state (assuming any transients have died off), the seasonal sensitivity patterns in the first few years become nearly identical and independent of *T* as long as *T* is sufficiently large (see Online Supplement Fig. S7); hence they approximate the seasonal patterns when *T→* ∞.

We now discretize the IPM into *m* log-load bins of width *h* each. Details of the discretized model are presented in Online Supplement Sec. S5.2. We apply Eqns. (11–12) to obtain the adjoint equations

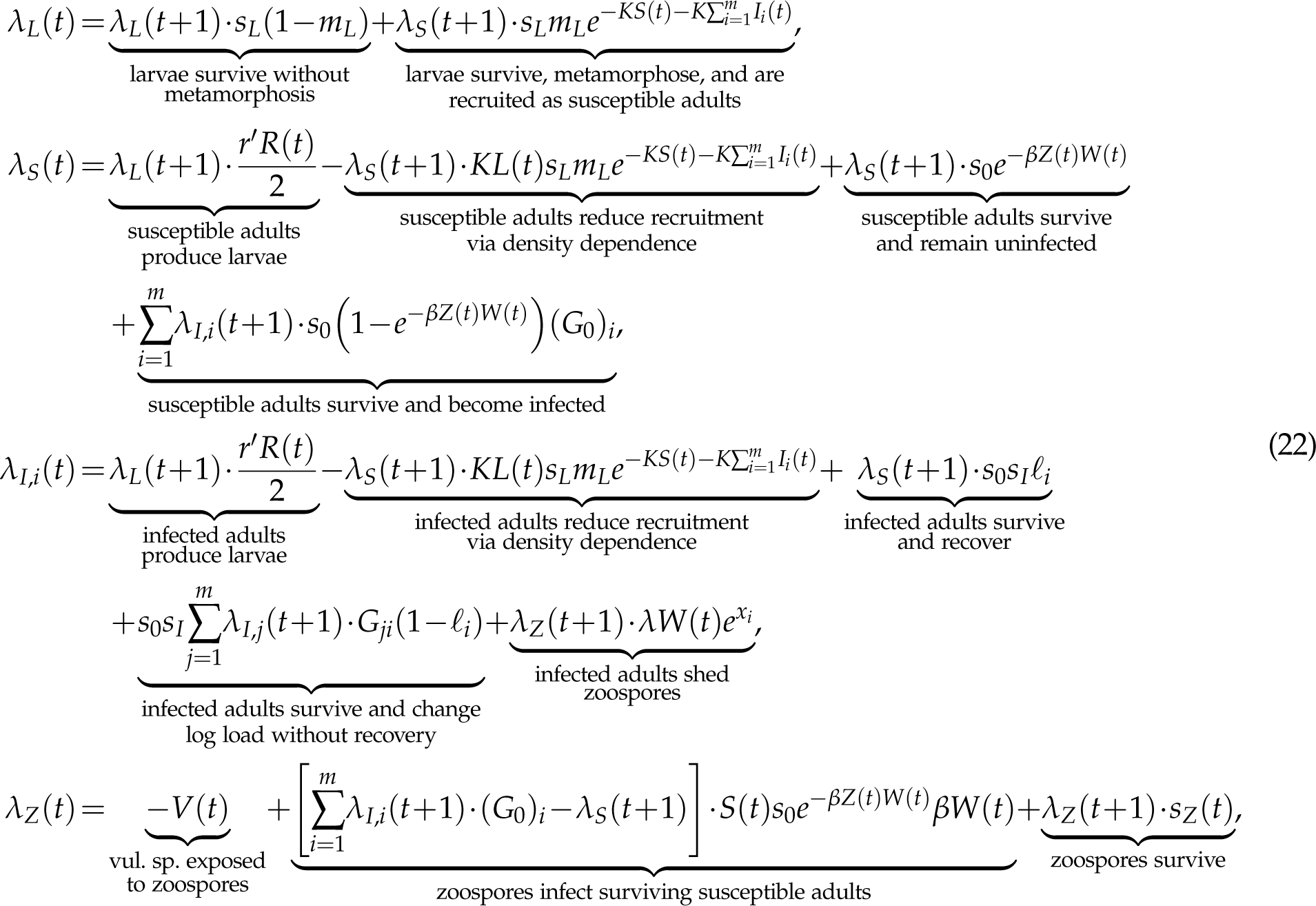

and terminal conditions

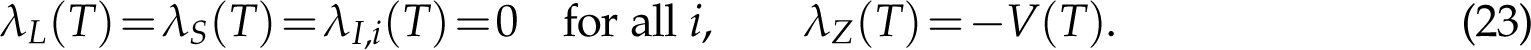

The adjoint variable *λ_I_*_,*i*_(*t*) represents the effect of perturbing *I_i_*(*t*), the number of infected adults in bin *i* (with log load *x_i_*) at time *t*; see Fig. S6. However, it is probably more realistic to consider the effect of, say, removing an infected individual sampled at random. Therefore, we introduce the sensitivity *λ_I_*, defined as a weighted average of *λ_I_*_,*i*_(*t*) with weight proportional to *I_i_*(*t*):

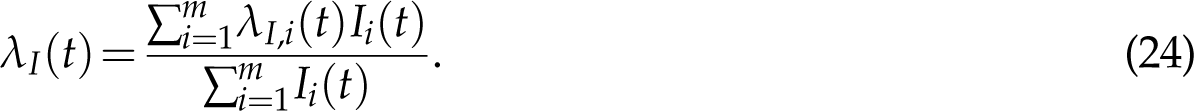

Fig. 6 shows the sensitivities when the vulnerable species have the same breeding season as the bullfrogs, so *V*(*t*) = *W*(*t*). For easier visualization, we have plotted the negative of the sensitivities, so a positive plotted value means that increasing the state variable increases the exposure of vulnerable species to zoospores. Also, we have only shown the first year out of a time horizon of ten years; the patterns are similar in the first few years, so we can think of the patterns as being periodic.

**Figure 6:**
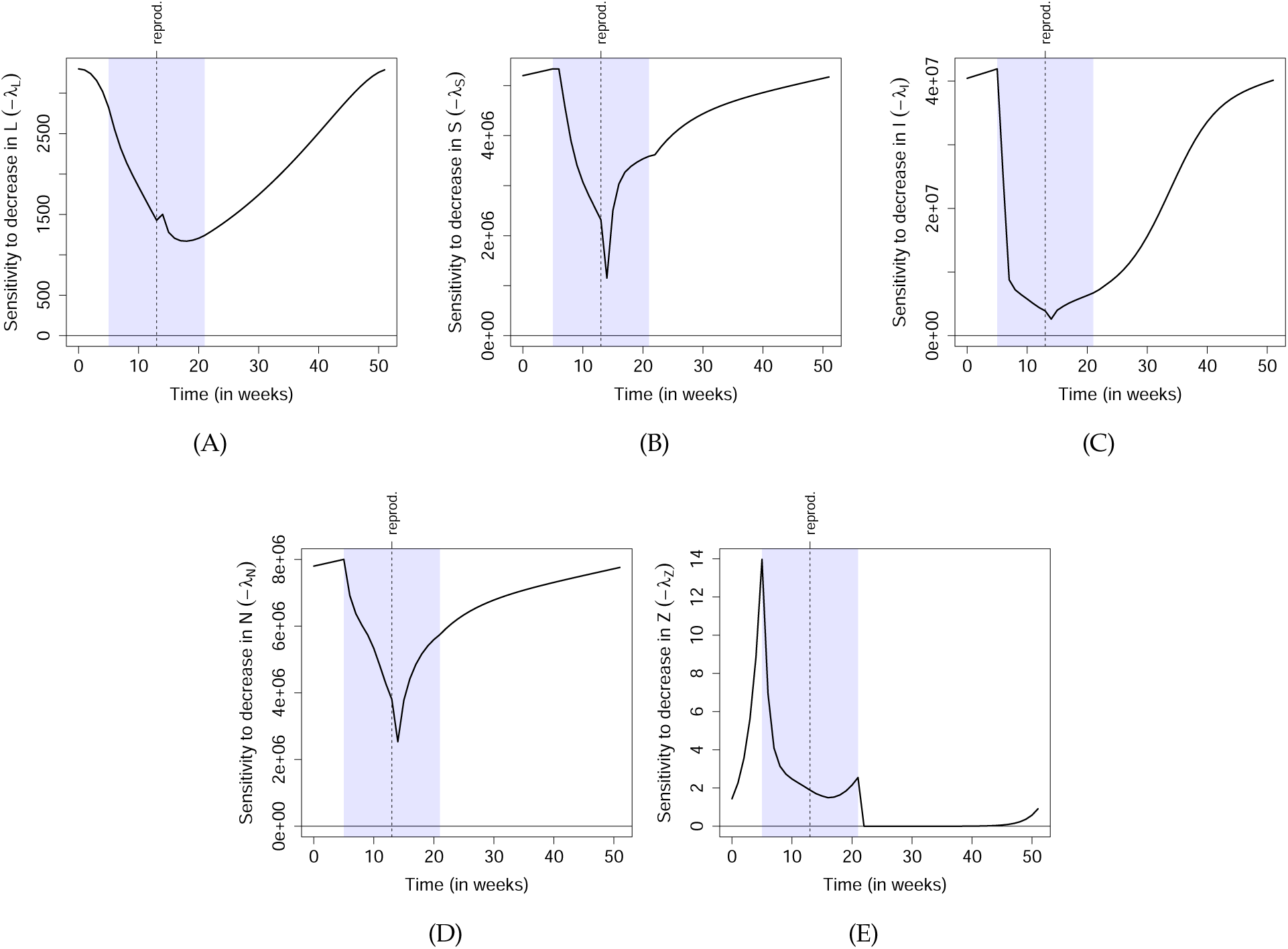
Time-dependent sensitivities of the *Bd*-bullfrog system. See the main text for the scenario and objective function. We assume that the vulnerable amphibian species of concern share aquatic habitats with bullfrogs during the same time period (the blue shaded region). Here, we only show the sensitivities in the 1st year (out of a time horizon of 10 years). Since the patterns in the few years are nearly identical, we can consider these patterns as periodic (i.e. the last week wraps around to the first week), and representative of an infinite-horizon scenario.

It is often said that “all models are wrong but some are useful” (Box, 1979). TDSA is a rigorous mathematical procedure applied to a user-specified model, but it cannot automatically distinguish between the wrong and the useful parts of a model. Hence, a practitioner should make an effort to interpret the important qualitative features in the sensitivities, and not just accept the results without question. Features that depend only on broad, qualitative model assumptions conceptualize the known biology of the system are more likely to be realistic and useful, while others may depend on (possibly questionable) model details often chosen for mathematical simplicity. For example, the sharp dips in *−λ_S_*(*t*) and *−λ_I_*(*t*) at *t* = *t*_repro._+1 are probably questionable. They result from density dependence in recruitment assuming that all new larvae appear simultaneously at *t* = *t*_repro._+1, and that some larvae can metamorphose in the next time step without delay. Because these detailed assumptions were likely chosen for simplicity rather than realism, the consequent features are unlikely to be realistic, and hence should not be taken literally when making management decisions.

As an example of a more realistic feature, we observe that *−λ_I_*(*t*) is lowest around the middle of breeding season, even compared to when the adults are non-aquatic. This only relies on the broad property (also present in the data; see Fig. 2 in Wilber et al. (2022)) that an infected adult introduced early in the season contributes many times more zoospores to the water than one introduced mid-season, directly because of its higher load (from the temperature-dependent load dynamics), and also indirectly because of the greater availability of susceptible adults that it can infect (since susceptibles become depleted mid-season). Hence, even though an infected adult introduced mid-season can immediately shed zoospores, an infected adult introduced after the breeding season is more likely to reach the start of the next breeding season alive and infected, simply by being closer to next season. The increased probability to contribute at the start of next season is more than enough to make up for not contributing immediately. This feature relies on less specific assumptions and is hence more likely to be realistic, although the modeler will still need to decide based on available knowledge. Note that the sensitivities need not reflect the relative efficiency of management action—for example, non-aquatic frogs may be harder to locate.

Although we have only presented the sensitivities in Fig. 6, a manager should consider whether sensitivities or demi-elasticities better reflect the costs and benefits of management actions. Demi-elasticities are more useful for actions whose direct effects on the state variables (e.g. population size) scale with the size of the state variables; for example, field capture of diseased frogs will probably yield more frogs per unit of time effort at higher frog densities. For state variables that exhibit temporal variations spanning many orders of magnitude, the demi-elasticities may show qualitative features that are completely different from the sensitivities, so while the sensitivities are still technically correct, their practical relevance may be limited. As an extreme example, we observe that the larval sensitivity *−λ_L_*(*t*) peaks in winter before the breeding season. This is because the model assumes that the larvae parameters *s_L_*and *m_L_* remain constant throughout the year, so a hypothetical tadpole introduced in winter has a good chance of metamorphosing into a susceptible adult around the start of breeding season, hence maximizing its contribution to zoospores through infection and shedding. On the other hand, this result is rather jarring since one is unlikely to find tadpoles in winter. By looking at the demi-elasticity, we take larval density into account and avoid this feature entirely.

Finally, whenever we discretize an IPM, it is good practice to check that the number of bins is large enough to approximate the continuum limit, by repeating the calculations with varying number of bins (see Online Supplement Fig. S8). We also recommend calculating the sensitivities directly by simulating explicit perturbations, to check that the adjoint equations were derived and implemented correctly (see Online Supplement Fig. S9). While directly calculating the sensitivities for all state variables at all time points may be computationally prohibitive (which is why the adjoint method is useful in the first place), one can still perform checks at a few time points of interest.

### Example 3: Population cycles in the pine looper and the larch budmoth

As our final examples we consider two models, both involving moth species that exhibit population cycles and cause forest defoliation in years of high abundance. The first model is a single-patch model of the pine looper, and the second is a spatially-explicit, multi-patch model of the larch budmoth. Both are discrete-time models with steps of one year. We present the pine looper model in detail in this section, and leave the details of the larch budmoth to Online Supplement Sec. S6.1 and S6.2.

#### Pine looper

The pine looper moth, *Bupalus piniarius*, exhibits large population cycles in parts of Europe, and can defoliate pine forests and plantations during outbreaks. While numerous explanations have been proposed for these cycles, Kendall et al. (2005) found that the maternal effects hypothesis had the strongest empirical support. The maternal effects model, a discrete-time model with steps of one year, is given by

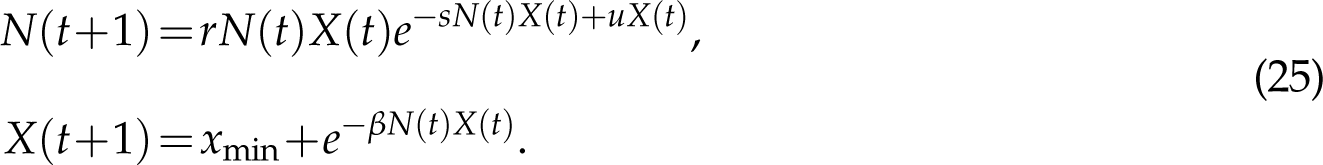

Here, *N*(*t*) is the density of pupae at year *t*, and *X*(*t*) a measure of their average individual quality. A constant proportion of pupae is assumed to survive to adulthood, so *N*(*t*) is a proxy for the adult abundance in that year. *X*(*t*) influences the per-capita fecundity, so the total number of offspring produced is proportional to *N*(*t*)*X*(*t*) in the first equation. As a maternal effect, *X*(*t*) also influences the probability of the offspring surviving from egg to adulthood the next year via the factor *e^uX^*^(^*^t^*^)^ in the first equation. Meanwhile, competition between the offspring reduces the probability of surviving to adulthood and also their average individual quality via the factors *e^−sN^*^(^*^t^*^)^*^X^*^(^*^t^*^)^ and *e^−βN^*^(^*^t^*^)^*^X^*^(^*^t^*^)^ in the first and second equations respectively. Kendall et al. (2005) fitted the model to data from three forest sites in Scotland: Roseisle, Tentsmuir and Culbin (Fig. 1B); see Table S1 for parameter values. As shown in Fig. 7(A), this model leads to oscillations in pupae density. In years of low pupae density, reduced competition between their offspring leads to an increase in offspring individual quality. The consequent increase in per-capita fecundity and egg-to-adult survival probability then leads to a population boom. The increased competition between offspring in the boom years then greatly reduces the individual quality and causes the population to crash, completing the oscillation. We note that the phase space trajectories are periodic in Roseisle (one complete cycle comprises two consecutive oscillations), and appear to be quasiperiodic in Tentsmuir and Culbin; see Fig. S10.

**Figure 7:**
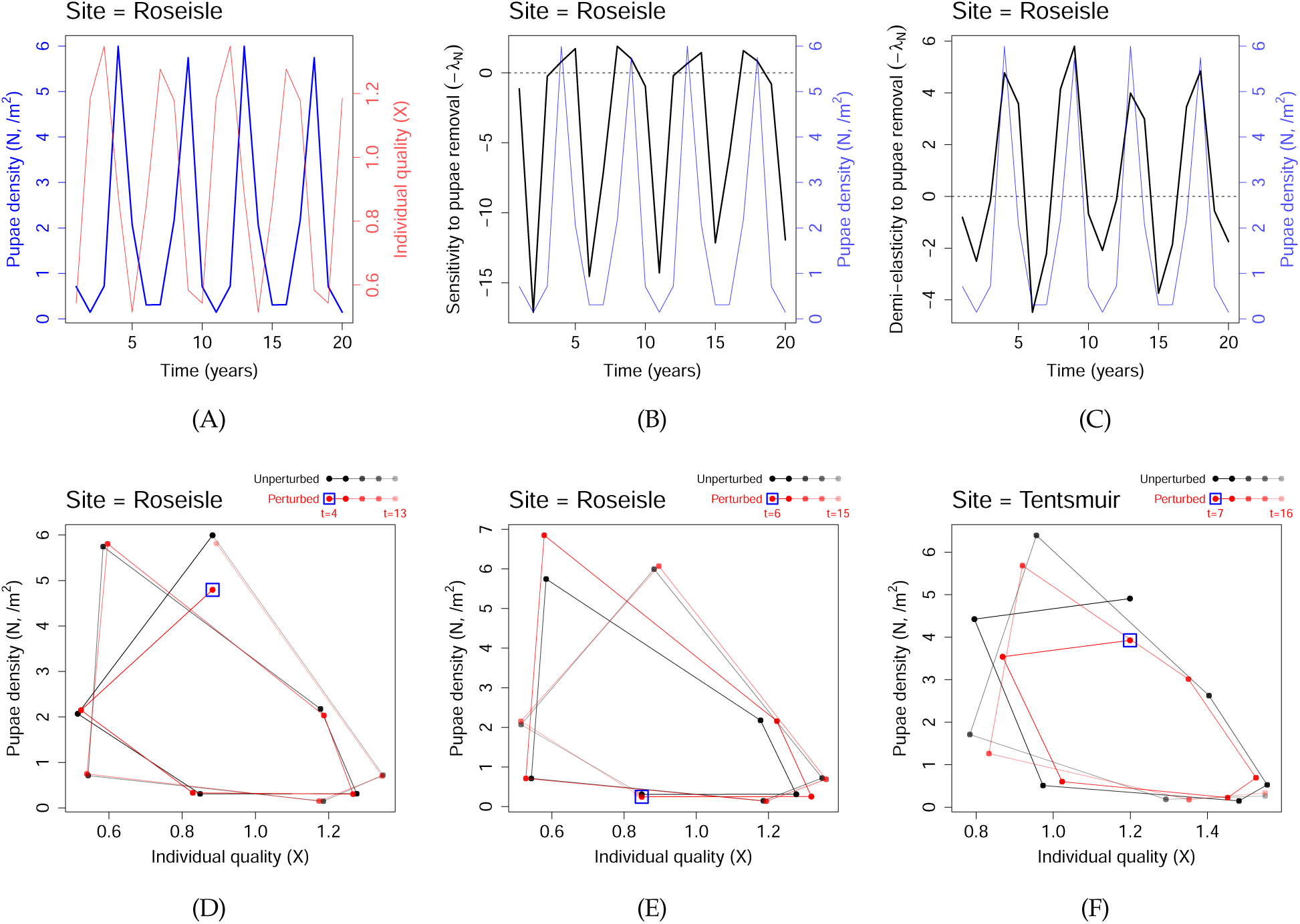
Dynamics and TDSA of the pine looper model. **(A)** Oscillatory dynamics in the pine looper, based on parameters estimated for Roseisle forest, Scotland. The blue line indicates pupae density, and the faint red line the average individual quality. **(B), (C)** The sensitivity and demi-elasticity to *culling* of pine looper, defined as *−λ_N_*(*t*) and *−N*(*t*)*λ_N_*(*t*) respectively. The reward function here is related to minimizing herbivory damage by the pine looper, see Eqn. (26). The pupae densities are plotted again in faint blue lines to facilitate comparison. **(D), (E)** Phase plane diagrams when 20% of the moths were culled in Roseisle at *t* = 4 (a positive demi-elasticity peak) and at *t* = 6 (a negative demi-elasticity valley). The black and red trajectories indicate the unperturbed and perturbed trajectories. The blue square highlights the start of the perturbed trajectory. **(F)** Similar to (D), except for the site Tentsmuir at *t* = 7 (a positive demi-elasticity peak). In (D-F), the differences between the vertical positions of the red and black dots are relevant to the change in reward function.

We now apply TDSA to the maternal effects model. To do so, we need to define the reward function. If we assume that tree damage is proportional to moth density, then a natural definition of the reward function will be 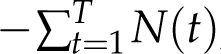, where *T* is the time horizon (note the overall minus sign). However, the quasiperiodic steady-state solutions for Tentsmuir and Culbin means that the effects of a state perturbation may persist indefinitely without damping out, as shown in Fig. S11. This is because when the system returns to the quasiperiodic solution after a small perturbation, it may be phase-shifted relative to the unperturbed trajectory, so the difference between the original and perturbed trajectories never damps to zero. This means that the change in reward function may depend on the choice of time horizon *T*. To avoid this, we choose a time-discounted reward function given by

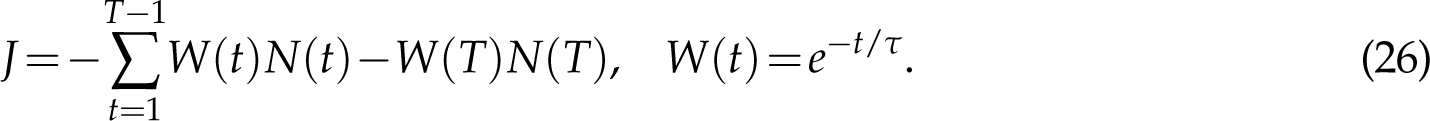

The exponentially-decaying discount *W*(*t*) prioritizes rewards at earlier times and reduces the dependence on *T*, as shown in Fig. S12. We choose *T* = 200 years and *τ* = 50 years. From Eqns. (11–12) we obtain the adjoint equations

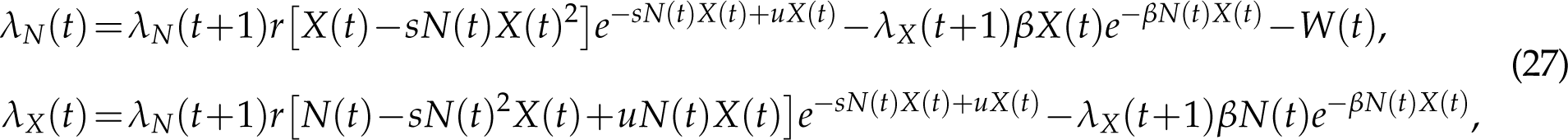

with terminal conditions

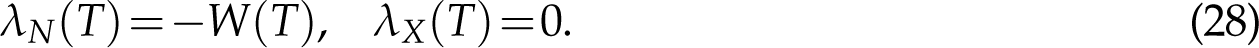

Fig. 7(B) shows *−λ_N_*(*t*), the sensitivity of the reward to moth removal (i.e. culling) at Roseisle for the first 20 years of the time horizon. The sensitivity is positive (i.e. culling is beneficial) near the peak pupae density. However, the maximum sensitivity is not exactly at the peak density, but rather alternates between one year before or after the peak. This alternating offset may be an artifact of the detailed model assumptions and parameter values, but even if real, it still highlights the practical challenge of intervening when sensitivity is highest, because we would need to know which phase of the alternation the system is at, despite measurement uncertainties. On the other hand, if culling is achieved through pesticide spraying, then the demi-elasticity, defined as *−N*(*t*)*λ_N_*(*t*), may be more relevant than the sensitivity if more moths are killed from the same pesticide application when moths are more abundant. In Fig. 7(C), we see that culling is consistently most effective at the peak moth density. This is also mostly true for the Tentsmuir and Culbin sites (Fig. S13).

To understand the values of the demi-elasticities, we consider two scenarios, the first a 20% cull in Roseisle at *t* = 4 (a positive demi-elasticity peak), and the second a 20% cull at *t* = 6 (a negative demi-elasticity valley). By comparing the unperturbed and perturbed trajectories in Fig. 7(D), we see that the increase in reward from a cull at *t* = 4 comes from the immediate reduction in moth density that year; effects in subsequent years are relatively small. (The latter observation is consistent with the observation in Appendix E of Kendall et al. (2005), that pesticide spraying had surprisingly little effect on the dynamics, probably because the outbreaks would have collapsed on their own.) In contrast, the decrease in reward from a cull at *t* = 6 occurs “downstream”: the cull is followed by a large increase in moth density (compared to the unperturbed trajectory) three years later (Fig. 7(E)). For Tentsmuir and Culbin, where the steady-state trajectories are quasiperiodic, immediate and downstream effects can both be large. For example, as shown in Fig. 7(F), the increase in reward from a 20% cull at *t* = 7 (a positive demi-elasticity peak) in Tentsmuir involves not just the immediate reduction in moth density that year, but also the net effect of subsequent years of decrease and increase relative to the unperturbed trajectory. In such situations a robust mechanistic explanation of the downstream changes following a perturbation may not always be possible. To assess whether results are biologically meaningful, a manager should also consider performing TDSA on variants of the model that can still fit the data relatively well, for example using different functional forms for the biological responses. If the demi-elasticities of these variants remain qualitatively similar, a manager can be more confident about using them to guide management actions, based on the idea that “truth is the intersection of independent lies” (Levins, 1966).

#### Larch budmoth

The larch budmoth, *Zeiraphera diniana*, also exhibits large population cycles in parts of Europe. It is believed that both parasitism by wasps and the decrease in tree needle quality after heavy budmoth herbivory play a role in driving the population cycles (Turchin et al., 2003). In addition, outbreaks of budmoths have been found to propagate spatially as recurrent traveling waves across much of Europe (Bjørnstad et al., 2002). To explain these recurrent propagating outbreaks, Johnson et al. (2004, 2006) proposed a tri-trophic (budmoth-plant-parasitoid), spatially-explicit multi-patch model with budmoth and parasitoid dispersal between patches. We performed TDSA on this model, assuming a reward function given by the total plant quality summed across all patches, and across years with exponential time-discount. Details of the model and the adjoint equations can be found in Online Supplement Secs. S6.1 and S6.2. Just as in Johnson et al. (2006), to capture the essential features of the observed recurrent traveling waves, we consider an idealized scenario where suitable budmoth patches are embedded in a larger landscape, with higher patch density towards the center. As shown in Fig. 8(A), the model was indeed able to capture the phenomenon of recurrent traveling waves. Animated maps of the state variables and time-dependent sensitivities can be found in the Online Supplement.

**Figure 8:**
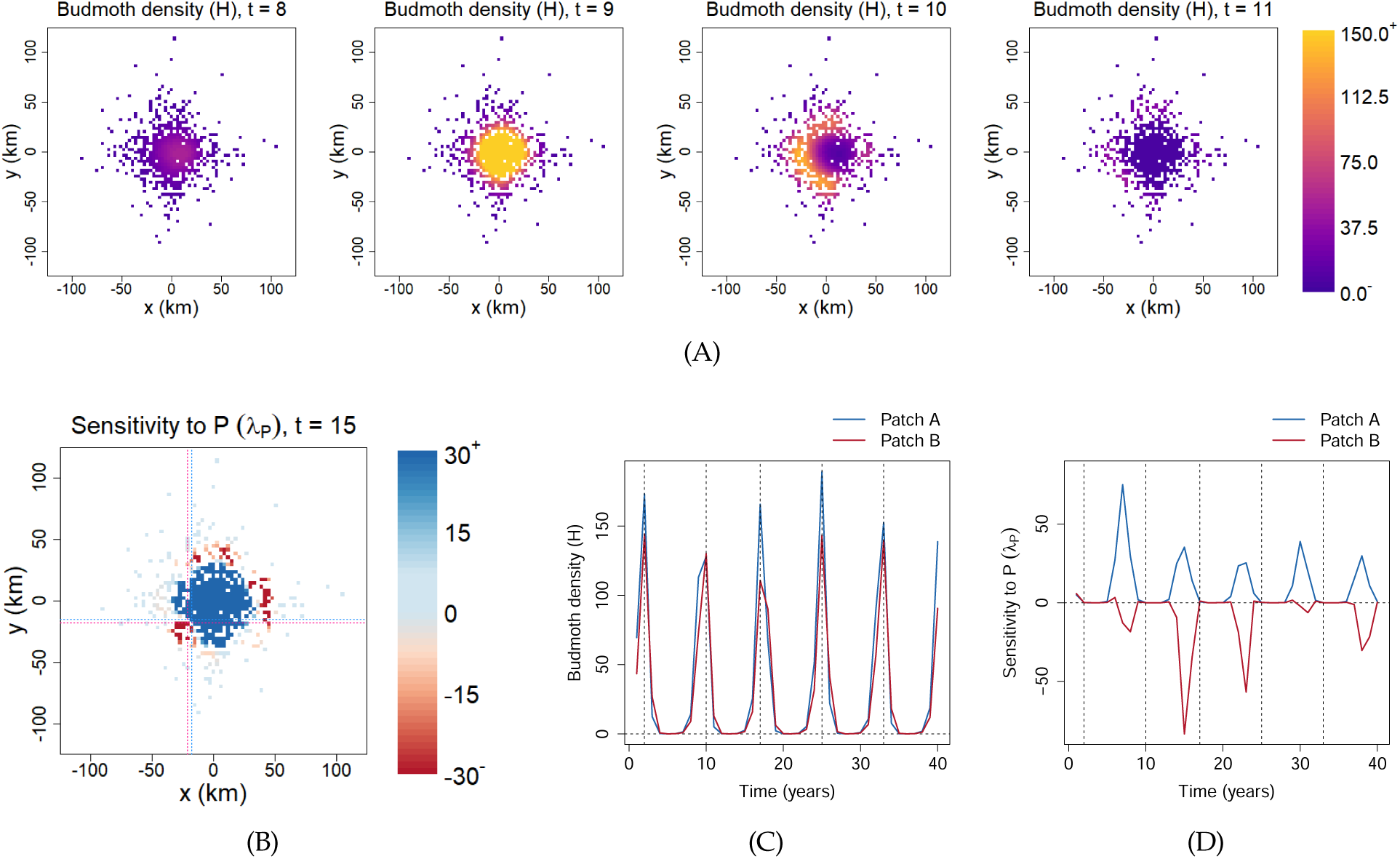
Dynamics and TDSA of the spatially-explicit larch budmoth model. Each colored pixel corresponds to a suitable habitat patch embedded in a larger landscape. Animated maps of the state variables and sensitivities can be found in the Online Supplement. **(A)** Snapshots of the budmoth density, showing radial traveling waves. These propagating outbreaks occur every 7–8 years. **(B)** Snapshot of *λ_P_*, the sensitivity to the addition of parasitoids, at Year 15. We observe a sharp transition from positive to negative values as we move away from the central region. To examine this transition more closely, we selected two adjacent patches, indicated by the intersections of the thin dotted lines (blue–Patch A; red–Patch B). The dynamics of the two patches turn out to be very similar. For example, the budmoth densities peak on the same years as shown in **(C)**. Yet, they show completely different patterns of *λ_P_* in **(D)**: adding parasitoids to Patch A can be beneficial in the right years, but the reverse is true for Patch B. (For reference, the vertical dashed lines indicate the years of peak budmoth densities at the two patches. Also, we have only shown 40 years out of a time horizon of 200 years.) As explained in the main text, this makes it extremely challenging to infer the correct patch-level management actions from the sensitivities alone.

One possible measure to reduce moth populations and improve plant quality is biological control by introducing more parasitoids into a patch; the relevant sensitivity is given by ^⃗^*λ_P_*(*t*). Fig. 8(B) shows the snapshot of ^⃗^*λ_P_*(*t*) at *t* = 15. We observe a sudden transition from positive to negative values as we move radially away from the central region. In other words, adding parasitoids to some of the outer patches actually *reduces* the overall reward.

Rather than attempt a detailed explanation of these results from TDSA, we will instead focus on some qualitative implications of the results. First, because of the large and sometimes abrupt spatial variability in patch-specific sensitivities, biocontrol through parasitoid addition will be more effective when implemented regionally, rather than at the single patch level. Viewing the entire region, it is clear there are large gains from parasitoid addition in a substantial central area, and interventions should be concentrated there. But at the single patch level, management actions may be very difficult to infer. We identified two adjacent patches that have opposite signs in their sensitivities, indicated by the intersections between the thin dotted lines in Fig. 8(B) (blue—Patch A; red—Patch B). We examined the dynamics of the local state variables, and did not notice any qualitative differences. For example, the budmoth densities peak at the same years as shown in Fig. 8(C); Patch B did not lag behind Patch A. Yet, as confirmed in Fig. 8(D), adding parasitoids to Patch A can be beneficial in some years, whereas the reverse is true for Patch B. We verified using explicit perturbations that these adjoint sensitivities were indeed correct (Fig. S14). The mechanism behind the negative sensitivities in Patch B is also not obvious. As shown in Fig. S15, adding parasitoids at *t* = 15 increased plant quality over the next few years in both patches, but only in Patch B did it lead to a larger cumulative decrease in plant quality over the following decades. Because neither the location nor the local dynamics clearly distinguish Patch A from Patch B, how would a manager know in practice whether to add or (if possible) remove parasitoids? Because the location of the transition between positive and negative sensitivities is likely to depend on model details and on parameter values, a manager operating at the single-patch level could find it very challenging to know what local actions are helpful in the long run for the region as a whole.

Inexplicable findings, such as the large differences between Patch A and Patch B, should evoke efforts to determine whether the results are robust, or instead reflect questionable model details. Given the (sadly typical) sparsity of data used to develop and parametrize the budmoth models, it is likely that different choices of functional forms for herbivory, parasitism, competition, and dispersal would fit the data more or less equally well. Would these other models lead to a drastic change in the position of the transition, or cause it to disappear altogether? Even in such a high-dimensional, complex system, TDSA makes it straightforward to get numerical values for the sensitivity of desired outcomes to any state or parameter perturbation; but whether or not to trust those values is an issue that any manager needs to consider. As mentioned in the pine looper example, performing TDSA on multiple variants of the mathematical model is one possible way to assess the robustness of the results.

## Discussion

In this paper, we introduced time-dependent sensitivity analysis (TDSA) as a method for assessing the sensitivity of a system’s dynamics to perturbations in state variables or parameters at any time. Our examples have demonstrated how TDSA can be applied to a wide range of models and applications, where sensitivities vary substantially over time due to environmental variation (e.g., seasonality) and/or transient dynamics. Often, TDSA provides useful insights about the dynamics of the system, some of which would not have been easily discovered without its help. At the same time, Examples 2 and 3 also show why it is important to make an effort to interpret the results and not accept them uncritically, so as to avoid being misled by qualitative features that are really artifacts of specific mathematical assumptions in the input model. Table 1 summarizes our recommended “best practices” to help the TDSA practitioner navigate some of the potential pitfalls.

**Table 1:**
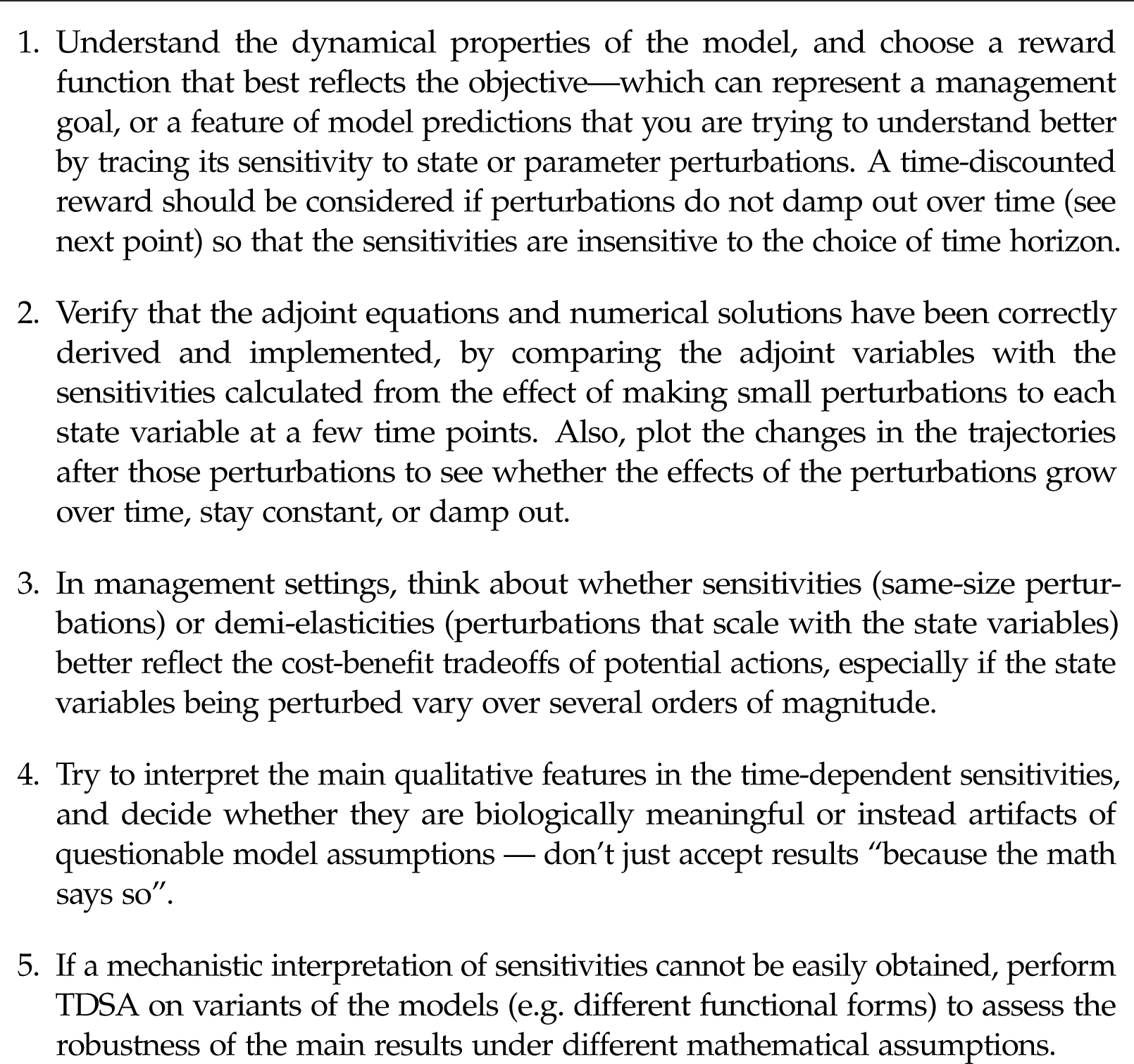
Recommended best practices when performing TDSA.

TDSA can be viewed as a stripped-down version of optimal control theory, which brings both advantages and disadvantages. The disadvantage is that it provides less information, because it is

only guaranteed to be accurate for small perturbations. The main advantage is that it requires fewer assumptions and therefore imposes fewer constraints on the modeler. As in optimal control, TDSA requires formalizing the management goal in the form of the objective function—it forces one to be explicit about exactly what constitutes a desirable outcome. However, an optimal control model must also include a model for the costs of any actions taken. Moreover, cost functions are often chosen in practice so as to satisfy methodological constraints, such as adding a small quadratic term just to satisfy mathematical convexity conditions that imply the existence of an optimal control. As shown in Online Supplement Sec. S2, time-dependent sensitivities can also be made to reflect both costs and benefits, but here only the linearized costs matter. Finally, time-dependent sensitivity analysis is also computationally much simpler and faster, requiring only straightforward numerical solution of the state and adjoint equations, rather than iterative solution of those plus the first-order optimality condition. The method of adjoint sensitivity analysis (ASA) which we used is not the only way of doing TDSA. An alternative, more direct method is forward sensitivity analysis (FSA) (e.g., Cacuci et al., 2003), which uses the variational equations of the state dynamics to calculate how a small change to each parameter affects state trajectories and the reward function. However, FSA requires a new solution of the variational equations for each parameter and each time of perturbation, so whenever one is interested in the effect of perturbations at many different time points, ASA will be far more efficient.

As we have demonstrated, for management purposes, it is sometimes more meaningful to convert sensitivities to demi-elasticities. For example, a high-sensitivity species may be very rare, making management actions targeting the species impractical. As shown in the pine looper example, demi-elasticities also better reflect the costs and benefits of management actions such as pesticide application, where the number of pests directly killed scales with the pest density. In addition, we recommend performing checks to confirm that TDSA has been correctly implemented, e.g. by comparing the adjoint variables to sensitivities calculated from explicit perturbations at a number of time points. Although we have performed the comparisons for *all* time points in Figs. S9 and S14, this is not necessary and may not even be computationally feasible for high-dimensional models (the very motivation behind the use of the adjoint formalism).

Although TDSA is mathematically rigorous, we advise against blind acceptance of its results, especially if they are to inform management actions. The results from TDSA ultimately depend on the choice of mathematical model used to describe the dynamical system. Hence, the practitioner should make an effort to interpret key features in the sensitivity, and decide whether they only rely on the biology broadly conceptualized by the model, or on specific mathematical assumptions of the model chosen for simplicity. We saw both types of features in the chytrid fungus example, and we explained why only the former should be used for assessing management decisions. If a mechanistic explanation is not possible, like in the pine looper example, the practitioner should consider performing TDSA for multiple variants of the model to assess the robustness of the results. In addition, the dynamics of the system matters. If quasiperiodic, the effects of a perturbation may persist without damping (e.g. Fig. S11(c) for the site Tentsmuir in the pine looper example); this will need to be considered when defining the reward function so that the sensitivities at early times do not depend too much on the time horizon. If model dynamics are chaotic, results from TDSA may be difficult to interpret (although this would also be true for other forms of sensitivity analysis).

We have shown how TDSA can be applied to a wide variety of deterministic models, including models with continuous independent variables through discretisation. Although we did not demonstrate this using an example, TDSA should also work for models with distributed time delays, e.g. by using the (generalized) “linear chain trick” (Hurtado and Richards, 2020; Hurtado and Kirosingh, 2019) to convert them into differential equation models, or by formulating the models as integro-differential equations with age classes (which can then be discretized). On the other hand, extending TDSA to stochastic systems is potentially challenging because the impacts of a perturbation will vary between different random realizations of the dynamics.

We live in a time-varying world, where knowing when to act is often just as important as knowing how to act. TDSA simultaneously addresses both questions, and offers a systematic way of probing the dynamics of a model, thereby enhancing our understanding of the biological system and facilitating decisions on how to achieve management goals. By presenting a balanced view that highlights both the strengths of TDSA as well the potential pitfalls, we hope that TDSA can become a useful addition to the toolkit of the modelers and natural resource managers.

## Manuscript elements

Figures 1–8, Table 1, Appendices A–B. Online Supplement includes Sections S1–S6, Figures S1–S15, Table S1, and an animated GIF file.

## Data and code accessibility

No original data are presented in this paper. All supporting computer scripts are included with the submission, and will be archived on Zenodo and the DOI added to the final manuscript if accepted.

## Acknowledgements

*Acknowledgments*: We thank Megan Greischar, Christina Hernández, Timothy Lambert, Martina Morelli, Anna Poulton and Andrew Siefert for helpful comments. Research reported in this publication was supported by the National Institute of General Medical Sciences of the National Institutes of Health under Award Number R01GM122062, as well as the USDA National Institute of Food and Agriculture under the Ecology and Evolution of Infectious Diseases grant no. 2021-67015-35235. Any opinions, findings, conclusions, or recommendations expressed in this publication are those of the author(s) and do not necessarily represent official views of the National Institutes of Health nor the U.S. Department of Agriculture.

NOTE: for the convenience of reviewers, figures and tables have been placed with their captions close to where they are referred to in the text. If the paper is accepted for publication, the final manuscript will be formatted in accordance with the Instructions for Authors.

Prepared using the suggested LAT_E_X template for *Am. Nat*.

## Appendix A: Derivation of the adjoint equations and terminal conditions

In this section, we present a modified version of the proof from Kamien and Schwartz (1991), Part II, Section 4, that the time-dependent sensitivity satisfies the adjoint equations and terminal conditions given in Eqn. (5). When we perturb the state vector at time *t*, only the contribution to the reward *J* downstream from the perturbation will be affected. Hence we introduce the value function

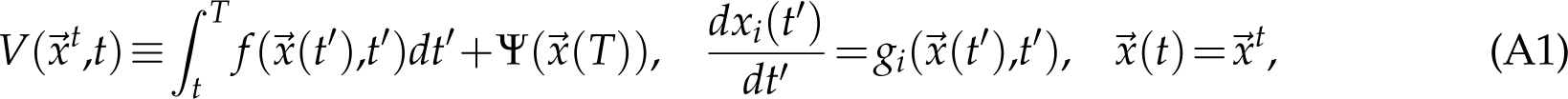

which gives the total contribution to *J* from time *t* to *T*, when the state vector is equal to *⃗x^t^* at time *t*. For the original unperturbed trajectory, which we will denote as *⃗x^∗^*(*·*) to avoid confusion, *⃗x^t^*=*⃗x^∗^*(*t*), but we will also consider other values of *⃗x^t^*, in which case the subsequent trajectory *⃗x*(*·*) will not be *⃗x^∗^*(*·*). (Think of the argument *⃗x^t^* as specifying the “initial conditions” at time *t*.) The value function is useful because if we perturb the original state vector to *⃗x^t^* at time *t*, the change in reward is then given by the difference Δ*J* = *V ⃗x^t^,t −V*(*⃗x^∗^*(*t*),*t*).

We now re-write the value function in a different form. First, we introduce a function ^⃗^*λ*(*·*) that is as of now arbitrary. From the definition in Eqn. (A1),

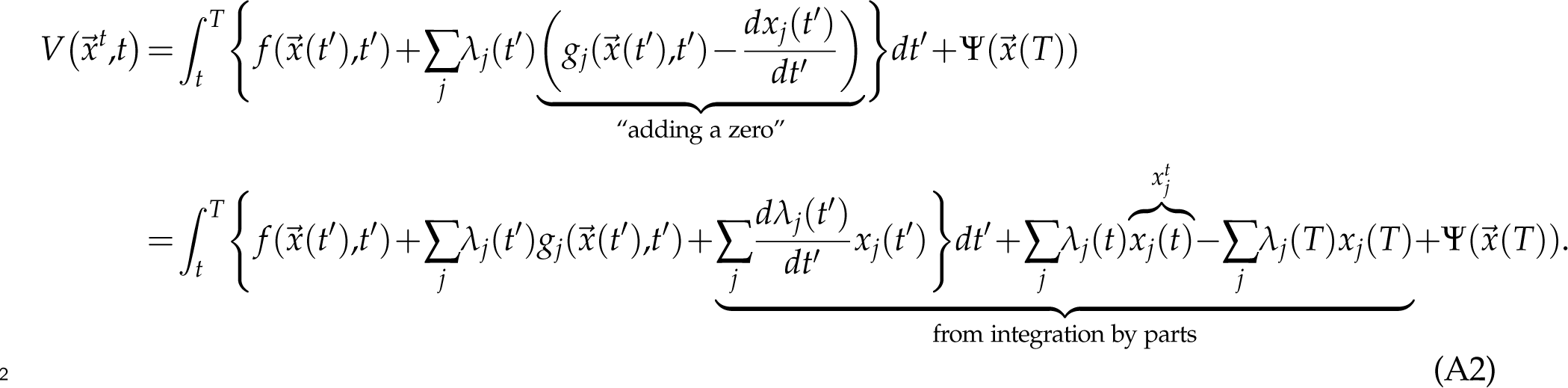

The change in reward can then be written as

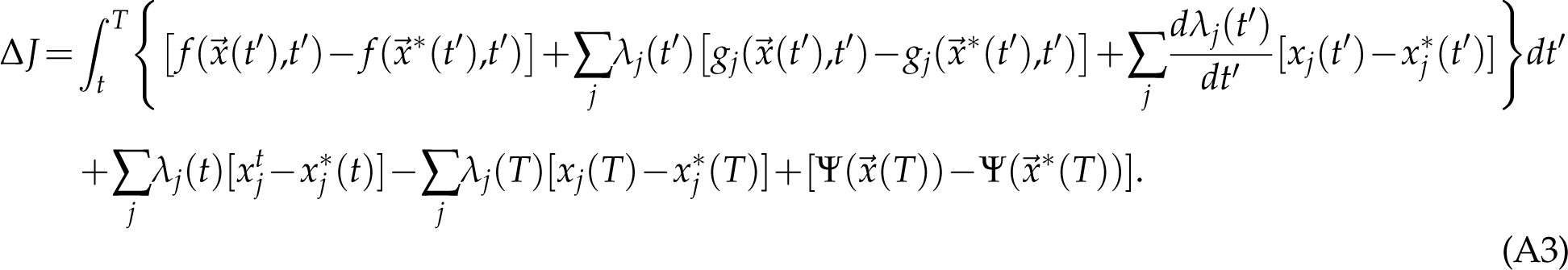

Now say we only perturb the *i*th state variable by an amount *ɛ* at time *t*, so 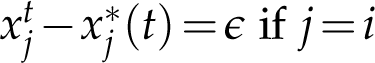 if *j* = *i* and 0 otherwise. From Taylor approximation, Eqn. (A3) becomes

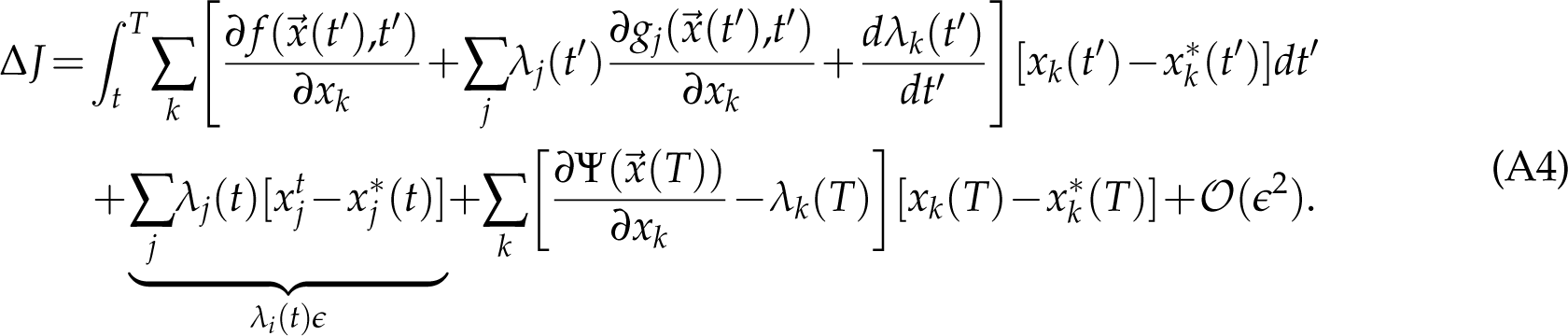

Notice that if we now choose the arbitrary function ^⃗^*λ* to satisfy the adjoint system Eqn. (5), the terms in large square brackets vanish, leaving

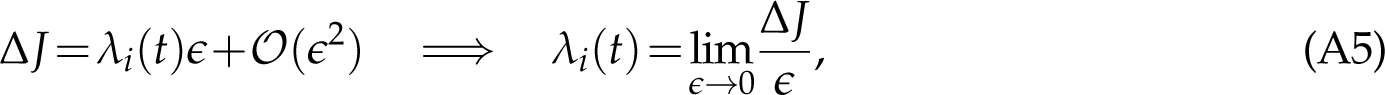

so *λ_i_*(*t*) is just the sensitivity to the above state perturbation. In other words, if a function ^⃗^*λ* satisfies the adjoint system, then it can be interpreted as a time-dependent sensitivity. Since the sensitivity is single-valued, this means that the converse must be true, that the sensitivity must satisfy the adjoint system. This completes the proof.

## Appendix B: Time-dependent parameter sensitivity

In this section, we derive Eqn. (A9), a formula that can be used to calculate time-dependent parameter sensitivities from the adjoint variables. Consider a parameter perturbation of the for^m⃗^*θ ^→^*^⃗^*θ*+*^ɛ^*^⃗^*h*, where ^⃗^*h* is a vector-valued function of time that indicates the relative size of perturbation in each parameter (so it will only have one non-zero component if we only perturb a single parameter), as well as the temporal pattern of the perturbation (so it will only be nonzero over a short time interval if we only perform a brief perturbation). *ɛ* is a small parameter that represents the size of perturbation. From Eqn. (1), the resulting changes in the state variables *⃗x →⃗x*+*δ⃗x* satisfy the following (linearized) dynamic equation:

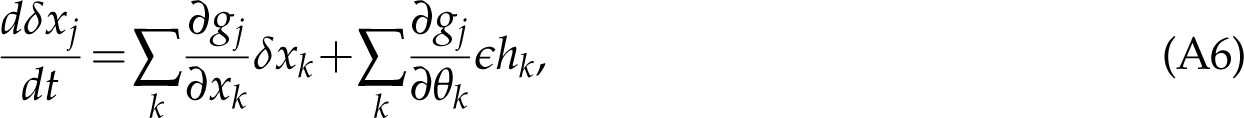

and *δ⃗x* in turn changes the reward function, Eq. (2), by

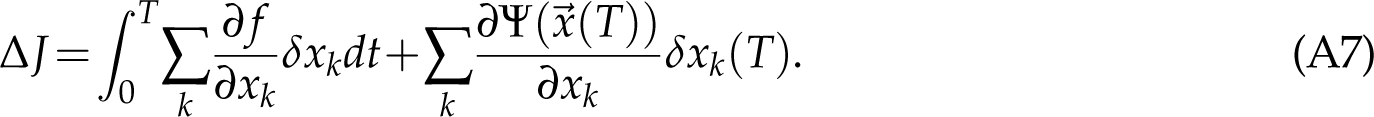

Plugging Eqns. (5–6) and (A6) into the above expression of Δ*J*, we get

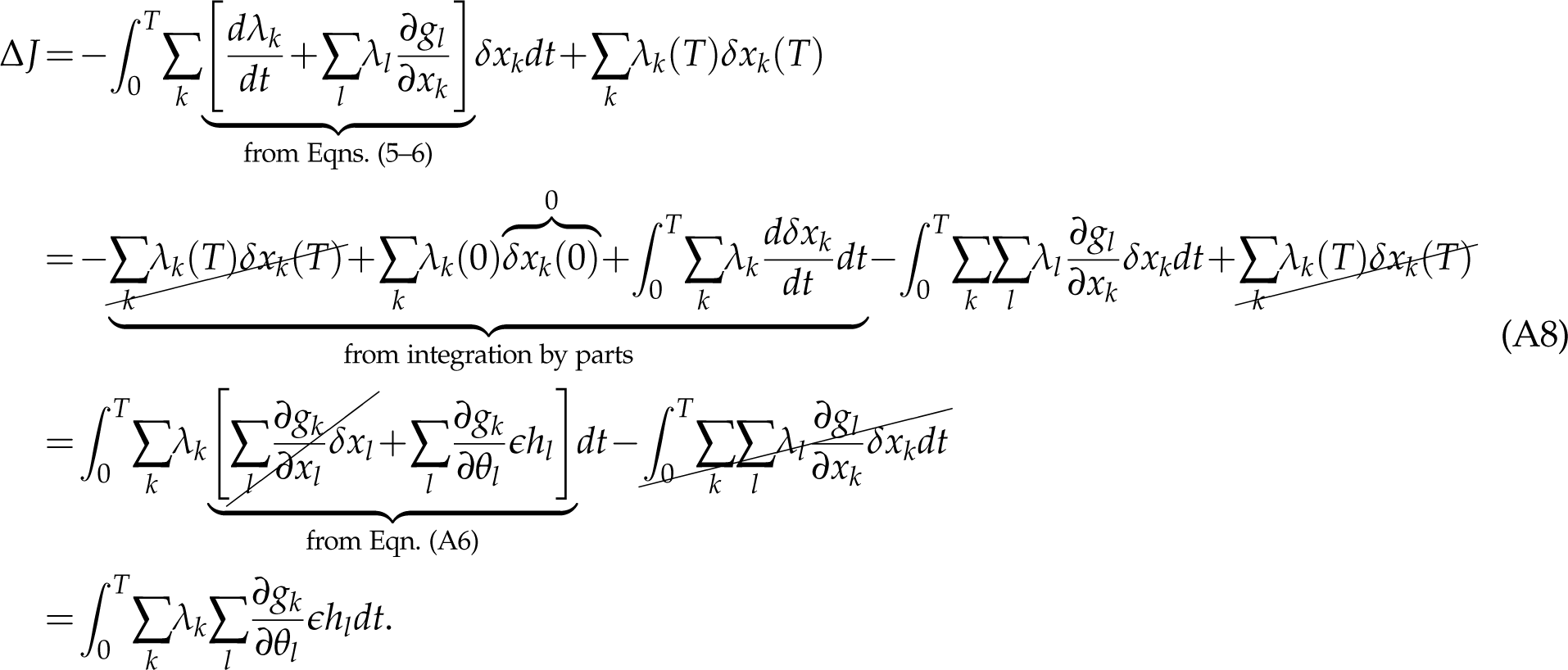

Note that in the second step, *δx_k_*(0) = 0 since a finite parameter perturbation starting at *t* = 0 should not cause a finite change in the state variables at *t* = 0. The sensitivity (a Gateaux derivative, in the language of functional analysis) is therefore given by

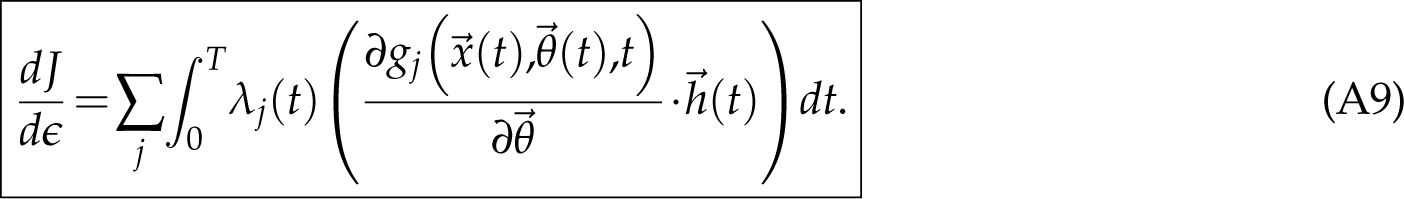

where 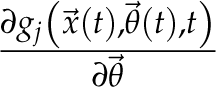 is the vector 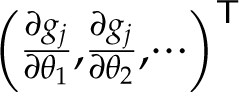.

Since the normalization of ^⃗^*h* affects the value of the sensitivity, if we are trying to compare perturbations associated with different management options, it is preferable that we normaliz^e⃗^*h* for each option in a way that permits a fair comparison. For example, if *^ɛ^*^⃗^*h* is a brief perturbation centered at time *t^∗^* only in the *k*th component o^f⃗^*θ*, and we normalize *h_k_* such that 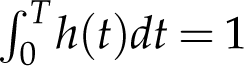, then Eqn. (A9) reduces to

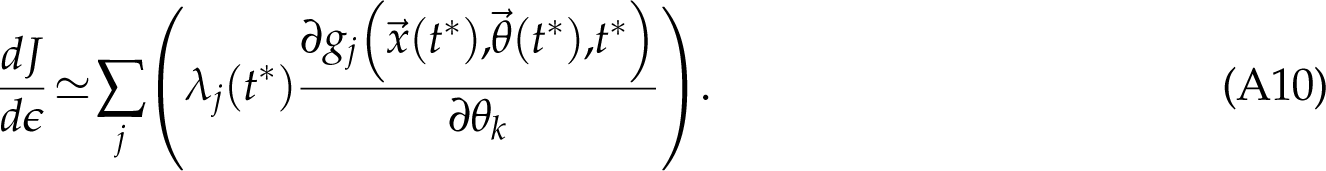

Eqn. (A10) also provides some insights into the interpretation of the more general expression, Eqn. (A9). Comparing the two equations, we see that the integral in Eqn. (A9) can be thought of as “chopping” up a more general *^ɛ^*^⃗^*h* into a series of brief perturbations centered at different times, and then summing over the sensitivities to these brief perturbations.

## Online Supplement

Ng et al, A time for every purpose: using time-dependent sensitivity analysis to help manage and understand dynamic ecological systems, *The American Naturalist*.

## S1 Parameter values for the introductory model

In this section, we provide the parameter values of the introductory model Eqns. (3) and (4) used to illustrate the adjoint method. As a reminder, the model describes a population in a sink habitat that is currently maintained through immigration, but the habitat is being restored so eventually the population will become self-sustaining. We use the abbreviation PU for the arbitrary population unit, and VU for the arbitrary value unit.

- Unregulated per-capita birth rate: We choose *b* = 1/year.
- Per-capita loss rate: We want *µ*(*t*) to decrease as a sigmoid, so we choose

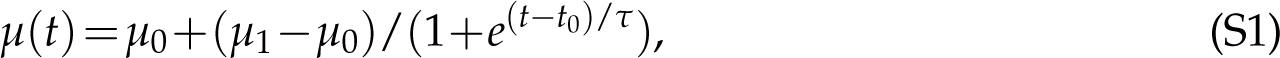

where *µ*_0_ = 1.5/year and *µ*_1_ = 0.5/year are the pre- and post-restoration per-capita loss rates, *t*_0_ = 10 years the time at the inflection point of the sigmoid, and *τ* = 2 years a timescale that characterises the steepness of the sigmoid.
- Coefficient for intraspecific competition: We choose *a* = 0.1/PU
- Immigration rate: We choose *σ* = 0.2 PU/year.
- Per-capita rate of contribution to ecosystem service: We choose *w* = 1 VU/year/PU.
- Per-capital terminal payoff: In this example, any perturbation will eventually decay downstream, so it is possible to eliminate the effects of a finite time horizon if we choose *v* such that it is equal to the ecosystem service contribution had the time horizon been extended indefinitely beyond *T*. To estimate this, we linearise Eqn. (3) about the post-restoration carrying capacity *K*, and find that any perturbation will decay exponentially at a rate *µ*_1_*−b*(1*−*2*aK*) and hence contribute a reward of *w*/[*µ*_1_*−b*(1*−*2*aK*)]. Based on this reward, we choose *v* = 1.74 VU/PU.
- Initial conditions: We want *x*(0) to be the steady-state population pre-restoration. Solving the equation *bx*(0)(1*−ax*(0))*−µ*_0_*x*(0)+*σ* = 0 gives us *x*(0) = 0.37 PU.

## S2 Incorporating perturbation costs into time-dependent sensitivities

Just like in optimal control theory, we now consider a manipulated system

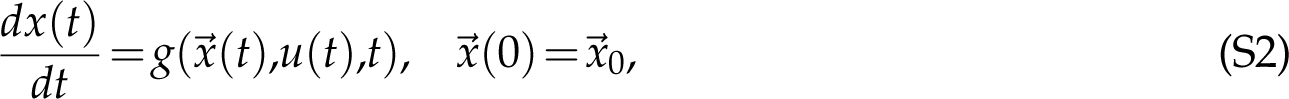

where *u*(*t*) quantifies the external manipulation. We also define

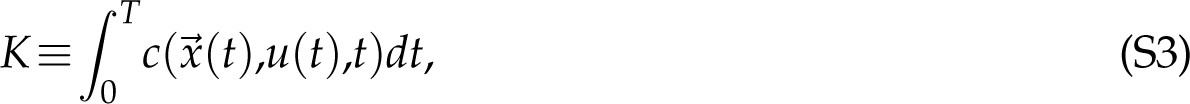

the total cost of the manipulation, analogous to the total reward function *J*. If there is no manipulation, there is no manipulation cost, so we require that *c*(*⃗x*,0,*t*) = 0 for any *⃗x* and *t*. At the same time, we assume that the integrand *f* (*⃗x*(*t*),*t*) of the total reward *J* does not depend directly on *u*(*t*).

We are interested in the effects of a small, brief manipulation at time *t^∗^* on the net value *J−K*. More specifically, we consider *u* = *ɛh*, where *h* is a narrow window function centered at time *t^∗^*, normalized such that 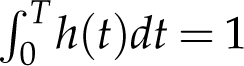. Since *J* is only indirectly affected by the manipulation through the effects on *⃗x*(*t*), if we interpret *u* as yet another parameter with an unperturbed value of 0, we can apply Eqn. (A10) from Appendix B, so

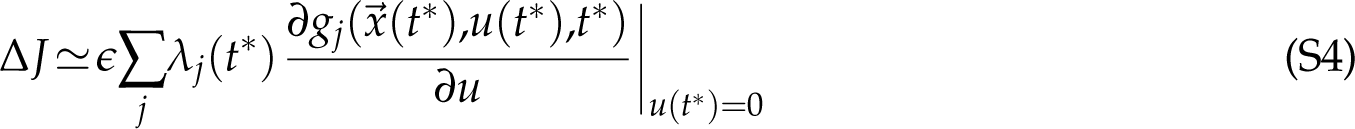

Meanwhile, since *c*(*⃗x*,0,*t*) = 0 for any *⃗x* and *t*, this is also true for its partial derivative in *⃗x*, so to order *O*(*ɛ*), Δ*K* only comes from the direct dependence of *c* on *u*. More specifically,

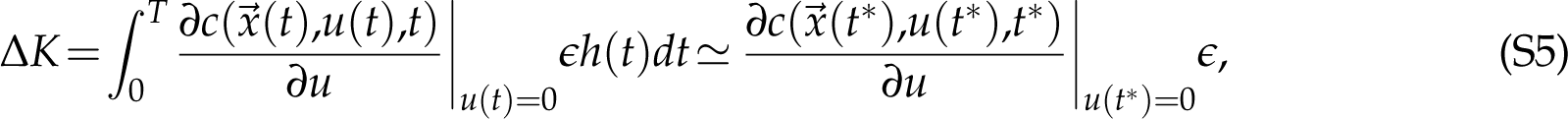

where in the second step, we used the fact that *h* is a normalized narrow window function centered at time *t^∗^*. Hence, the sensitivity to a small, brief manipulation at time *t^∗^* is given by

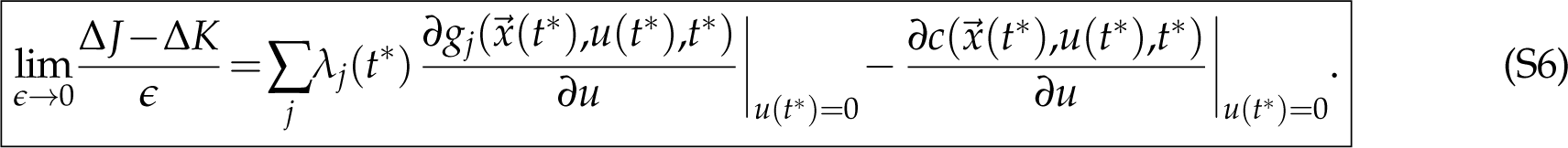

Note that unlike optimal control theory, we only need the linearized versions of the functions *g_j_* and *c* about *u* = 0 and not their full functional forms in order to calculate the sensitivity.

### S3 Change of adjoint variables under a change of state variables

Let *⃗x* be the original state variables, and *⃗y* be the new state variables. For simplicity, assume that the transformation is invertible and also has no explicit time dependence, so we can write each new variable *y_i_* as a function *y_i_*(*⃗x*) of the old variables, and each old variable as a function *x_i_*(*⃗y*) of the new variables. When taking partial derivatives, it is important to keep track of what other variables are being held constant. We will use the notation 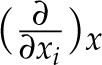 to mean holding all other *x_j̸_*_=_*_i_* constant. The old and new variables satisfy the dynamic equations

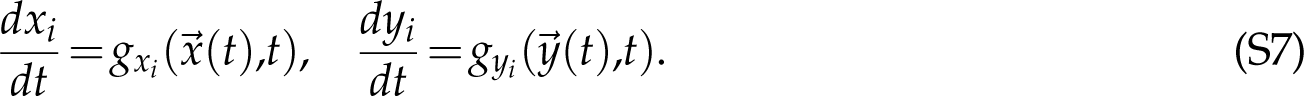

Since the transformation does not contain any explicit time dependence, chain rule tells us that

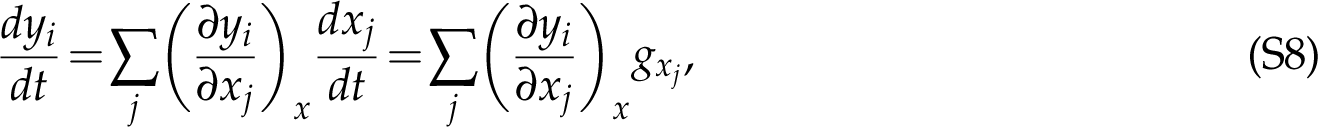

so we have the relation and inverse relation

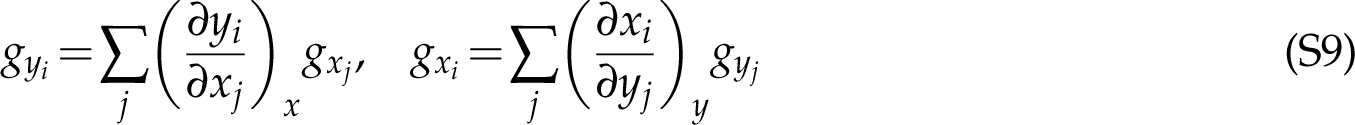

Let the reward function be

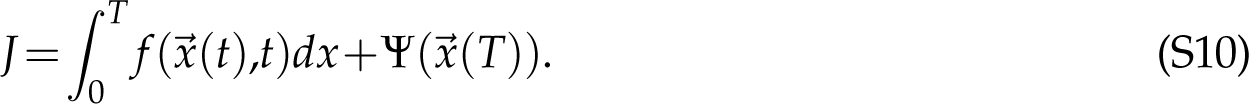

The old adjoint variables satisfy the adjoint equations and terminal conditions

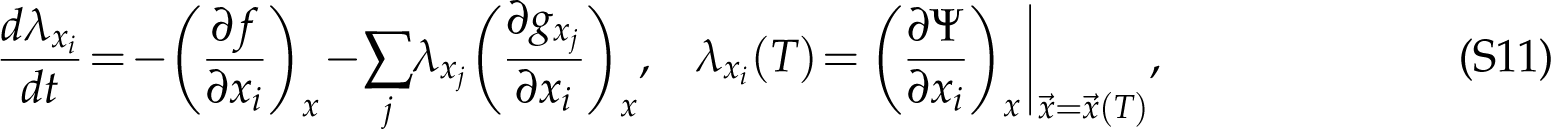

while the new adjoint variables satisfy

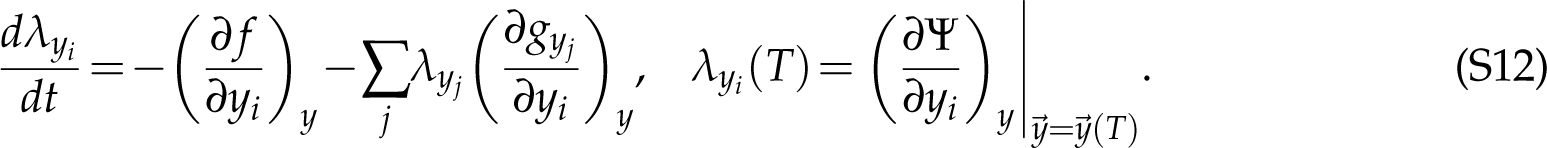

In the remainder of this section, we will prove the relation

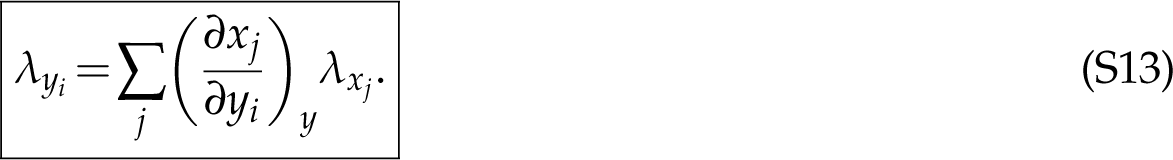

First, we define

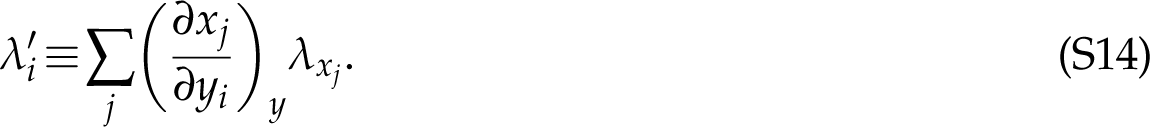

Our strategy is to show that *λ^′^* satisfies the same adjoint equations and terminal conditions as *λ_yi_*, so we can then conclude that *λ^′^* = *λ_yi_*, hence proving the relation. Consider

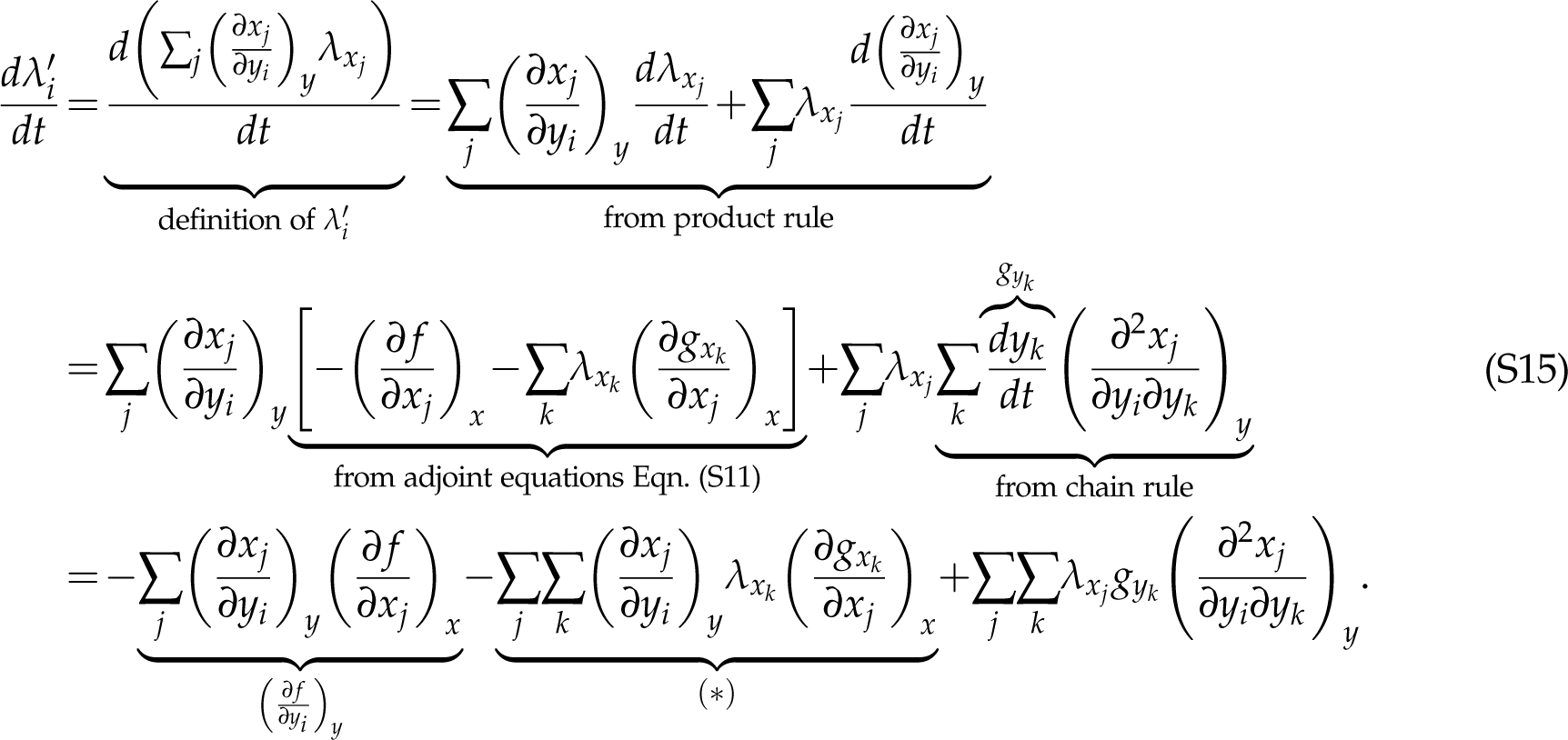

We will first simplify the term (*∗*) before returning to the equation. We have

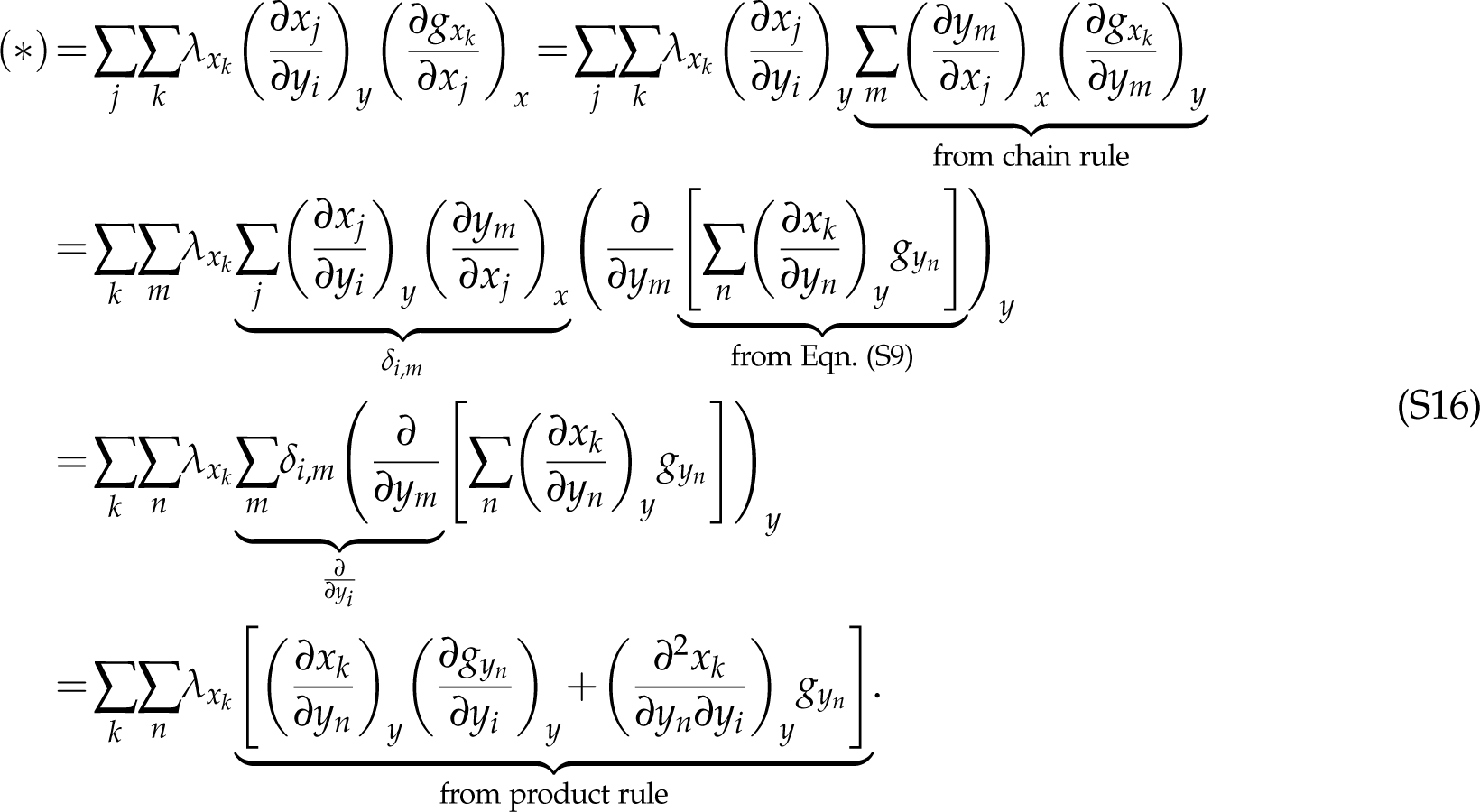

Now we replace the dummy variables *k* and *n* in (*∗*) by *j* and *k* respectively, and plug it back into Eqn. (S15). We get

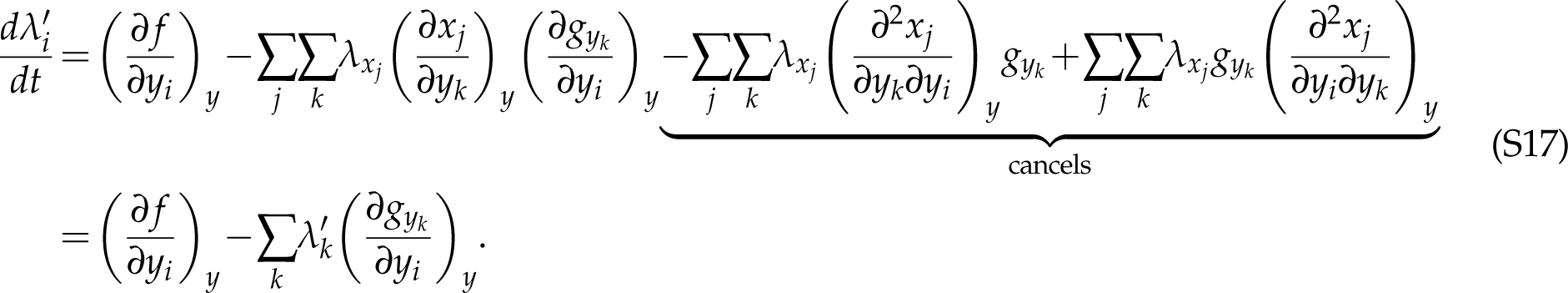

Comparing Eqn. (S17) to Eqn. (S12), we see that *λ^′^* does indeed satisfy the same adjoint equations in Eqn. (S12) as *λ_yi_*. All that is left is to show that *λ^′^* also satisfy the same terminal conditions in Eqn. (S12).

Consider

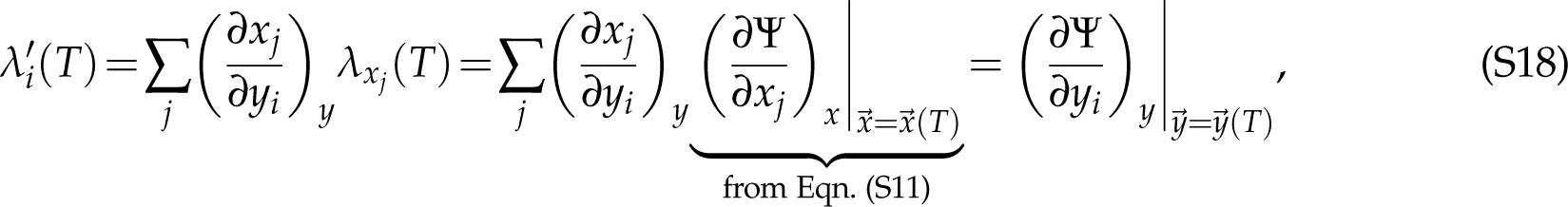

hence completing the proof.

More elegant proofs probably exist from optimal control theory, but this version is the most straightforward.

## S4 Parameter values for Example 1: Disease spillover into multi-species sink communities

As mentioned in the main text, the parameter values have been chosen to best illustrate the qualitative features of interest. We explain the choices in more details below.

- Disease-free mortality (*µ_j_*): For simplicity, we assume that all species have the same *µ_j_*. Without loss of generality, we choose the units of time so that one unit corresponds to one lifespan, so *µ_j_* = 1 for all *j*.
- Unregulated per-capita birth rate (*B_j_*): For the species of concern, we want there to be a substantial population decline despite the low infection prevalence (especially if the disease reaches the species of concern from the exogenous source only after a long chain of transmission), so that control measures are necessary. Therefore, we choose *B_jC_*= 1.02 so that it is only very slightly above *µ_jC_*. For all other species, as explained in the main text, culling an intermediate species too early in the season is ineffective since the population would have mostly recovered by the time the chain of infection reaches the species. To demonstrate this point clearly, we want *B_j_ ≫µ_j_*, so we choose *B_j_* = 5.
- Intraspecific competition coefficient (*a_j_*) or carrying capacity (*K_j_*): We can specify either *a_j_* or *K_j_* since they are related by *K_j_* =(1*−µ_j_*/*B_j_*)/*a_j_*. For simplicity, we assume that all species have the same *K_j_*, and without loss of generality, we choose the units of population size so that *K_j_* = 1 for all *j*. This means that *a_j_* = 0.8 for all species, except the species of concern, where *a_jC_≃* 0.02. In other words, the large carrying capacity in the species of concern despite the low birth rate is due to low intraspecific competition. Alternatively, we could have chosen the same competition coefficient *a_j_* = 0.8 for all *j*, in which case all species will have *K_j_* = 1 except for the species of concern, where *K_jC_≃* 0.02, i.e. a low carrying capacity. We find that most qualitative features observed in the two networks are still present under this alternative scenario.
- Disease-induced mortality (*ν_j_*): We want a large disease-induced mortality in the species of concern, so we choose *ν_jC_*= 5. In contrast, for all other species, we choose *ν_j_* = 0, so the disease has no impact on their populations.
- Recovery rate (*γ_j_*): Again, for there to be a substantial population decline in the species of concern, we need a high per-capita rate of infection in the species of concern, even after a long chain of transmission, while still keeping *R*_0_ <1. Numerically, we find that this is easiest to achieve when all species have comparable infectious lifetimes 1/(*µ_j_* +*ν_j_* +*γ_j_*). Since the species of concern already has a short infectious lifetime due to the large disease-induced mortality *ν_jC_*, we set *γ_jC_*= 0. For all other species without disease-induced mortality, we choose *γ_j_* = 5, so that they recover quickly from infection.
- Length of active season (*T*): Even though both networks were meant to be hypothetical, we designed them with pollinators in mind. Since the average lifespan of a bee is of order 20–30 days, we choose *T* = 5 so that the active season would correspond to a realistic period of 100–150 days.
- Coefficients in the reward function (*W_SjC_*, *W_IjC_*, *V_SjC_*, *V_IjC_*): Without loss of generality, we choose the units of value so that *W_SjC_*= 1. We assume that infected individuals are just as capable of providing the ecosystem service, so *W_IjC_*= 1 as well. (One possible scenario is that most infected individuals in the species of concern start off as asymptomatic carriers, but quickly die once the symptoms set in. Therefore, the fecundity of infected individuals as well as the ecosystem service they provide remain unaffected before they die.) For the terminal payoffs, we arbitrarily choose *V_SjC_*= *V_IjC_*= 1. We find that most qualitative features observed in the networks are still present under other choices of *W_IjC_*, *V_SjC_* and *V_IjC_*.
- Transmission coefficients (*b_j_*_,*k*_): We parametrize *b_j_*_,*k*_ according to the network structure and then rescale them so that the dominant eigenvalue of the next-generation matrix is *R*_0_. Below, we present the values of *b_j_*_,*k*_ *before* rescaling.

**–** Network 1: We take the *c→* ∞ limit of the trait-matching model, which gives

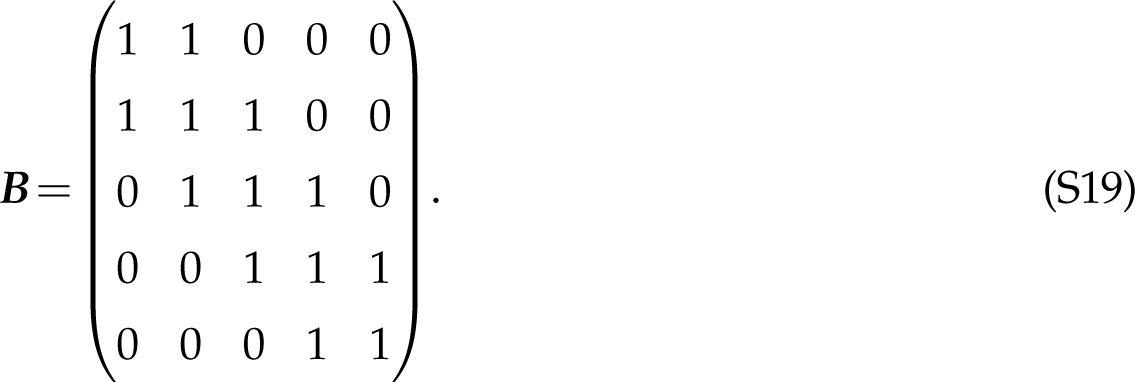

**–** Network 2: We first define resource utilization *r_j_*_,*k*_ as the relative frequency an individual of species *k* chooses to utilize resource type *j*. As explained in the main text, there are two resource types, and bridge species 3 (the species of concern) is less specialized, so we choose

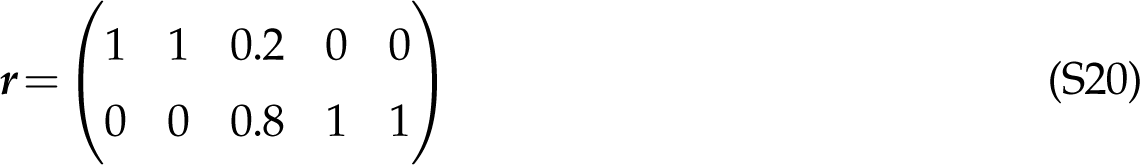

We then assume that ***B*** is given by ***B*** = ***r****^T^**r***. To enhance intraspecific transmission in species 5, we also double the value of *b*_5,5_.

- Basic reproduction number (*R*_0_): We choose *R*_0_ = 0.9 for Network 1, and *R*_0_ = 0.95 for Network 2.
- Spillover coefficient (*σ_j_*): In both networks, only the first species receive exogenous spillover. We choose *σ*_1_ = 0.2 for both networks.
- Initial conditions (*S_j_*(0), *I_j_*(0)): We choose *S_j_*(0) = *K_j_* and *I_j_*(0) = 0 for all *j*. In other words, we assume that each species starts the current season disease-free at the carrying capacity. This is mainly for simplicity, so that the transient dynamics mostly reflect disease transmission and not population growth.

## S5 More details on Example 2: Leopard frogs as reservoirs of the amphibian chytrid fungus

### S5.1 Functional forms and parameter values

The load-dependent functions ℓ(*x*), *G*_0_(*x*) and *G*(*x^′^|x*) are assumed to take the form

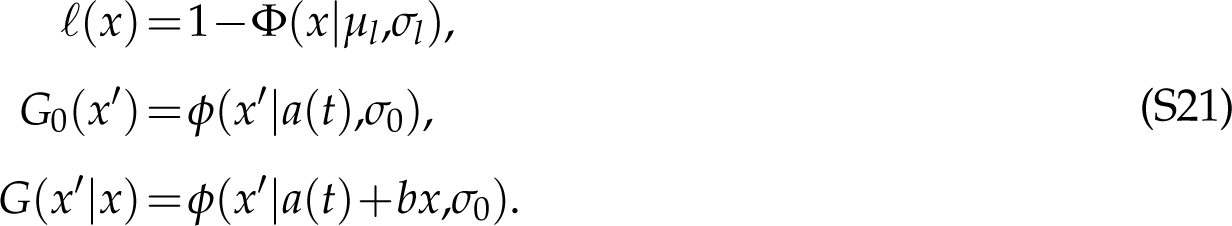

Here *ϕ* and Φ are the probability density and cumulative distribution functions of the normal distribution, with mean and standard deviation given by the two parameters after the vertical bars.

The temperature-dependent functions *a*(*T*) and *s_Z_*(*T*) are assumed to take the form

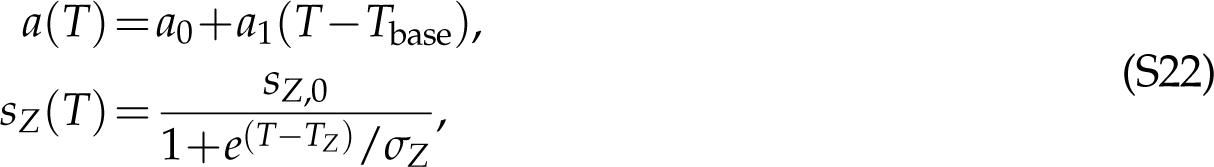

The temperature is assumed to vary sinusoidally across the year, and is given by

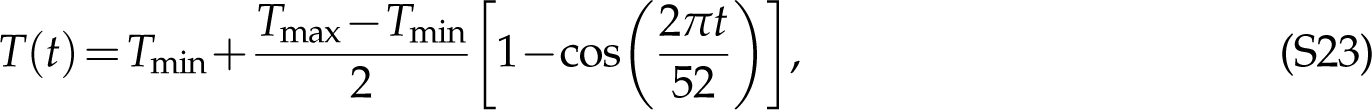

where *t* here is in weeks, and it is assumed that one year has exactly 52 weeks.

Wilber et al. (2022) fitted separate *Bd* transmission models at four geographic locations (Louisiana, Tennessee, Pennsylvania, and Vermont), and at three possible values of the parameter *K* controlling density dependence in recruitment: *e*^10^ (low density), *e*^8^ (medium density) and *e*^4^ (high density). Most parameter values can be found in Table S2 from Wilber et al. (2022); we chose parameter values for Tennessee under the high-density assumption, as well as *s_I_* = 1. Other parameter values that can only be found in the main text or in their scripts are: *T*_min_ = 4*^◦^*C, *T*_max_ = 27*^◦^*C, aquatic calendar days 30–150 (so *W*(*t*) = 1 for week numbers 5–21), and reproduction calendar day 90 (so *R*(*t*) = 1 for week number 13).

### S5.2 Discretizing the IPM

We discretize the IPM in Eqn. (20) into *m* bins each of width *h*. The *i*th bin has midpoint *x_i_*, lower and upper boundaries *<u>x</u>_i_* and *x_i_*, and contains *I_i_*(*t*) infected individuals (so *I_i_*(*t*) approximates *I*(*x_i_*,*t*)*h*). The discretized equations are then given by

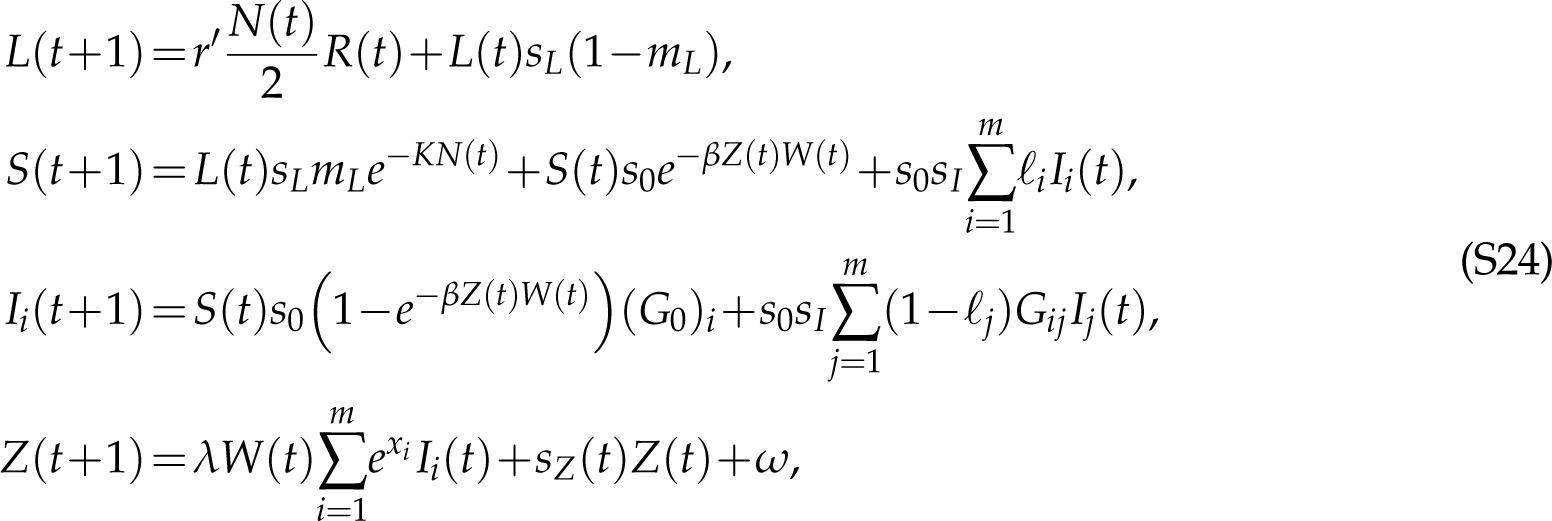

where

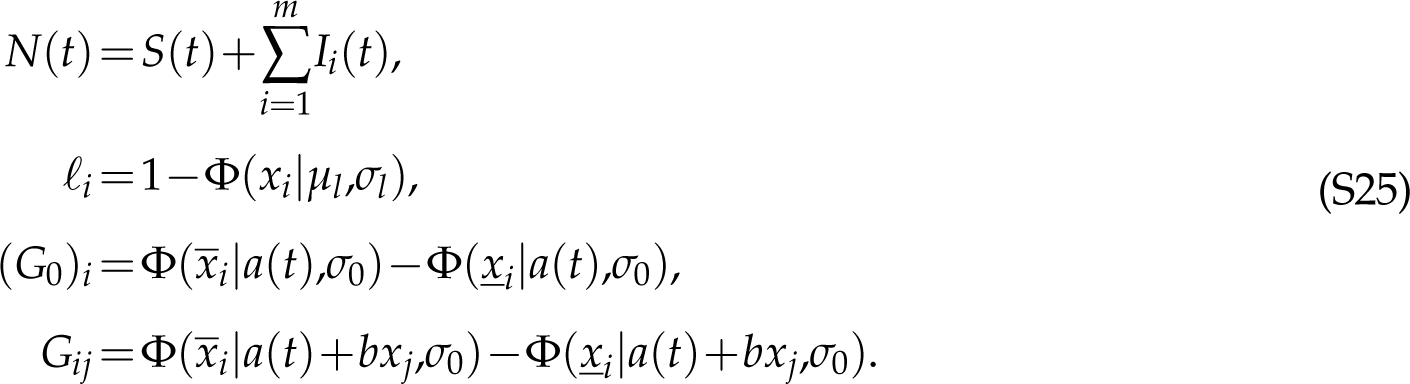

### S5.3 Deriving the adjoint equations

To derive the adjoint equations, we first write down the Hamiltonian

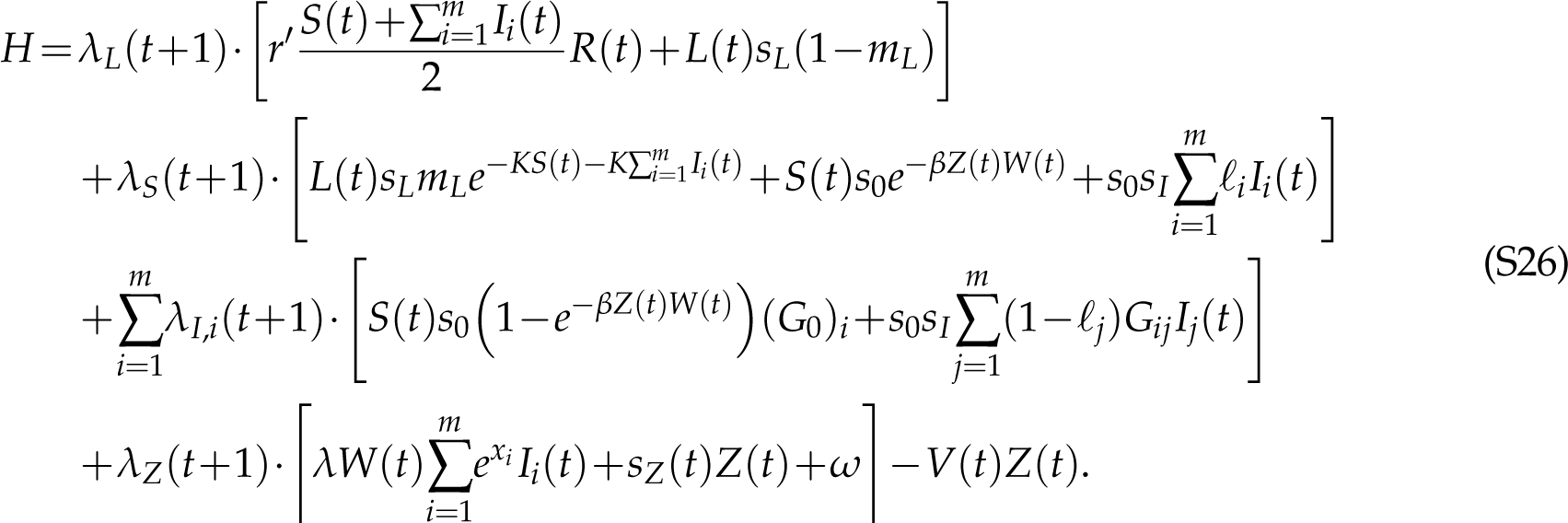

We then obtain the adjoint equations, Eqn. (22), by taking partial derivatives of the Hamiltonian *H* according to Eqn. (11).

## S6 More details on Example 3: **Population cycles in the pine looper and the larch budmoth**

### S6.1 Larch budmoth: Model details

Johnson et al. (2004, 2006) proposed a tritrophic, spatially-explicit, discrete-time model, where budmoths and their parasitoids are located in patches of suitable habitats embedded within a larger landscape. In each patch, which we index by *i* (maximum *n*), and at year *t*, the local densities of budmoths and parasitoids are represented by state variables *H*(*i*,*t*) and *P*(*i*,*t*), while the local plant quality is represented by the state variable *Q*(*i*,*t*) with a maximum value of 1. The dynamics can be represented by the equations

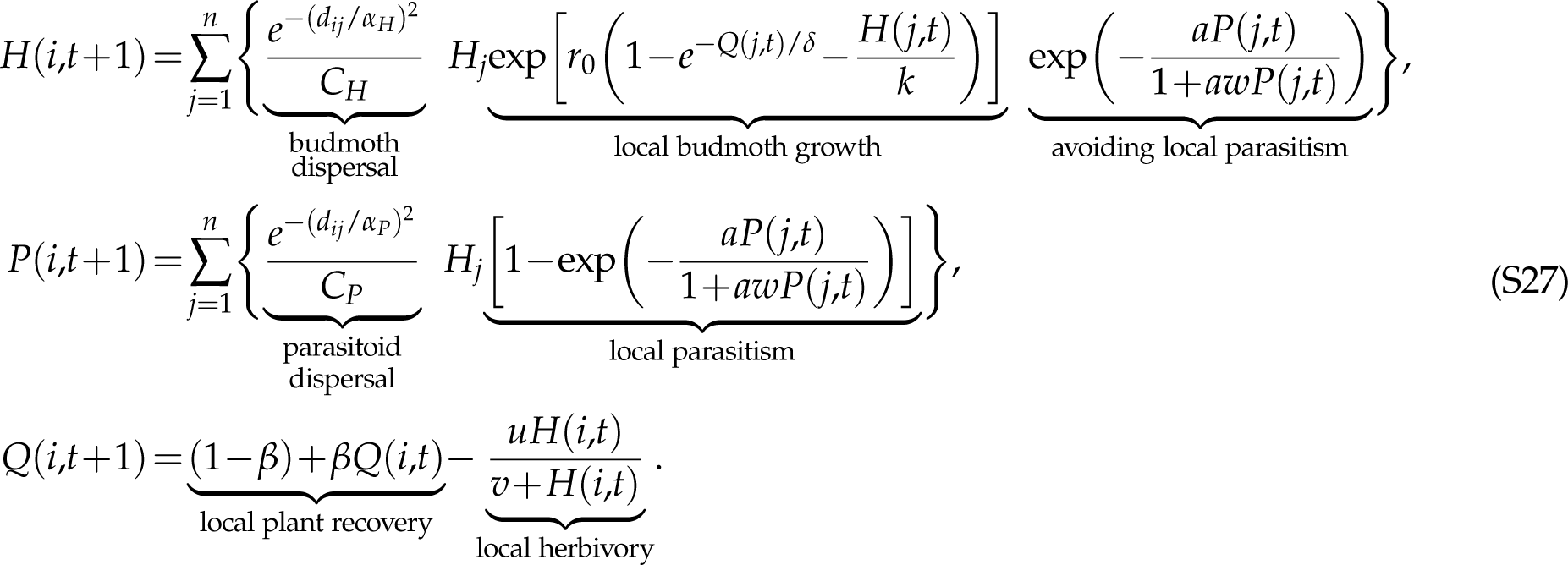

For dispersal, *d_ij_* is the distance between patches, and we assume a Gaussian kernel with dispersal parameters *α_H_*and *α_P_* for the budmoths and parasitoids; *C_H_*and *C_P_* are normalization constants. Before dispersal, we assume that the local budmoth and parasitoid densities change in accordance to the local dynamics. For the budmoth, *r*_0_ is the maximum growth rate^2^, *δ* is a scale parameter that determines how fast the growth rate approaches *r*_0_ with increasing plant quality *Q*(*j*,*t*), and *k* is the budmoth carrying capacity in the limit of large *Q*(*j*,*t*), so 1/*k* characterizes intraspecific competition. Local parasitism is described by a modified Nicholson-Bailey framework: the exponential describes the probability of a budmoth avoiding parasitism, and is parametrized by *a* and *w* representing the search efficiency of a parasitoid and the mutual interference between parasitoids. Finally, for local plant dynamics, *β* represent the rate at which plant quality *Q*(*i*,*t*) recovers towards 1, while *u* and *v* characterize the impact of budmoth herbivory on plant quality. We note that Johnson et al. (2004) also introduced an additional parameter that is meant to approximate the effects of demographic stochasticity, although it was omitted in Johnson et al. (2006); we chose to omit it as well.

Most parameter values can be found in Table 1 of Johnson et al. (2006), although note that the parameter labels (*r*_0_,*K*,*A*,*W*,*A*,*C*,*D*,*δ*) should be corrected to (*r*_0_,*k*,*a*,*w*,*β*,*u*,*v*,*δ*). Other parameter values that can only be found in the main text are: *α_H_* = 10 km and *α_P_* = 5 km. For the normalization constants *C_H_* and *C_P_*, the authors stated that they were chosen such that the “total proportion of dispersal across suitable and unsuitable habitat sums to one”. Therefore, we discretized the landscape into an arbitrarily large spatial grid of resolution 3*×*3 km (based on the patch dimensions in Johnson et al. (2004)), and assumed that the Gaussian kernel applied to any pair of grid cells, and not just grid cells assigned as suitable patches. We then obtained *C_H_* using

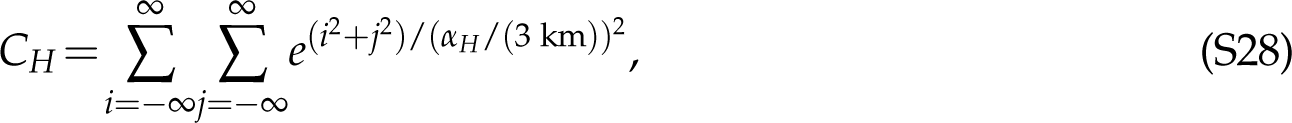

where *i* and *j* here are grid indices (not patch indices). A similar expression was used for *C_P_*.

We wanted to replicate the scenario in Johnson et al. (2004, 2006) where patches near the center of the landscape had the highest connectivity. According to Johnson et al. (2004), “habitat configurations were created by assuming that the probability of a patch being suitable declined exponentially with the distance from the focal location”. Therefore, we drew random samples from an exponential distribution with a mean of 5 grid units, applied a random sign, and rounded them to the nearest integer. Pairs of these integers were then used as grid indices for the suitable patches. We generated 500 unique patches this way.

Since we were only interested in the deterministic version of the model, we did not introduce random variations into *r*_0_ for each patch and timestep as was done in Johnson et al. (2006). Also, even though we initialized the simulation the same way as Johnson et al. (2006), we ran the simulation for many time steps before the start of the time horizon, to allow any transients to die off.

### S6.2 Larch budmoth: Objective function and adjoint equations

A possible objective function is to maximize the plant quality over a time horizon from *t* = 1 to *T*, with weight *W*(*i*,*t*) assigned to patch *i* at time *t*, so

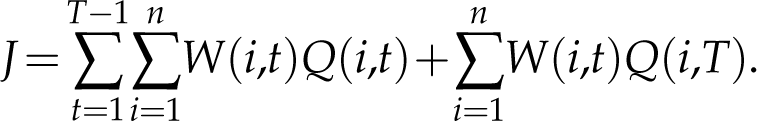

We choose an arbitrary time horizon of *T* = 200 years, and we assigned equal weight to all patches, but more weight to more recent years, by having

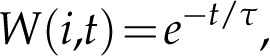

where *τ* = 50 years. Just as in the pine looper example, the decaying weights reduce the dependence of the time-dependent sensitivities on the time horizon, should the dynamics be quasiperiodic.

The Hamiltonian (which we denote by *H* to avoid confusion with the budmoth density) is given by

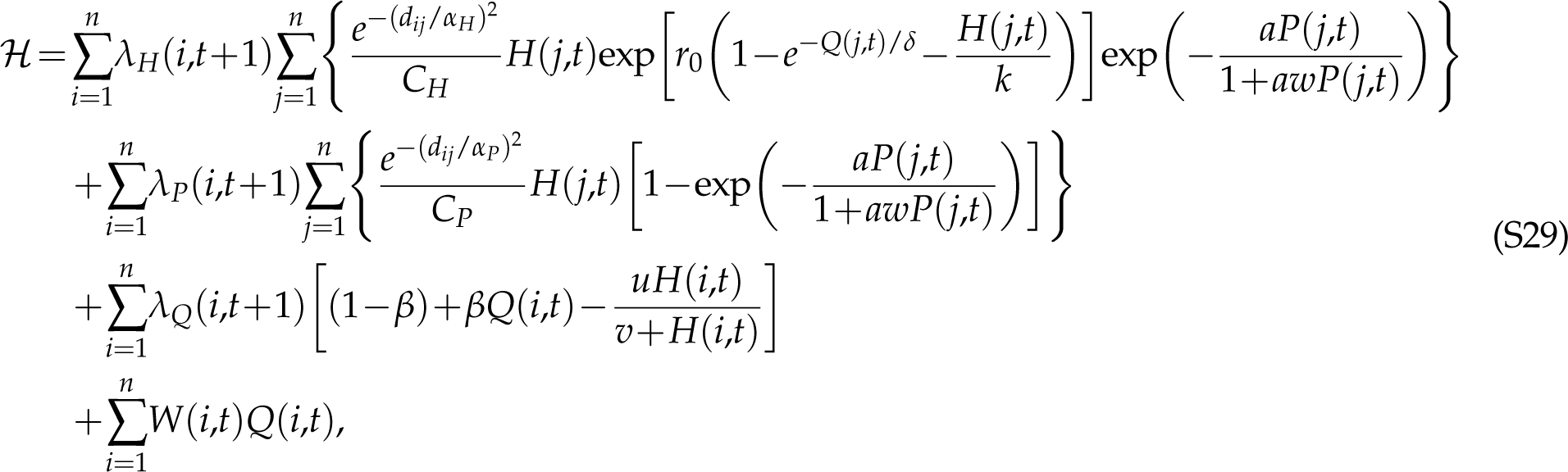

where the last term comes from the objective function. The adjoint equations are then given by

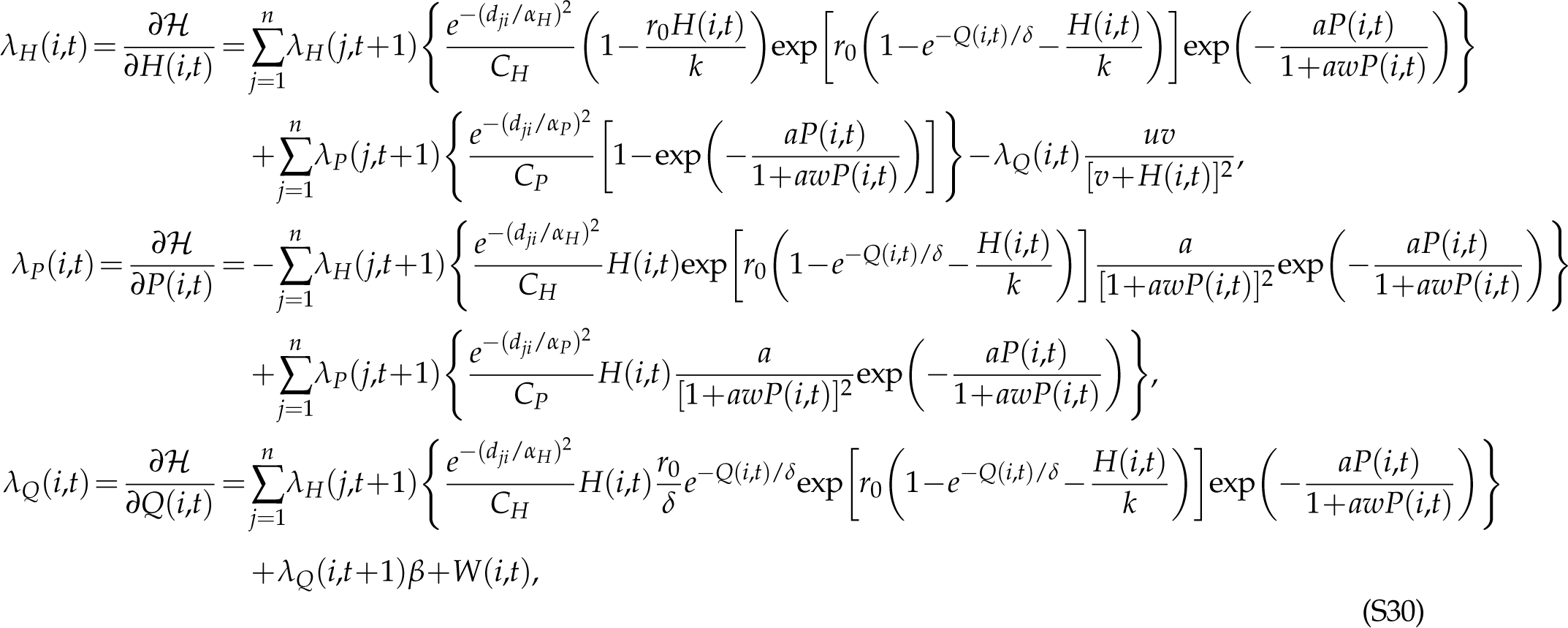

with terminal conditions

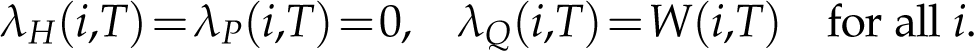

## S7 Supplementary figures and tables from Example 1: Exogenous disease spillover in multi-species sink networks

## S8 Supplementary figures and tables from Example 2: Leopard frogs as reservoirs of the amphibian chytrid fungus

## S9 Supplementary figures and tables from Example 3: Population cycles in the pine looper and the larch budmoth

### S9.1 Pine looper

### S9.2 Larch budmoth

**Figure S1:**
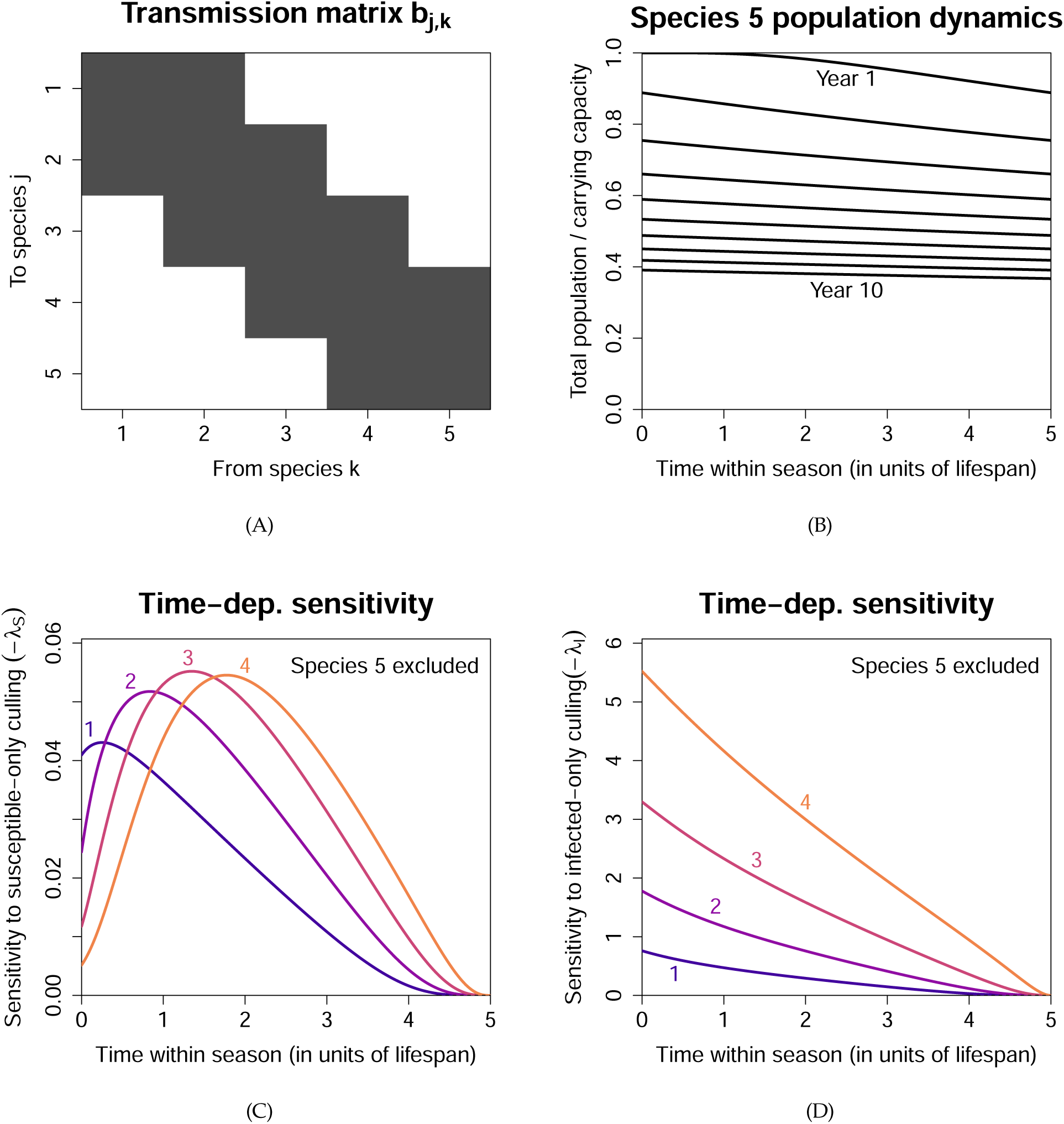
Additional figures from Network 1. **(A)** Matrix representation of the transmission coefficients *b_j_*_,*k*_. **(B)** Population decline in the species of concern (species 5) over a 10-year period, assuming that the population size at the end of one season carries over to the start of the next season. The purpose is to show that the population decline can be significant despite the low infection prevalence shown in Fig. 3(D). **(C)** Time-dependent sensitivity when only susceptible individuals are culled. **(D)** Time-dependent sensitivity when only infected individuals are culled (*−λ_Ij_*). The weighted sum of (C) and (D) gives the time-dependent sensitivity to indiscriminate culling (*−λ_Nj_*) shown in Fig. 3(G).

**Figure S2:**
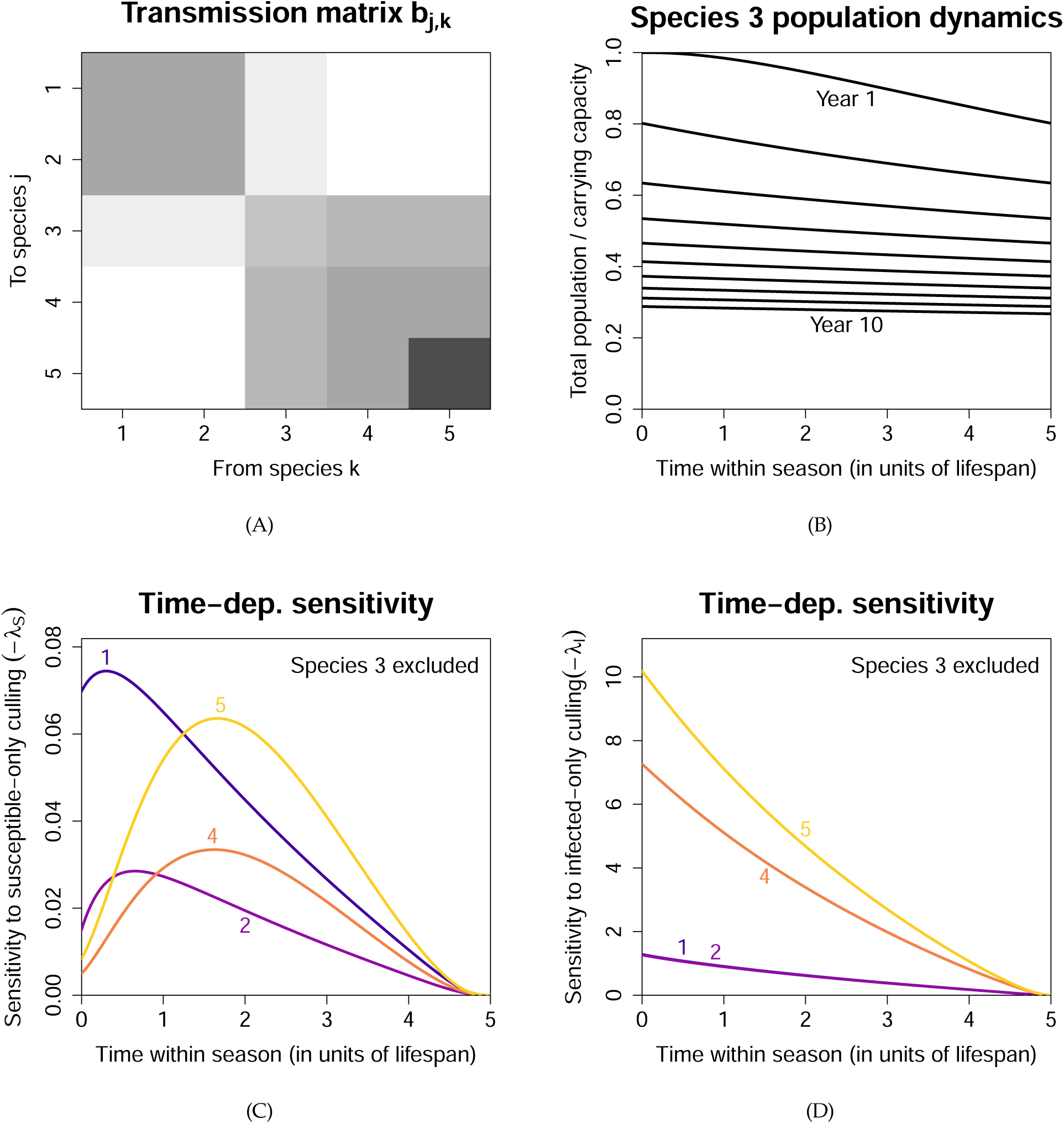
Similar to Fig. S1, except for Network 2.

**Figure S3:**
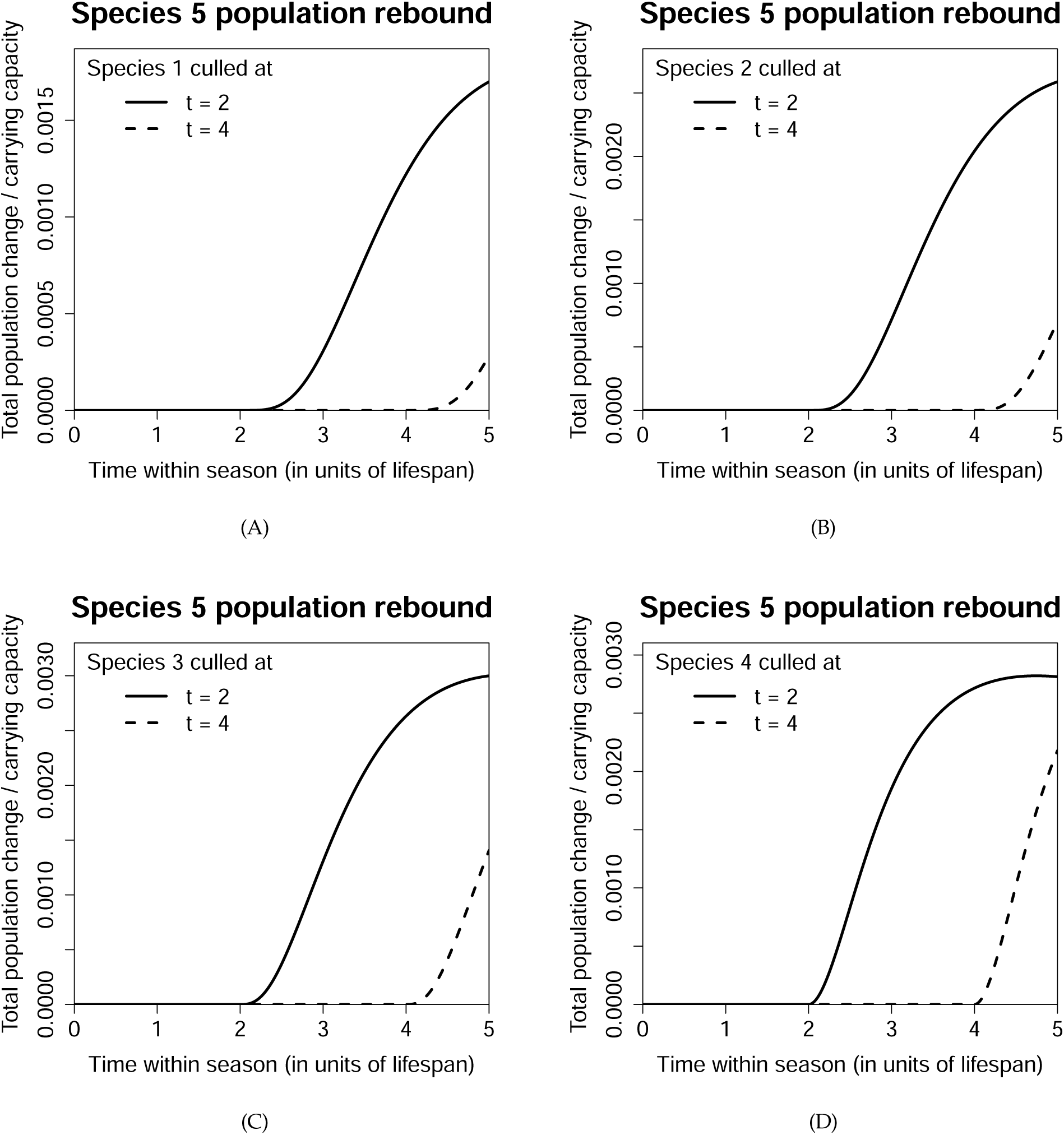
For Network 1, the graphs above show the population rebound in the species of concern (species 5) when 10% of another species is indiscriminately culled. Late culling leaves less time for the population to rebound (affecting the terminal payoffs *V_SjC_* and *V_IjC_*), and also less time for the rebound to contribute to the integral in the reward function.

**Figure S4:**
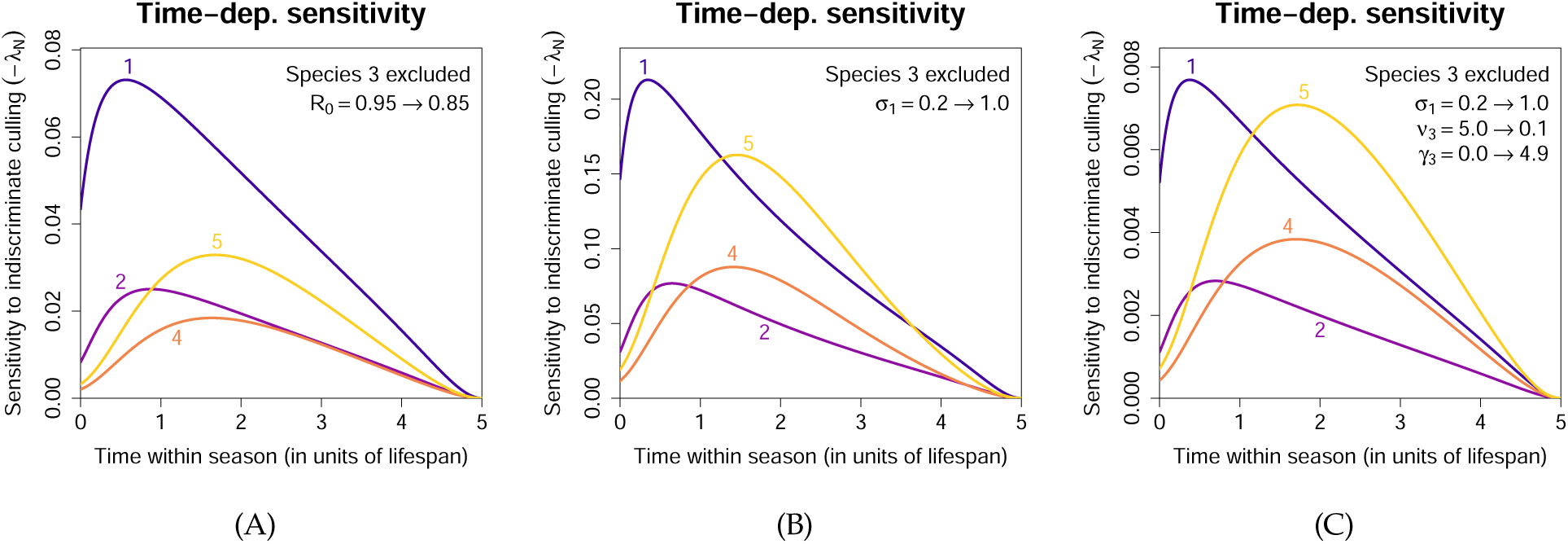
More results from Network 2, obtained using modified parameter values. **(A)** Reducing *R*_0_ caused the importance of species 5 to fall entirely below species 1, due to multi-step within-module transmission becoming less likely at a lower *R*_0_. **(B)** Increasing the exogenous spillover rate *σ*_1_ caused the most important species to switch from species 5 back to species 1 towards the end of the season. This is due to the large decrease in the population of species 3 resulting from the increased spillover; the switch no longer occurred in **(C)** when we converted most of the disease-induced mortality rate in species 3 to its recovery rate.

**Figure S5:**
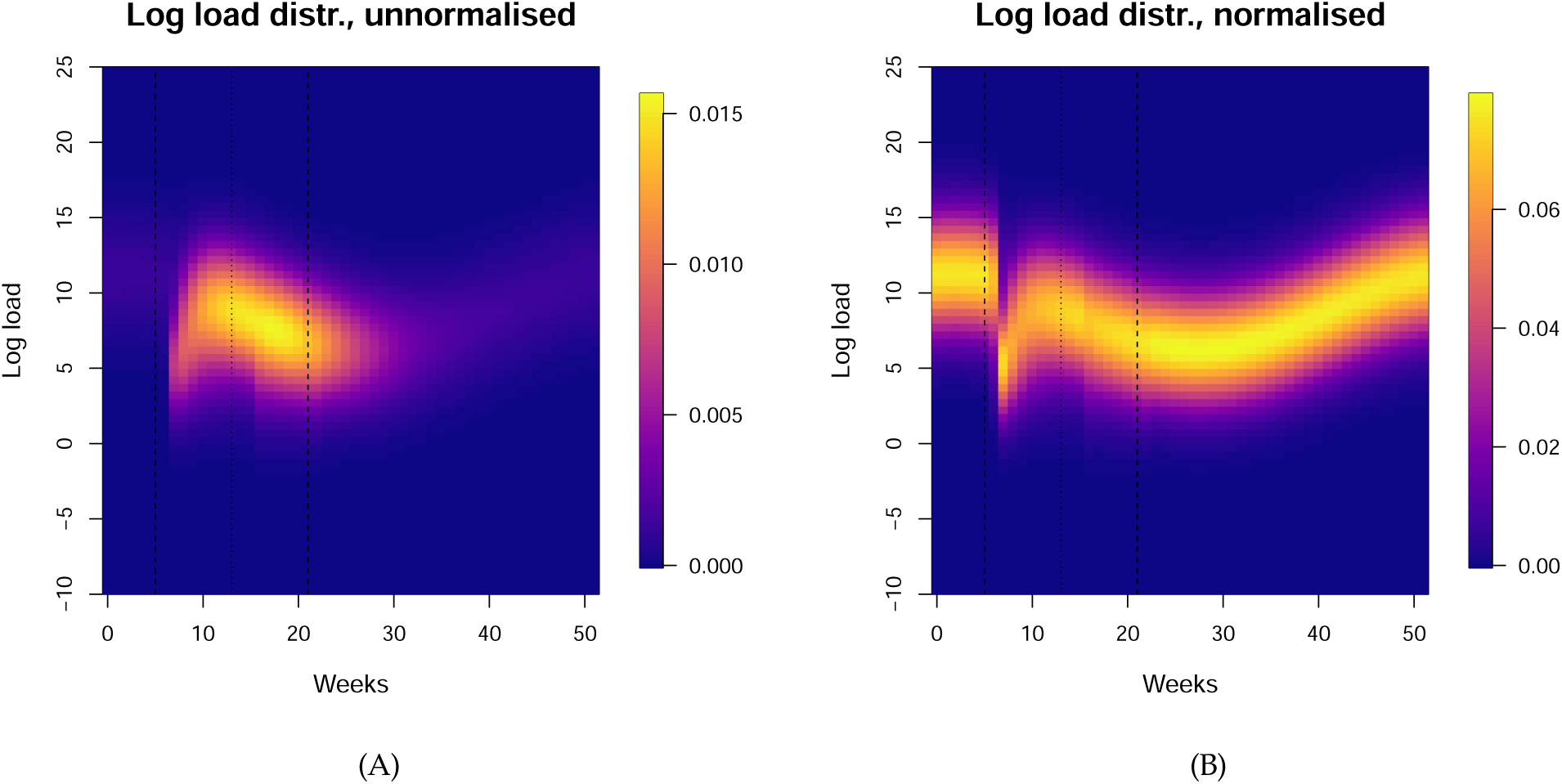
**(A)** Number of infected frogs in each log load bin, each week across the year, at steady state. **(B)** Log load distribution each week, obtained by normalizing the sum of each vertical column in (A) to 1. Due to the temperature-dependent load dynamics, we see that the load is the lowest in summer and the highest in winter.

**Figure S6:**
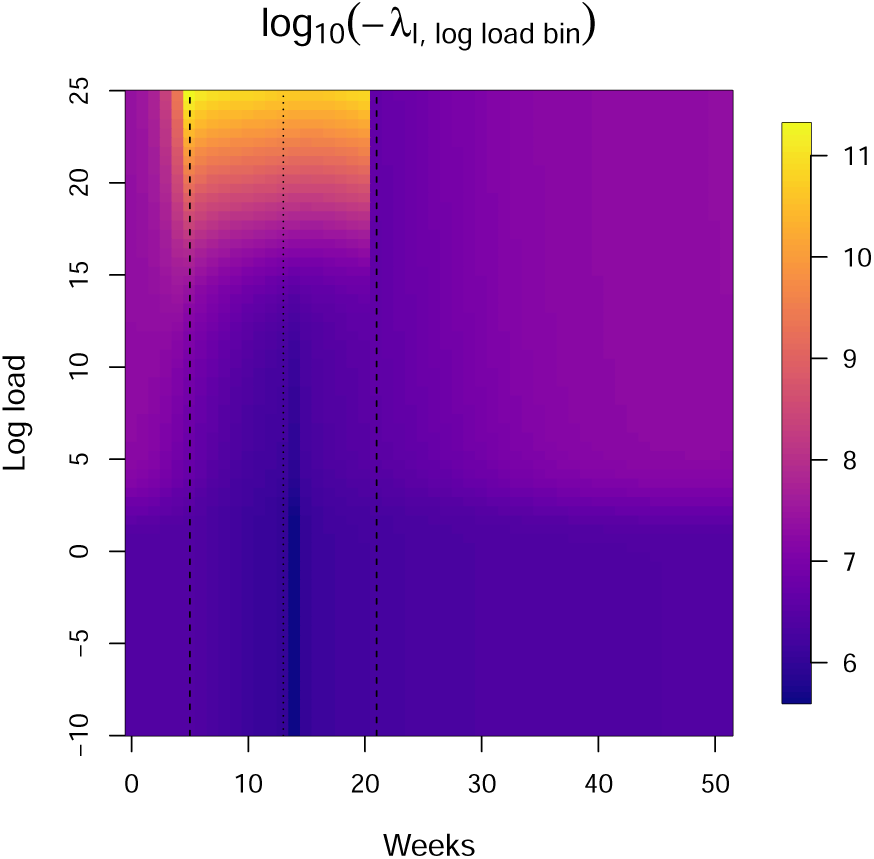
The sensitivity to removing an infected frog from each log load bin, each week across the year. Note that this sensitivity does not take into account whether the log load bin is actually “occupied” which is why we choose to work with *−λ_I_*(*t*) as defined in Eqn. (24) instead.

**Figure S7:**
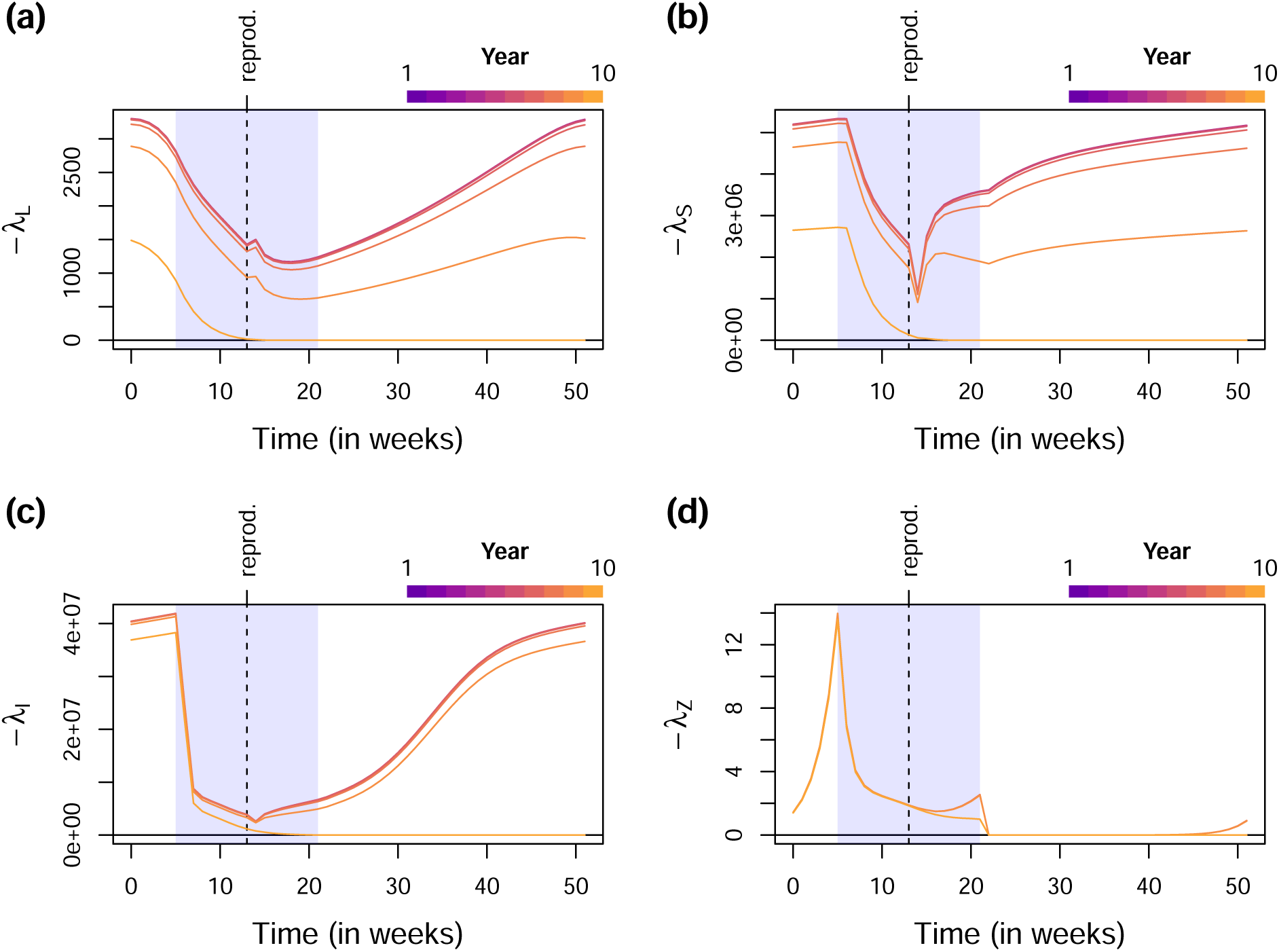
Effects of the time horizon *T*. Similar to Fig. 6, except that we have also shown the sensitivities every year within the time horizon. We see that if the time horizon is sufficiently long, the seasonal sensitivity patterns during the first few years are identical. At steady state, each year starts with the same “initial conditions”, so the second year can be thought of as the same system with a time horizon of 9 years, the third year a time horizon of 8 years, etc. Therefore, the fact that the early years show identical seasonal patterns means that the early-year patterns are independent of the time horizon, and hence expected to be the same as when the time horizon is infinite.

**Figure S8:**
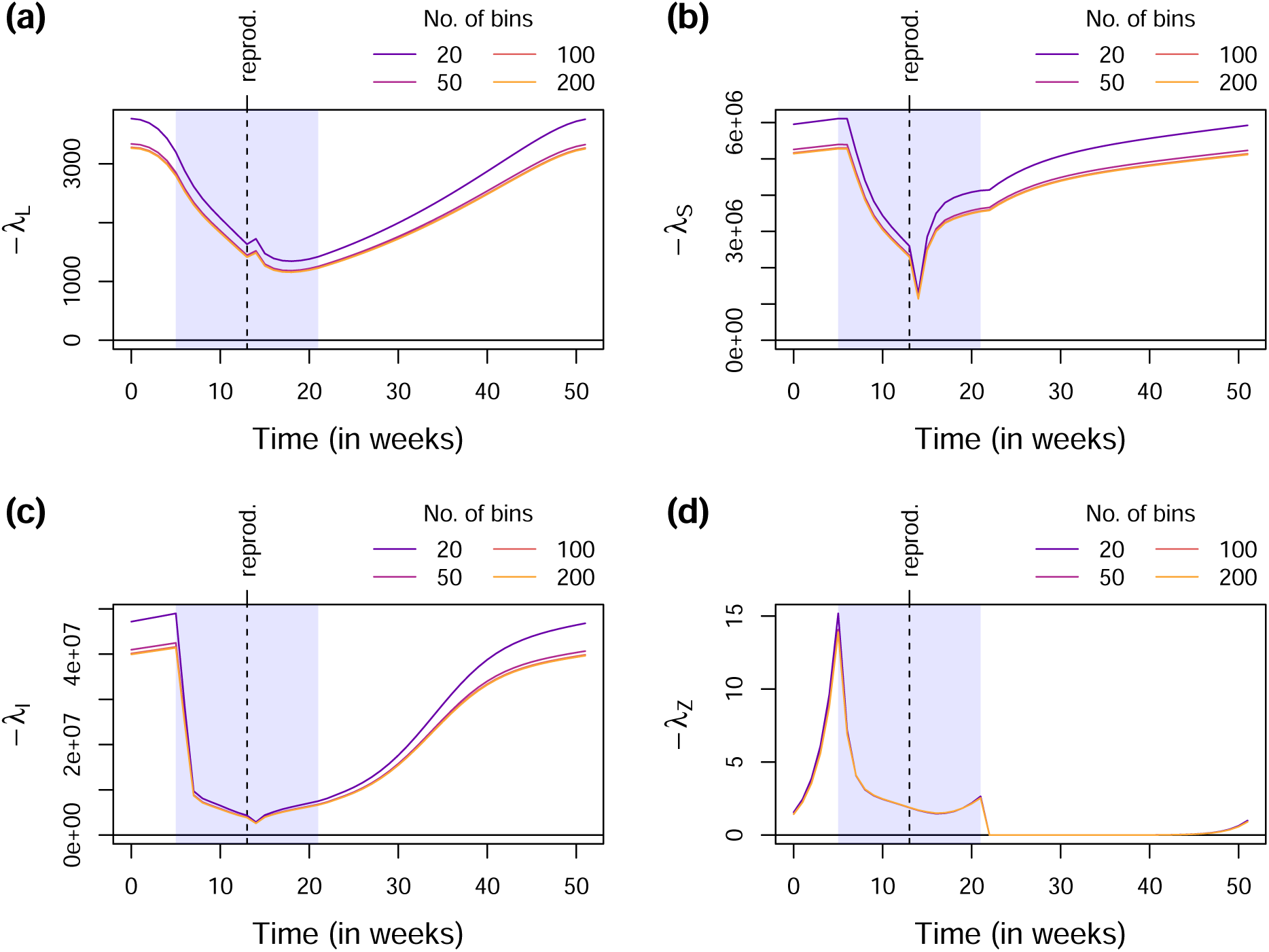
Varying the number of bins in the discretized IPM. Similar to Fig. 6, except that we have varied the number of bins used when discretizing the IPM.

**Figure S9:**
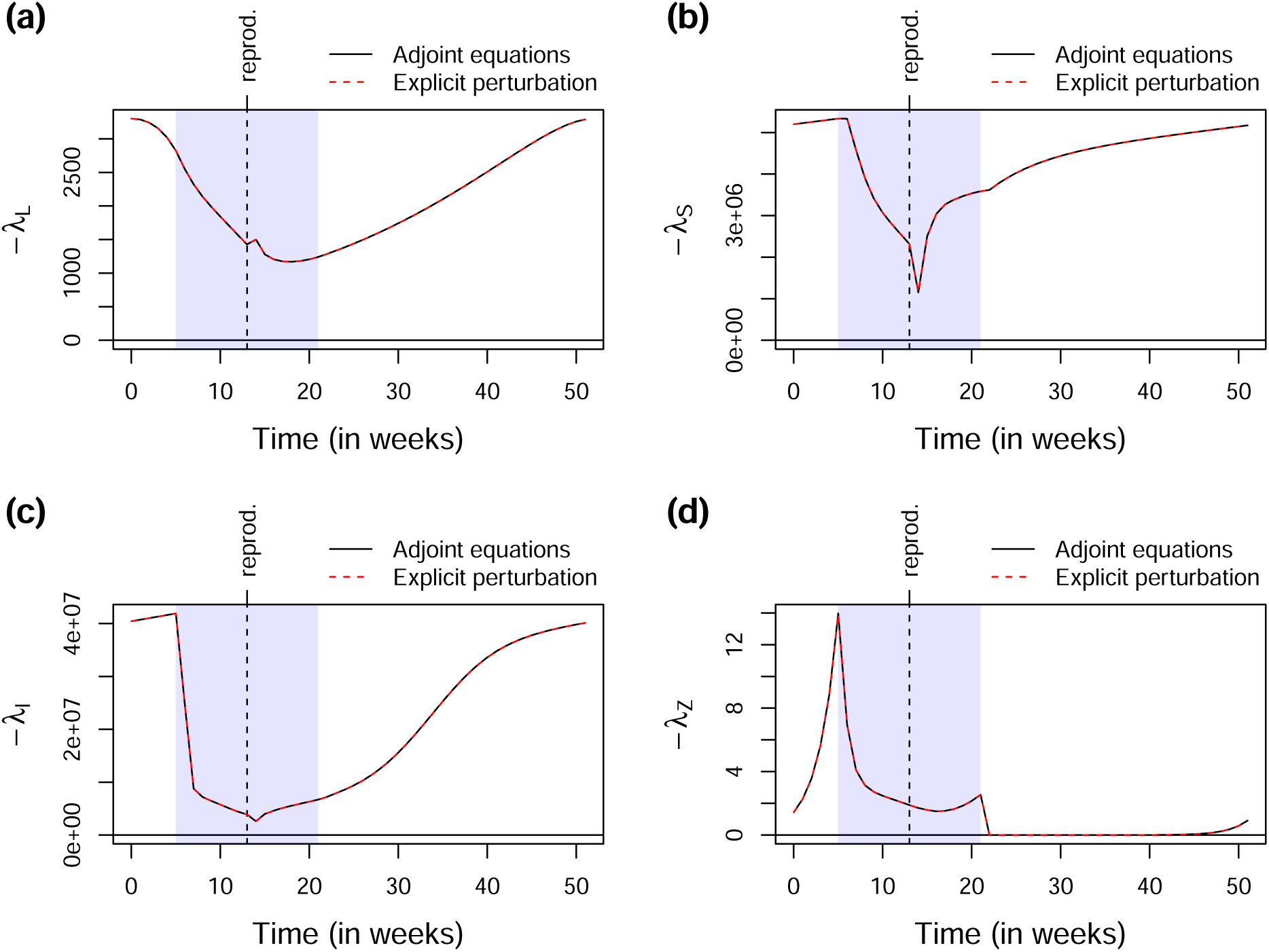
Checking against explicit perturbations. Similar to Fig. 6, except that we have also shown the sensitivities obtained by explicitly perturbing the state variables at each time point (red dashed lines). The perfect agreement with the adjoint variables implies that the adjoint equations have been correctly derived and implemented.

**Table S1:** Parameter values of the maternal effects model, fitted separately using data at three sites.

| Site | $r$ | $s$ | $u$ | $x_{\min}$ | $\beta$ |
| --- | --- | --- | --- | --- | --- |
| Culbin | $5.064 \times 10^{-5}$ | 0.079 | 3.364 | 2.150 | 0.204 |
| Roseisle | $5.760 \times 10^{-2}$ | 0.246 | 3.644 | 0.510 | 1.016 |
| Tentsmuir | $5.677 \times 10^{-3}$ | 0.000 | 4.075 | 0.618 | 0.294 |

**Figure S10:**
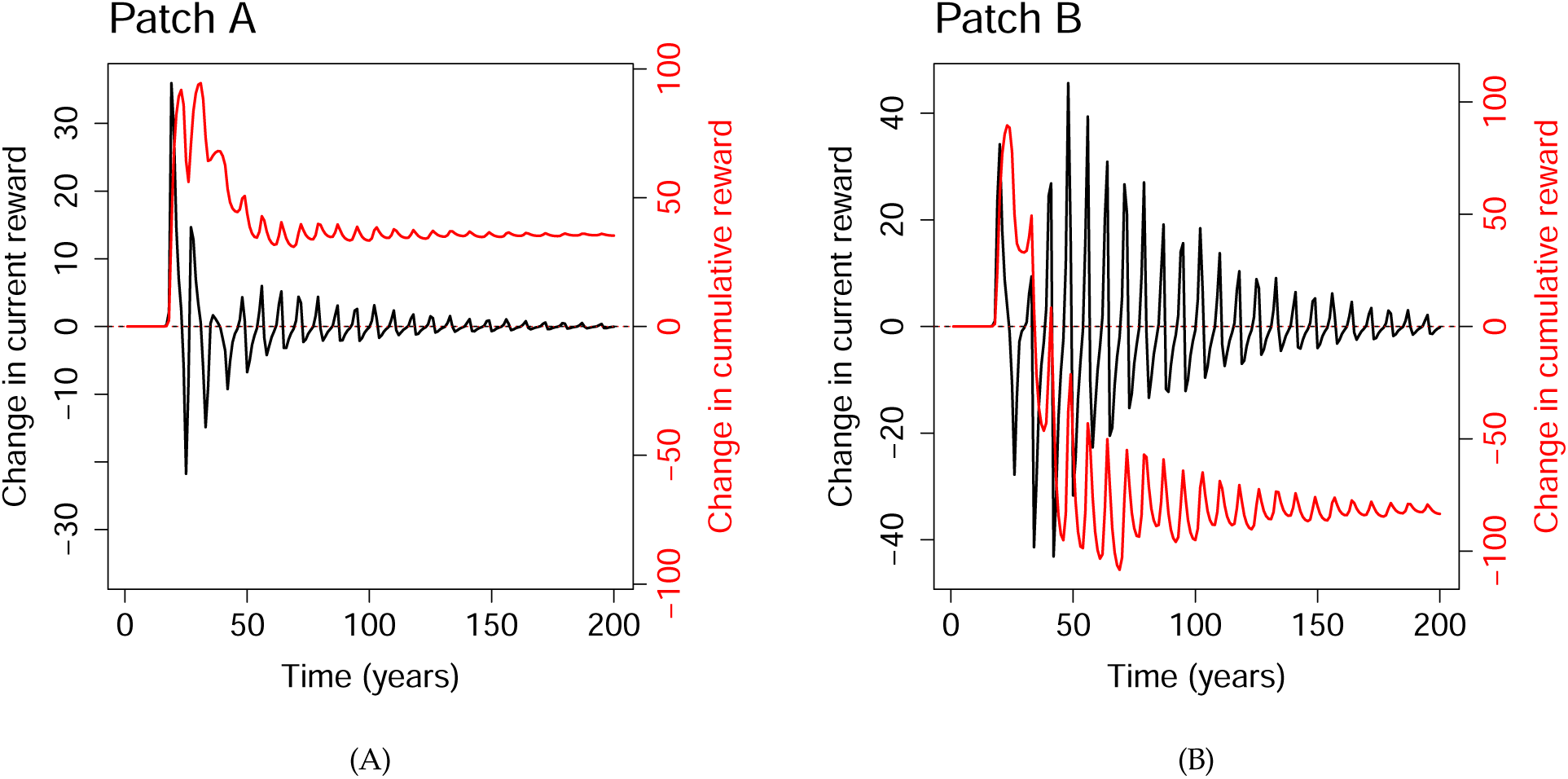
Phase plane diagram at Roseisle, Tentsmuir and Culbin, showing the periodic steady-state solution at Roseisle, and the quasiperiodic steady-state solutions at Tentsmuir and Culbin. At Roseisle, we only showed 10 years to illustrate one complete cycle of two oscillations, whereas at Tentsmuir and Culbin, we showed every year across the time horizon of 200 years.

**Figure S11:**
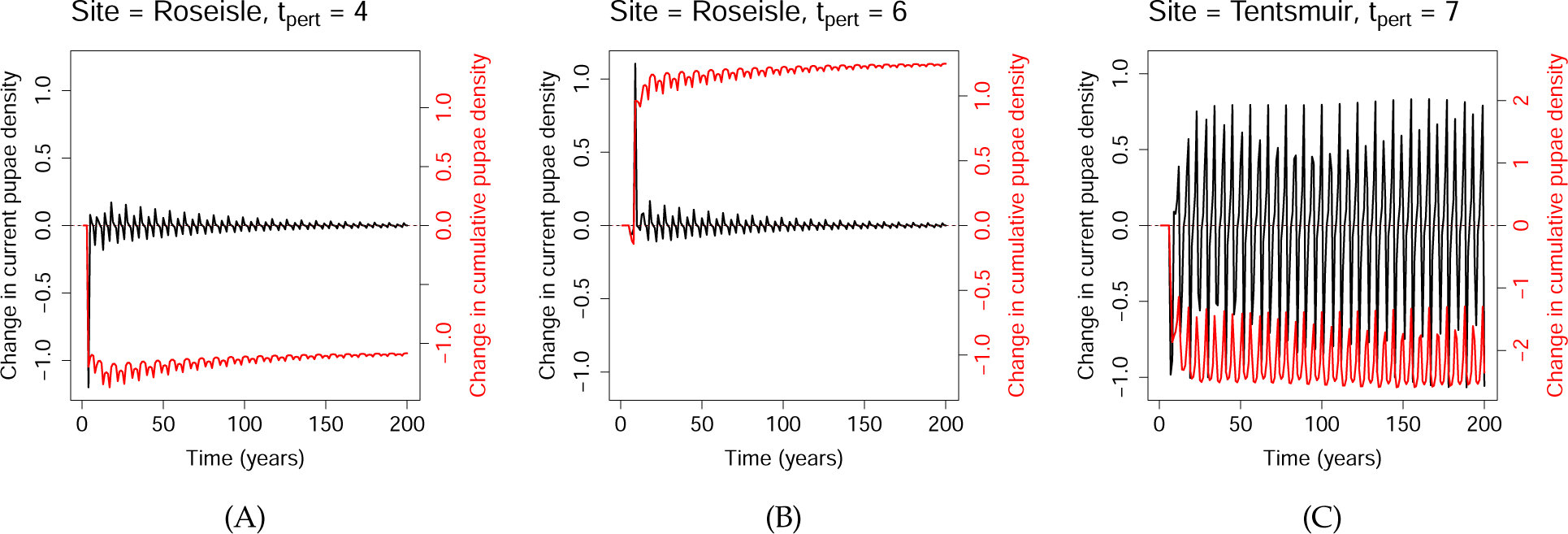
Changes in the current pupae density *N*(*t*) and the cumulative moth density ∑*^t^ N*(*t^′^*) at all *t*, following a 20% cull at *t* = *t*_pert_. **(A)** Roseisle; *t*_pert_ = 4. **(B)** Roseisle; *t*_pert_ = 6. **(C)** Tentsmuir; *t*_pert_ = 7. We see that the changes in current density decay with time in (A) and (B), but persist indefinitely in (C), likely because of the steady-state trajectories being periodic in Roseisle, but quasiperiodic in Tentsmuir. As a result, the cumulative changes approach constant, non-oscillatory values in (A) and (B), but remain oscillatory in (C). Note that the choices of *t*_pert_ are unimportant here; we made these specific choices only to facilitate comparison with Fig. 7(D-F) and Fig. S12.

**Figure S12:**
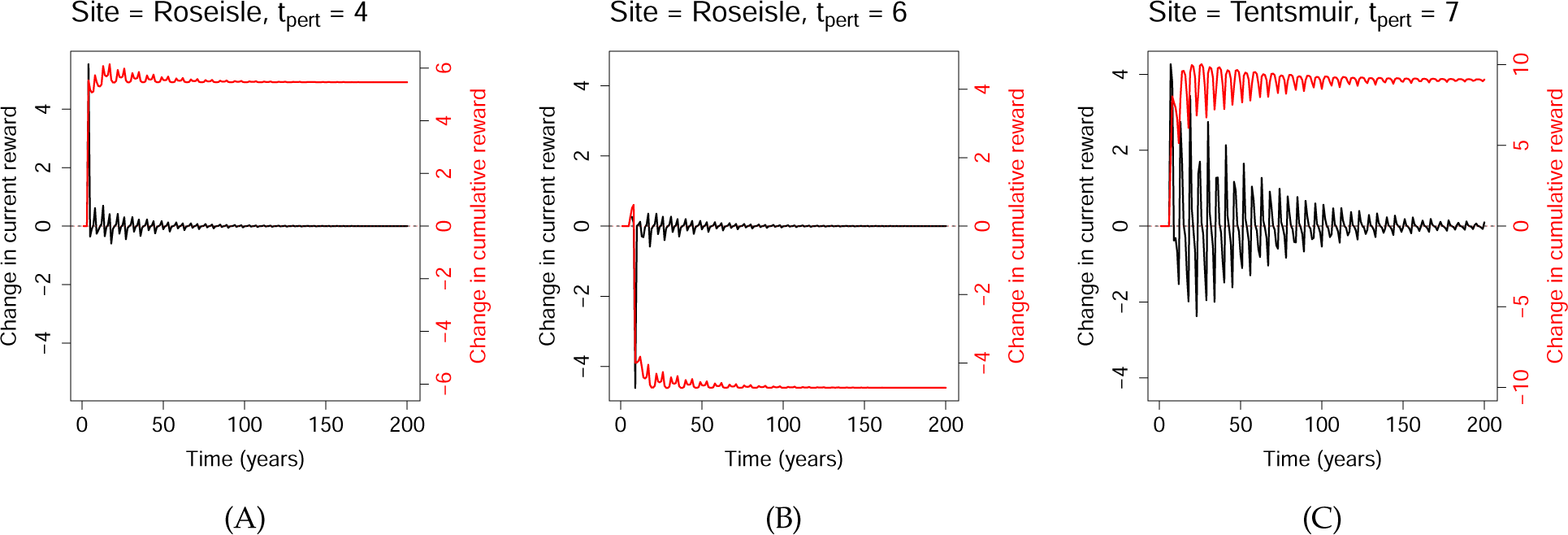
Changes in the current reward *−N*(*t*)*W*(*t*) and the cumulative reward *−*∑*^t^ N*(*t^′^*)*W*(*t^′^*) at all *t*, following a 20% cull at *t* = *t*_pert_. We have rescaled these changes by a factor of 1/0.2, so that the cumulative reward at *t* = *T* = 200 should be approximately equal to the demi-elasticity in Fig. S13 at *t* = *t*_pert_; any small discrepancies are due to nonlinearities from the relatively large perturbation. **(A)** Roseisle; *t*_pert_ = 4. **(B)** Roseisle; *t*_pert_ = 6. **(C)** Tentsmuir; *t*_pert_ = 7. Note that unlike Fig. S11(C), the changes in current reward decay in time because of the decaying weight *W*(*t*). This allows the cumulative reward to approach a constant, non-oscillatory value, and hence reduces the dependence of the demi-elasticities on the time horizon *T*.

**Figure S13:**
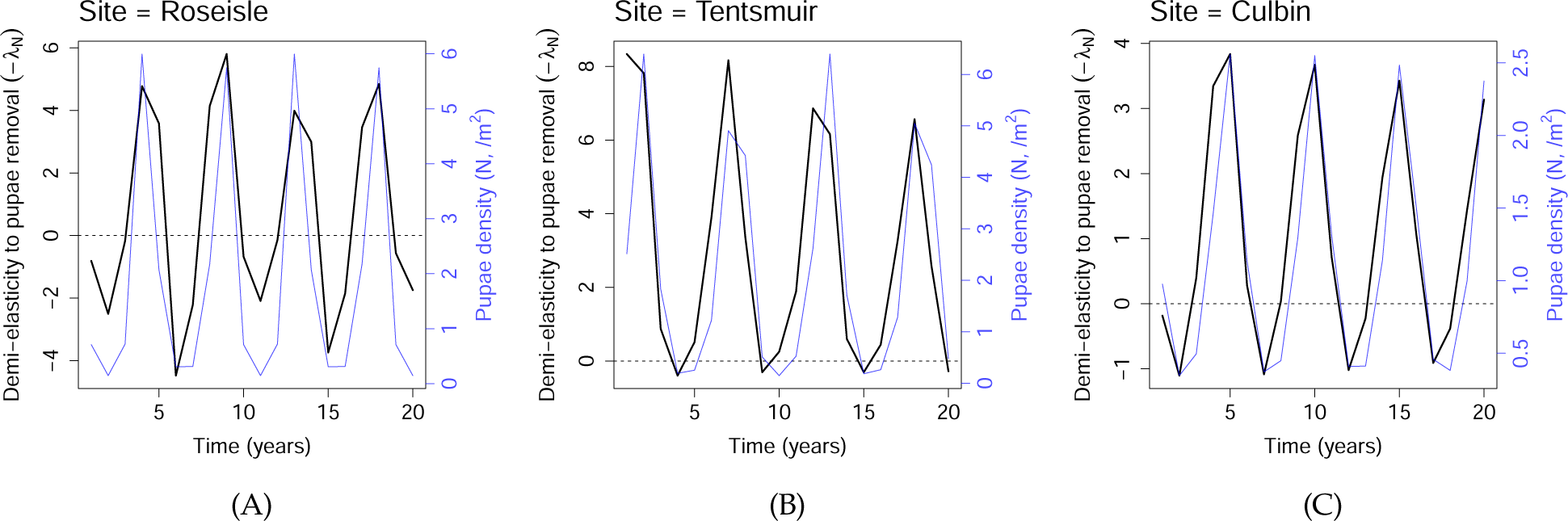
Demi-elasticities of the reward to the culling of pine looper at **(A)** Roseisle, **(B)** Tentsmuir and **(C)** Culbin.

**Figure S14:**
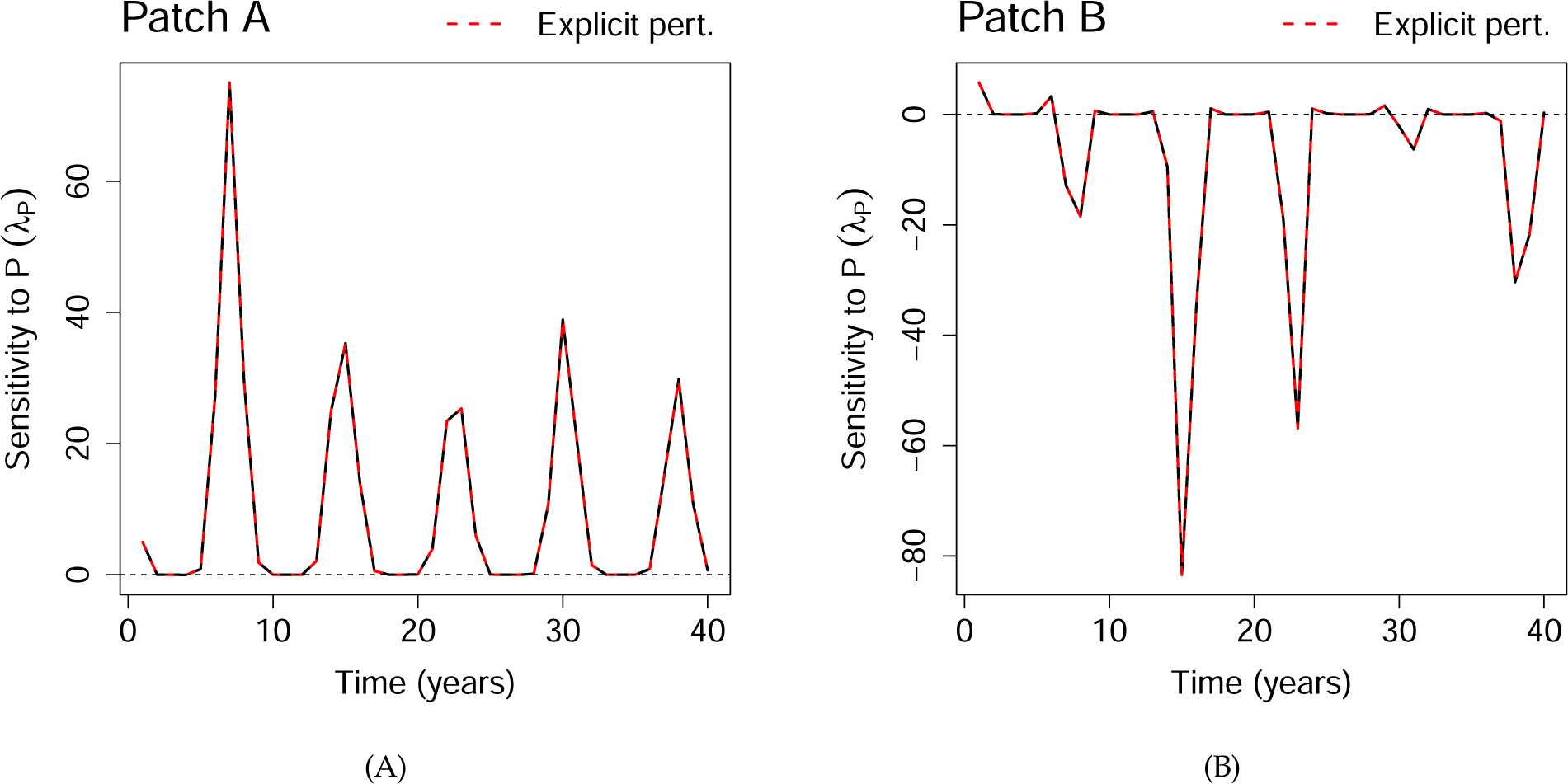
Verifying that TDSA gives the correct sensitivities for the larch budmoth model using explicit perturbations. We focused on the two patches discussed in Fig. 8.

**Figure S15:**
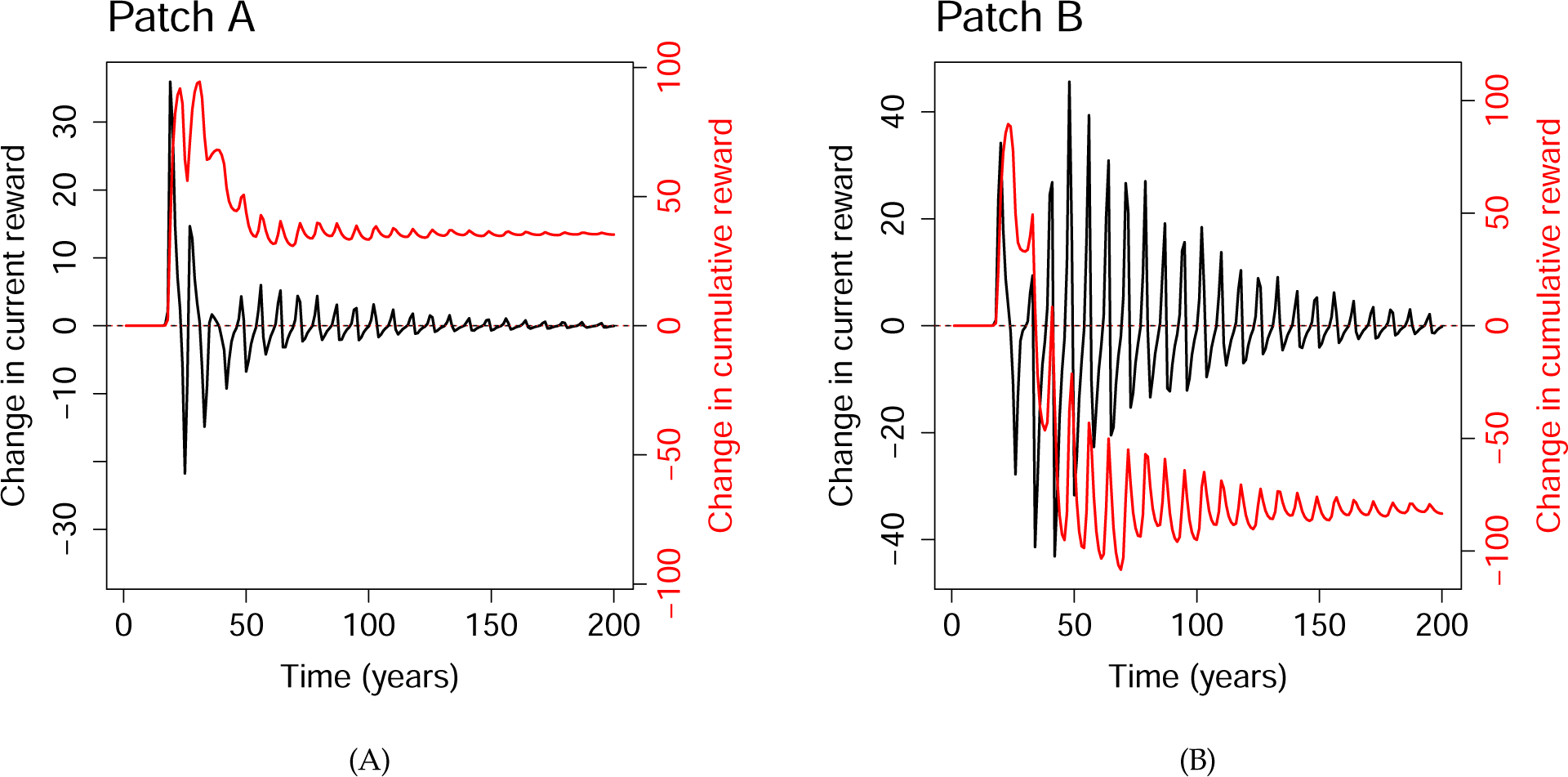
The effects of adding parasitoids at *t* = 15 to the two patches discussed in Fig. 8. The current reward refers to the sum of plant quality times the weight in the current year, and the cumulative reward the sum of current rewards from *t* = 1 up to the current year. We used small perturbations to ensure linearity, but scaled the results by the inverse of the perturbation size, so that the change in cumulative reward at *t* = *T* = 200 (the end of the time horizon) should be equal to the sensitivity at *t* = 15 (the time of perturbation). As expected, they indeed agree with Fig. S14 at *t* = 15 (*∼* 40 for Patch A, *∼−*80 for Patch B).

## Footnotes

1 We chose *demi-elasticity*, because the more obvious choice of *semi-elasticity* is often used in economics to represent the fractional change in objective given an absolute change in the perturbed variable, exactly the opposite of demi-elasticity. “Demi” is also a useful mnemonic for “denominator”. One author’s suggestion of *sensi-lasticity* went unheeded.

2 Or nearly so, since *Q*(*j*,*t*) cannot exceed 1, so the maximum growth rate is really *r*_0_(1*−e^−^*^1/*δ*^) *≃* 0.989*r*_0_ for the chosen value of *δ* = 0.22.

